# Imaging cellular activity simultaneously across all organs of a vertebrate reveals body-wide circuits

**DOI:** 10.1101/2025.08.20.670374

**Authors:** Virginia M. S. Ruetten, Wei Zheng, Igor Siwanowicz, Brett D. Mensh, Mark Eddison, Amy Hu, Yunfeng Chi, Andrew L. Lemire, Caiying Guo, Mykola Kadobianskyi, Marc Renz, Sara Lelek-Greskovic, Yisheng He, Kari Close, Gudrun Ihrke, Mariela D. Petkova, Michael Cook, Christopher J. Knecht, Aparna Dev, Alyson Petruncio, Yinan Wan, Jeff W. Lichtman, Florian Engert, Mark C. Fishman, Benjamin Judkewitz, Mikail Rubinov, Philipp J. Keller, Chie Satou, Guoqiang Yu, Paul W. Tillberg, Maneesh Sahani, Misha B. Ahrens

## Abstract

All cells in an animal collectively ensure, moment-to-moment, the survival of the whole organism in the face of environmental stressors^1,2^. Physiology seeks to elucidate the intricate network of interactions that sustain life, which often span multiple organs, cell types, and timescales, but a major challenge lies in the inability to simultaneously record time-varying cellular activity throughout the entire body.

We developed WHOLISTIC (WHole Organism Live Imaging System for recording Tissue and IntraCellular activity), a method to image second-timescale, time-varying intracellular dynamics across cell-types of the vertebrate body. By advancing and integrating volumetric fluorescence microscopy, machine learning, and pancellular transgenic expression of calcium sensors in transparent young *Danio rerio* (zebrafish) and with proof of concept in adult *Danionella*, the method enables real-time recording of cellular dynamics across the organism. Calcium is a universal intracellular messenger, with a large array of cellular processes depending on changes in calcium concentration across varying time-scales, making it an ideal proxy of cellular activity^3^.

Using this platform to screen the dynamics of most cells in the body, we discovered unexpected responses of specific cell types to stimuli, such as chondrocyte reactions to cold, meningeal responses to ketamine, and state-dependent activity, such as oscillatory ependymal-cell activity during periods of extended motor quiescence. At the organ scale, the method uncovered pulsating traveling waves along the kidney nephron. At the multi-organ scale, we uncovered muscle synergies and independencies, as well as muscle-organ interactions. Integration with optogenetics allowed us to all-optically determine the causal direction of brain-body interactions. At the whole-organism scale, the method captured the rapid brainstem-controlled redistribution of blood flow across the body.

Finally, we advanced Whole-Body Expansion Microscopy^4^ to provide ground-truth molecular and ultrastructural anatomical context, explaining the spatiotemporal structure of activity captured by WHOLISTIC. Together, these innovations establish a new paradigm for systems biology, bridging cellular and organismal physiology, with broad implications for both fundamental research and drug discovery.

## Main

Cells across an organism must continuously coordinate with one another to sustain life and adapt to changing external environments and alterations within the body. This is achieved through a complex network of dynamic interactions that maintains homeostasis and underpins the organism’s resilience to stress^5^. Disease can arise from breakdowns in these intercellular feedback mechanisms^6,7,8,9,10,11^. Although biomedicine has made strides in uncovering key interactions, such as stress responses mediated by neuroendocrine signaling^12^ or neural pathways between the brain and the gut^13,14,15,16^, a vast number of mechanisms of whole-organism function remain elusive^1,17,2^. Our understanding is limited by the challenges of accessing body-wide cellular dynamics in both health and disease, highlighting the need for advancements in technologies to measure, analyze, and model body-wide control mechanisms at the cellular level.

Modern synergies between imaging technology and protein engineering allow for time-varying molecular signals to be recorded in tissues. This has caused revolutions in fields like neuroscience through the imaging of calcium — a fast, universal intracellular messenger involved in a wide range of cellular processes, including neuronal action potentials^3^ — and other signals including voltage and neuromodulators across many neurons simultaneously^18,19,20,21,22^. However, time-varying activity patterns of most cell types in the body have not yet been recorded.

For most vertebrate models, optical access to large, opaque tissues poses a currently insurmountable challenge to whole-body imaging. Transparent vertebrate animals such as young zebrafish and adult *Danionella* ^23,24,25,26,27,28^ overcome this barrier, making them uniquely suited as models for in vivo studies of cellular dynamics across the body^29^, offering unparalleled access to the inner workings of evolutionarily conserved organs such as the liver, pancreas, gut, brain, and the immune and cardiovascular systems^30,31^.

This study introduces WHOLISTIC (WHole Organism Live Imaging System for recording Tissue and IntraCellular activity), a platform that enables in vivo imaging of cellular calcium dynamics, generalizable to other molecular dynamics, across nearly all cells of transparent vertebrates, such as the young zebrafish. By extending and integrating an array of technical advances, including pancellular transgenic lines expressing genetically encoded calcium indicators in almost all cells in the body, high-speed volumetric fluorescence imaging, and a suite of computational methods for registration and cell population analysis, along with Whole-Body Expansion Microscopy, we capture, analyze, and interpret cellular activity across tissues and organ systems.

Together, these techniques provide a discovery platform that allows for the comprehensive investigation of coordinated cellular activity within an intact and awake vertebrate.

### WHOLISTIC captures cellular calcium activity across organ systems

To image dynamic fluctuations of intracellular calcium across cell types of the body of young zebrafish (Fig. 1a), we engineered a pancellular transgenic zebrafish line that expresses the genetically encoded calcium indicator GCaMP7f under the control of the ubiquitin promoter^32,33^, *Tg(ubi:tTA; TRE:GCaMP7f)* (Fig. 1b,c). To avoid transient or sparse expression^34,35,36^, we employed the binary expression system tTA / TRE to enhance the concentration of GCaMP7f^37^, which resulted in extensive and sustained pancellular expression with no observed pathological manifestations (Fig. 1b, Extended Data Fig. 1a,b). This approach can be used with a range of available molecular sensors that report dynamic properties other than calcium concentration^38^, such as transmembrane voltage^39^, membrane tension^40^, and intra- or extracellular concentrations of diverse molecules^41,42^. Here, we chose to focus first on intracellular calcium due to its broad involvement in numerous cellular processes across all cell types^3^.

**Fig. 1:**
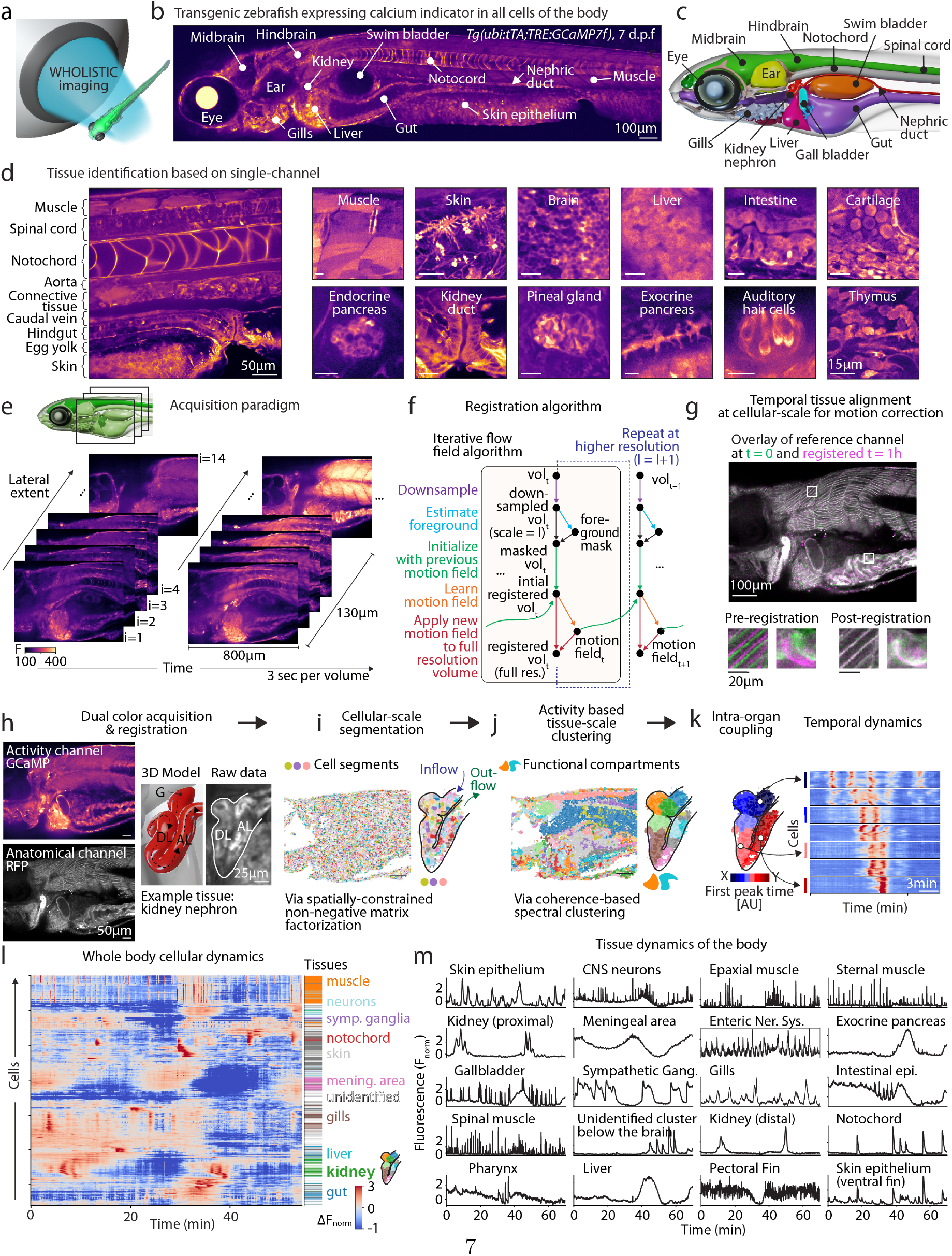
WHOLISTIC imaging of intracellular calcium dynamics across the body. **a**, Schematic of WHOLISTIC (WHole Organism Live Imaging System for recording Tissue and IntraCellular activity). **b**, Transgenic zebrafish (7 days post-fertilization) expressing GCaMP7f pancellularly (*Tg(ubi:tTA;TRE:GCaMP7f)*), imaged with spinning-disk confocal microscopy. **c**, Annotated 3D bio-realistic anatomical model of young zebrafish derived from on Whole-Body Expansion Microscopy (WB-ExM) data. **d**, Single fluorescence channel enables identification of organs and tissues. Enlarged images showing examples of tissues across the body displaying distinct visual textures. **e**, Volumetric imaging acquisition paradigm with typical volumetric acquisition parameters used. Fluorescence intensity variations reflect changes in intracellular calcium. Images covering an area of 800 µm × 600 µm are acquired in steps of ∼ 9 µm, spanning a depth of ∼ 130 µm using a 20× 0.75 NA objective. **f**, Data registration workflow utilizing multi-scale iterative estimation of local motion fields. **g**, Registration results demonstrating temporal alignment of the data. *Top*: overlay of data from the start (green) and 1 hour into an experimental recording (magenta). Enlarged view of myocytes and gut epithelium before (left inset) and after (right inset) registration. **h**, *Left*: dual-color imaging combining pancellular sensor line (e.g., GCaMP7f) with membrane-targeted anatomical reference line. *Right*: example tissue: inset showing 3D model and enlarged view of the kidney nephron. **i**, Cellular-scale segmentation of WHOLISTIC data. Example plane showing the weighted spatial footprints of cell segments derived from Voluseg, a method based on constrained non-negative matrix factorization. Inset showing enlarged view of segmentation results for the kidney nephron (1265 cellular fragments). **j**, Tissue-scale clustering of WHOLISTIC data. Example plane showing the spatial footprints of functional tissue compartments derived from coherence-based spectral clustering. Inset showing enlarged view of clustering results for the kidney nephron (11 functional tissue compartments). **k**, *Left*: Nephron clusters ordered by time to first calcium activity peak within a burst. *Right*: raster of cellular activity within a single nephric burst, grouped by functional compartment and ordered within compartments using Rastermap ^52^. **l**, Body-wide cellular activity raster ordered using Rastermap. On the right spectral cluster assignment shown for some of the major organ groups. For visualization purposes, a random subset of 50 cells per cluster were included in the plot to reduce density of the figure, which were then ordered by Rastermap. **m**, Population-level dynamics. Mean activity of example functional tissue compartments showing distinct dynamic profiles: high-frequency dynamics (e.g.: neurons and muscles), slower evolving activity (e.g.: nephric tissue), tonic bi-stable dynamics (e.g.: sympathetic ganglion) as well as oscillatory activity (e.g.: enteric tissue).

The utility of pancellular imaging is contingent on the ability to discern organs and cell types within densely labeled tissue. The ubiquitous expression of GCaMP7f in tightly packed cells might have led to a conglomeration of indistinguishable cells; however, we found that major organs and tissue types could be distinguished by their distinct visual textures arising from variations in cell size, morphology, subcellular structure, baseline calcium, and sensor expression level (Fig. 1d). Together, this enables the recognition of bodily tissues, including muscle, liver, kidney, brain, and intestine, as well as smaller cellular populations such as the pineal gland and the endocrine pancreas (Fig. 1d), subsequently verified at a more detailed scale using Whole-Body Expansion Microscopy (Methods). Furthermore, we developed a pancellular *Danionella cerebrum* GCaMP transgenic line to adapt WHOLISTIC to adult transparent vertebrates (Extended Data Fig. 1c,d). Thus, the expression of calcium sensors under the ubiquitin promoter has the potential to enable the monitoring of dynamic cellular signals throughout the organism, both developing and mature, while also affording the identification of tissue types, with or without the addition of cell-type specific markers.

To acquire and process spatiotemporal WHOLISTIC data and extract interpretable cellular calcium activity timeseries that can be related to the tissues of origin, we developed a data collection and analysis workflow (Fig. 1e-j, Methods). Acquiring high-quality data across the body was found to be more challenging than imaging the brain due to greater tissue inhomogeneities and complex geometry resulting in increased scattering and diffraction (Extended Data Fig. 1e), such that even advanced forms of light-sheet microscopy^21,43,44,45^, which are often used for whole-brain imaging in zebrafish, were not suitable at the age ranges of interest. We evaluated multiple optical microscopy techniques for rapidly acquiring volumetric data with cellular resolution and found spinning-disk confocal microscopy to be a better option, as it directs excitation light to each scanned point from multiple angles and rejects scattered emission light, making it more robust to diffraction and scattering by the tissue (Extended Data Fig. 1f). As most major internal organs in zebrafish are located in the anterior third of the body^46^, we opted to primarily image this anterior region from the side (Fig. 1e, Supplementary Video 1, Supplementary Video 2).

Smooth and skeletal muscle contractions throughout the body result in the displacement of cells (Supplementary Video 3), such that accurately extracting the activity of individual cells over time requires accounting for this cellular displacement. While motion in the brain is relatively rigid, motion in the viscera is much less constrained, making standard registration algorithms ineffective or slow. To overcome this challenge, we first performed dual-color imaging of calcium dynamics concurrently with an anatomical reference channel to aid alignment (Extended Data Fig. 1g). Next, we developed a registration algorithm based on iterative optical flow estimation^47^ that estimates motion patch-wise (11 *×* 11 pixels) and combines smoothness regularization, signal-to-noise weighting of gradients, and multi-resolution registration to favor convergence to an optimal tissue alignment for each time point (Fig. 1f, Methods). This achieved high-quality alignment, and most cells were accurately registered throughout the experiments (Fig. 1g, Extended Data Fig. **??**a-b, Supplementary Video 4). Subsequent analysis allowed for the identification of single-cell responses to externally applied stimuli, including previously unknown responses of cartilage chondrocytes to cold temperatures and responses to ketamine by cells located at the meningeal boundary of the brain (Extended Data Fig. 3, Extended Data Fig. 4, Supplementary Video 5). Thus, WHOLISTIC enables the analysis of spontaneous and stimulus-driven responses across the animal at the single-cell level.

### WHOLISTIC data processing workflow and identification of Functional Tissue Compartments

To extract cellular-scale calcium activity traces from registered imaging data (Fig. 1h), we employed a segmentation method based on constrained non-negative matrix factorization (Voluseg) (Fig. 1i, Methods)^48^. We tuned this approach to minimize false mergers, favoring a conservative over-parcellation of the data into subcellular segments, which can later be aggregated.

Remarkably, we found that the majority of cells exhibit measurable, time-varying calcium dynamics even at rest (Fig. 1j-m, Supplementary Video 1). While this finding aligns with the well-documented importance of cellular calcium signaling, it has never been demonstrated that most cell types in disparate organs exhibit measurable second-scale calcium fluctuations even in the absence of specific stimuli, nor that the dynamic range of GCaMP7f is sufficiently broad to capture them. We refer to these calcium dynamics as cellular ‘activity’.

The utility of pancellular imaging increases with the ability to readily assign cells to their organs and cell types of origin. While imaged volumes can be manually annotated with the identity of tissue types based solely on anatomical features (Fig. 1d), it is laborious, especially for sparse or highly distributed populations, and misses finer delineations within cellular populations that are defined not by their appearance but by their calcium activity patterns. We thus explored whether a more data-driven methodology, centered on the aggregation of cells with related activity patterns, could more effectively delineate functional anatomical boundaries within and across tissues. This would also provide a more compact, dimensionally reduced and interpretable representation of the data. To this end, we developed Coherence-based Spectral Clustering (Fig. 1j, see Methods), a variant of spectral clustering that utilizes coherence as a similarity metric^49^. Coherence is a frequency-based analogue to correlation, which is invariant to phase lags and thus better groups cells engaged in temporally coupled activity (e.g., traveling waves of activity across tissues) (Extended Data Fig. 4).

Examining the anatomical footprint of the spectral clusters, we found that they predominantly corresponded to sub-regions of individual tissue types, such as the kidney, gallbladder, liver, gut, and muscle (Fig. 1j,k,m, Extended Data Fig. 5). This can be leveraged to organize the data into interpretable and readily identifiable clusters at the tissue level (Fig. 1j,k), which we denote ‘functional tissue compartments’. These can then be annotated with tentative tissue-type identities when discernible. As an example, the WHOLISTIC workflow parcellates the kidney nephron into contiguous functional tissue compartments, each exhibiting distinct dynamics, that together couple into traveling waves that occur in bursts (Fig. 1j,k). The spatial propagation, coupled with the observed quiescence between bursts, suggests that kidney filtration – mostly measured at low temporal resolution^50^ and conceptualized as a continuous or continuously oscillating process^51^ – may in fact be intermittent and pulsatile on shorter timescales. Clusters across the body exhibited a range of calcium activity, showcasing diverse temporal spectra, including oscillatory, pulsatile, and bistable activity (Fig. 1l,m).

Collectively, these findings demonstrate how the ubiquitous expression of molecular sensors, in conjunction with advanced motion-corrected spinning-disk confocal microscopy and the computational workflow for cellular and tissue segmentation, together comprising WHOLISTIC, give access to a rich array of cellular dynamics across tissues of the body, thereby facilitating the study of their interactions.

### WHOLISTIC combined with optogenetics identifies and causally tests coordination of activity within and between bodily organs

#### WHOLISTIC identifies body-wide skeletal muscle coordination

To exemplify the use of WHOLISTIC to study coupling of calcium activity within tissue types, we began by inspecting functional tissue compartments corresponding to skeletal muscle, in which the rise of intracellular calcium induces contraction^53,54^. Muscle compartments were classified based on their distinctive cellular morphology (Fig. 1d), stereotyped spatial location (Fig. 2a), and characteristic activity waveform. Next, the compartments were organized through hierarchical clustering of their calcium activity patterns and visualized by projecting their spatial footprint back into anatomical space (Fig. 2b, see Methods). This approach readily enabled the identification of the primary epaxial and hypaxial trunk muscles, which exhibited correlated activity patterns (as represented by the union of the blue and green clusters in Fig. 2b), underscoring their propensity to contract synchronously, as documented in other species^55^. Additionally, through this method, we could identify the abdominal (also known as hypaxialis), sternohyoid, pectoral fin, and branchial muscles (Extended Data Fig. 6a)^56,57^. More surprisingly, we noted that the antero-dorsal portion of the epaxial muscle was grouped in a distinct functional tissue compartment (‘cervical-epaxial muscle’), and displayed correlated calcium activity with the abdominal muscle (Fig. 2c, *right*), a finding not previously reported. These results can be visualized within the raw data by examining time points at which specific muscle groups are selectively activated (Fig. 2c, *left*).

**Fig. 2:**
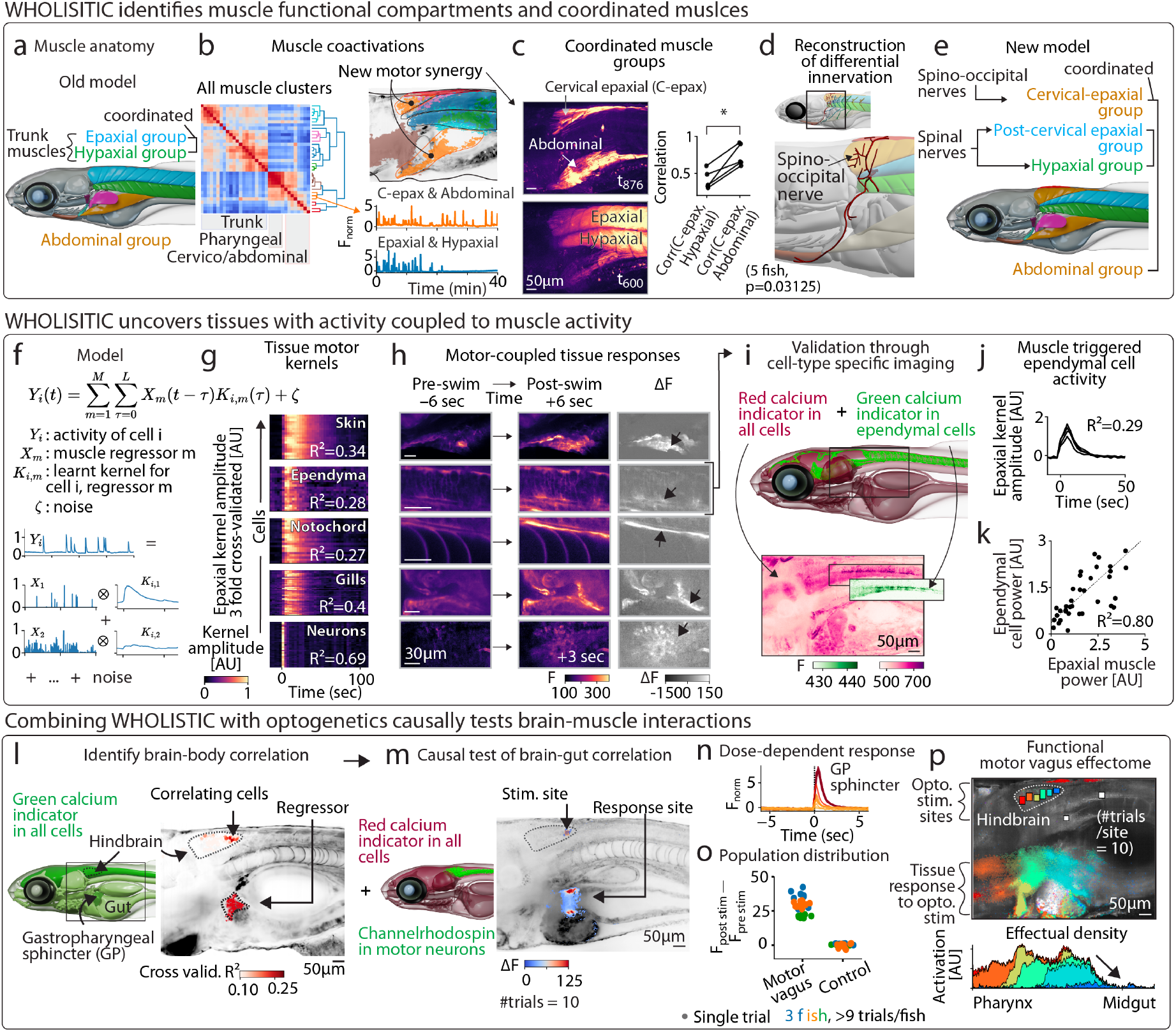
Identification and causal test of intra- and inter-organ coupling through WHOLISTIC. **a**, Model of zebrafish muscle anatomy showing a subset of muscle groups reported in the literature. **b**, *Left*: cross-correlation matrix of muscle functional compartments hierarchically ordered with dendrogram, revealing three main groups: 1) trunk, 2) pharyngeal, and 3) abdominal/cervical-epaxial muscle group. *Top-right*: spatial footprints of muscle clusters (colored as in the dendrogram) overlaid on anatomical reference (gray). The abdominal muscle and cervical-epaxial (‘c-epax’) muscle are part of the same functional cluster (orange). *Bottom-right*: average calcium activity traces from muscle clusters showing distinct activity patterns. **c**, *Left*: selected time-points highlighting co-activation of distinct muscle groups (maximum intensity projection, Δ*F*). *Right*: correlation between cervical-epaxial (c-epax) and abdominal muscles vs. hypaxial muscles (5 animals, one sided Wilcoxon signed-rank test, p=0.3125). **d**, Reconstruction of anterior motor nerve tracts showing (*top*) spino-occipital nerves innervating cervical portion of the epaxial muscle along with the sternohyoid muscles and (*bottom*) spinal nerves innervating post-cervical epaxial and hypaxial muscles. **e**, Revised model of zebrafish muscle anatomy based on WHOLISTIC analysis. **f**, Lag regression model to characterize motor responsiveness of tissues. **g**, Motor response kernel of various tissues and three-fold cross-validated variance explained by the model (averaged across cells shown). **h**, Example of tissue activity 6 sec pre- and 6 sec post-swim and ΔF (gray). *Top to bottom*: 1) ventral fin skin epithelium, 2) midline cells in the spinal cord (later identified as ependyma), 3) notochord sheath cells, 4) gills, 5) neurons in the hindbrain (here 3 sec post-swim activity is shown as activity is short-lived). **i**, *Top*: genetic strategy to validate ependymal cell motor-responsiveness. *Bottom*: sagittal section of transgenic line expressing GCaMP7f in ependymal cells and red calcium indicator pancellularly, with inset of the hindbrain and spinal cord. **j**, Motor response kernel of ependymal cells fit using independent channels shown in (**i** from *Tg(VAChTa:gal4;UAS:CoChR-eGFP)*;*Tg(ubi:tTA;TRE:jRGECO1b)*. **k**, Scatter plot of muscle power (red channel) vs. ependymal cell activity power (green channel) for individual motor events. Dashed line: linear fit capturing 80.0% of variance. **l**, *Left*: schematic outline of the location of the hindbrain and the gastropharyngeal (GP) sphincter. *Right*: a small group of cells located in the hindbrain have high loadings on the GP sphincter regressor. Maximum projection of brain, alpha-weighted by the variance explained by sphincter calcium activity (*R*^2^) in held-out data, overlaid onto anatomical reference (gray), with location of GP sphincter functional compartment shown in flat red. **m**, *Left*: genetic strategy for optogenetic stimulation of motor vagal cells during WHOLISTIC imaging. *Right*: stimulus-triggered average responses overlaid with anatomical reference (# trials = 10, where # stands for ‘number’ throughout); stimulation site and full motor vagus outline indicated. **n**, Gastric sphincter calcium activity showing dose-dependent response to optogenetic activation of caudal motor vagal neurons. **o**, Quantification of change in fluorescence at the gastric sphincter following stimulation of caudal motor vagus or control location along the spinal cord (3 animals, *>*9 trials/animal). **p**, *Top*: optogenetic tiling across the motor vagus to assess the functional topological relationship between motor vagus and visceral organs. Trial-triggered average response of motor vagal nuclei stimulation sites, pixels weighted and color coded according to the ROI (region of interest) inducing the most activity (# trials/location = 10), overlaid with anatomical reference (gray). *Bottom*: plot of responsive tissue volume (activation density) along the anterior-posterior axis of the gastrointestinal tract.

To ascertain the neural underpinning of the independence of activity of the cervical-epaxial muscle from the primary epaxial muscle, we traced the anterior motor nerve tracts using a transgenic line labeling motor neurons, *Tg(VAChTa:eGFP)* (see Methods). This revealed that, although the spinal nerves predominantly innervate the majority of the epaxial and hypaxial musculature, the spino-occipital nerve – known to innervate the ventral sterno-hyoid and pectoral fin muscles^58^ – extends axons dorsally to innervate the cervical portion of the epaxial muscle (Fig. 2d, Extended Data Fig. 6b). This dual innervation of the epaxial muscle supports the observed independent regional activation, consistent with prior studies in other species demonstrating regional activation of the epaxial muscle, with ventral and dorsal fibers recruited during swimming and feeding, respectively^59^. Together, this shows how WHOLISTIC can uncover, even within extensively studied tissue dynamics – muscles – new functional units as well as reveal previously unidentified synergies and independencies among muscle groups (Fig. 2e).

#### WHOLISTIC identifies body-wide skeletal and smooth muscle-coupled dynamics

To demonstrate the ability of WHOLISTIC to identify inter-organ interactions, we next studied the coupling between skeletal muscle and other bodily tissues. While skeletal muscles enable animals to act on the external environment and move, they also generate mechanical forces within the organism, which are detected by mechanosensitive cells throughout the body, triggering a variety of physiological processes^60,61,62,63,64,65^. Connections through the nervous, vascular, and other systems can additionally induce coupling between muscles and other organs. However, the acute effects of muscle activity on non-muscle tissue types, and vice versa, the effects of body organs on muscles, remain largely uncharted.

We thus asked: how does motor activity acutely affect organ and tissue dynamics throughout the body? We examined the temporal relationships between the activation of skeletal muscles and calcium signaling in other cells of the body (Fig. 2f-h) by fitting a lag-regression model to all cells within the imaged volume, in which cellular calcium activity is modeled as the convolution of muscle activity and a cell-specific motor-response kernel (Fig. 2f, Extended Data Fig. 6c,d; see Methods) (we note that this model does not purport to infer causal direction). We quantified motor-related cellular activity throughout the imaged volume using statistical parametric mapping, which highlighted responses in a variety of tissues, including the brain, skin epithelium, notochord sheath cells, and gills (Fig. 2g,h). This coupling exhibited distinct, tissue-specific responses, often enduring beyond the timescale of muscle activity, except in the brain, where a cluster of brainstem neurons showed short coupling timescales (Fig. 2g,h – row 4).

Remarkably, within the brain, beyond motor-coupled neurons^66^, a cohort of cells in the midline of the hindbrain and spinal cord exhibited temporally extended responses to muscular activity, enduring significantly longer than the neuronal responses (∼15 sec) (Fig. 2g,h – row 2). Based on their anatomical position and morphology, we hypothesized that these were ependymal cells, glia-like cells that line the ventricles and the central canal^67,68^. To test this hypothesis, we generated a transgenic line expressing GCaMP7f specifically in ependymal cells (*Tg(foxj1a:GCaMP7f)*), as well as a new pancellular transgenic line expressing a red fluorescent calcium sensor^38^, *Tg(ubi:tTA;TRE:jRGECO1b)*. Crossing these two lines enabled the simultaneous recording of definitive ependymal cell calcium activity, as well as comprehensive whole-body dynamics in independent channels (Fig. 2i) and confirmed that ependymal cells exhibit Ca^2+^ transients subsequent to motor activity, with response amplitudes that are proportional to the motor events’ intensity (Fig. 2j,k, Extended Data Fig. 6e). This suggests that ependymal cells integrate body movements into their cellular state, potentially contributing to ventricular tissue homeostasis or modulation of cerebrospinal fluid (CSF) dynamics^69,70,71^.

The capability of WHOLISTIC and whole-body single-cell analysis to reveal the coupling between ependymal cell calcium levels and muscle activity highlights the sensitivity and efficacy of integrating whole-body imaging with computational analysis, illustrating its potential for the discovery of novel functional properties even amid sparse, distributed cellular populations.

#### Combining WHOLISTIC with optogenetics to causally test brain-body coupling

To expand the analysis to include visceral organs as regressors, we considered relationships between cellular and smooth muscle activity, which exist in tissues forming the gut, the lining of blood vessels, the gallbladder, and others, by incorporating smooth muscle compartments into the lag regression model. Among the various relationships identified, the model captured a relationship between the caudal hindbrain and smooth muscle of the gastropharyngeal sphincter. Indeed, this regressor accounted for the largest fraction of variance of a small group of cells within the caudal hindbrain, specifically within the region of the motor vagal nuclei (Fig. 2l, Extended Data Fig. 6f). As the model is agnostic to causal direction, this is consistent with these motor vagal neurons driving the sphincter calcium activation via the vagus nerve. Given the established connection between the motor vagus and the gut across species^15,72,14,73,74^, this finding represents the identification of a conserved vagus-gut circuit from an unbiased WHOLISTIC analysis.

We sought to probe whether the relationship between this motor vagal area and the gastropharyngeal sphincter was causal. To do this, we combined WHOLISTIC with optogenetics by crossing a transgenic line expressing channelrhodopsin in the motor vagus with the pancellular genetically-encoded calcium indicator line (*Tg(VAChTa:gal4;UAS:CoChR-eGFP)*;*Tg(ubi:tTA;TRE:jRGECO1b)*) (Extended Data Fig. 6g). We found that activation of neurons within the most caudal segment of the motor vagus induces a dose-dependent calcium activation of the gastric sphincter (Fig. 2m-o), recapitulating the observed coupling identified by the model. Moreover, examination of the entire motor vagus through the successive activation of subsets of neighboring vagal neurons revealed a topological mapping between the motor vagus and visceral tissues along the anterior-posterior axis (Fig. 2p), with more anterior motor vagal neurons controlling more anterior pharyngeal muscles^72,14^. We noted that neural activity exerted control of smooth muscle up to the anterior gut but not beyond. To assess whether there was an anatomical underpinning for this spatially restricted control, we visualized the distribution of motor vagal axons along the gastrointestinal tract using Whole-Body Expansion Microscopy (Methods, Extended Data Fig. 6h,i), revealing a significant drop in innervation density at the level of the anterior gut, paralleling the functional results and offering a potential anatomical basis for the functional findings. This combination of WHOLISTIC, optogenetics, and ExM-based anatomical mapping underscores the capacity of this integrated approach to rapidly uncover and mechanistically examine brain-body circuits.

#### Bridging behavior and physiology: WHOLISTIC and Whole-Body Expansion Microscopy uncover ultraslow oscillations associated with motor quiescence and pinpoint their subcellular origin

As a demonstration of the broad applicability of WHOLISTIC, we next considered the converse of movement-coupled modulation of tissue activity: bodily processes that occur during extended periods of motor quiescence. In between phases of sustained motor activity, animals engage in equally important periods of muscle inactivity associated with rest, sleep, and tonic immobility^75,76,77,78^. These quiescent states are not merely passive intervals but active phases of physiological regulation, essential for maintaining organismal health and enabling adaptive behaviors. While the role of neuronal activity during these states has been extensively studied, the contributions of other cell types – both within and outside the nervous system – remain less characterized.

We therefore aimed to identify cells – whether neuronal or non-neuronal, within or outside the brain – that exhibited heightened activity during motor-quiescent periods. We analyzed the variance in cellular calcium activity during episodes of extended muscle inactivity (>10 minutes) compared to periods of regular motor activity. This analysis highlighted a region in the central nervous system that exhibited increased activity during rest (Extended Data Fig. 7a). These signals were identifiable along the midline of the hindbrain and spinal cord, displaying high-amplitude, ultraslow oscillatory activity during motor-quiescent phases, with a periodicity of 3 to 7 minutes (Fig. 3a, Extended Data Fig. 7b,c). In contrast, during periods of motor activity, faster and abrupt muscle-locked activity was observed in these cells, and the slow oscillations were either absent or significantly attenuated (Fig. 3b-d). Imaging neuronal and astrocyte-specific lines (*Tg(elavl3:H2B-GCaMP7f), Tg(gfap:jRGECO1b)*) failed to recover such dynamics, leading to the hypothesis that these slow signals may originate from ependymal cells. Ependymal cells have been associated with sleep and alterations in CSF flow^79,80,81,82,83^ and are juxtaventricular cells, with their cell bodies residing along the central canal and ventricles^68^. However, the oscillations were primarily observed at the base of the hindbrain, above the notochord – an area far from the ventricles or central canal, where ependymal cells are typically assumed to reside (Extended Data Fig. 7d) – bringing into question whether the oscillations originate from ependymal cells. To resolve this discrepancy, we sought to visualize the detailed morphology of ependymal cells that reside deep within the brain.

**Fig. 3:**
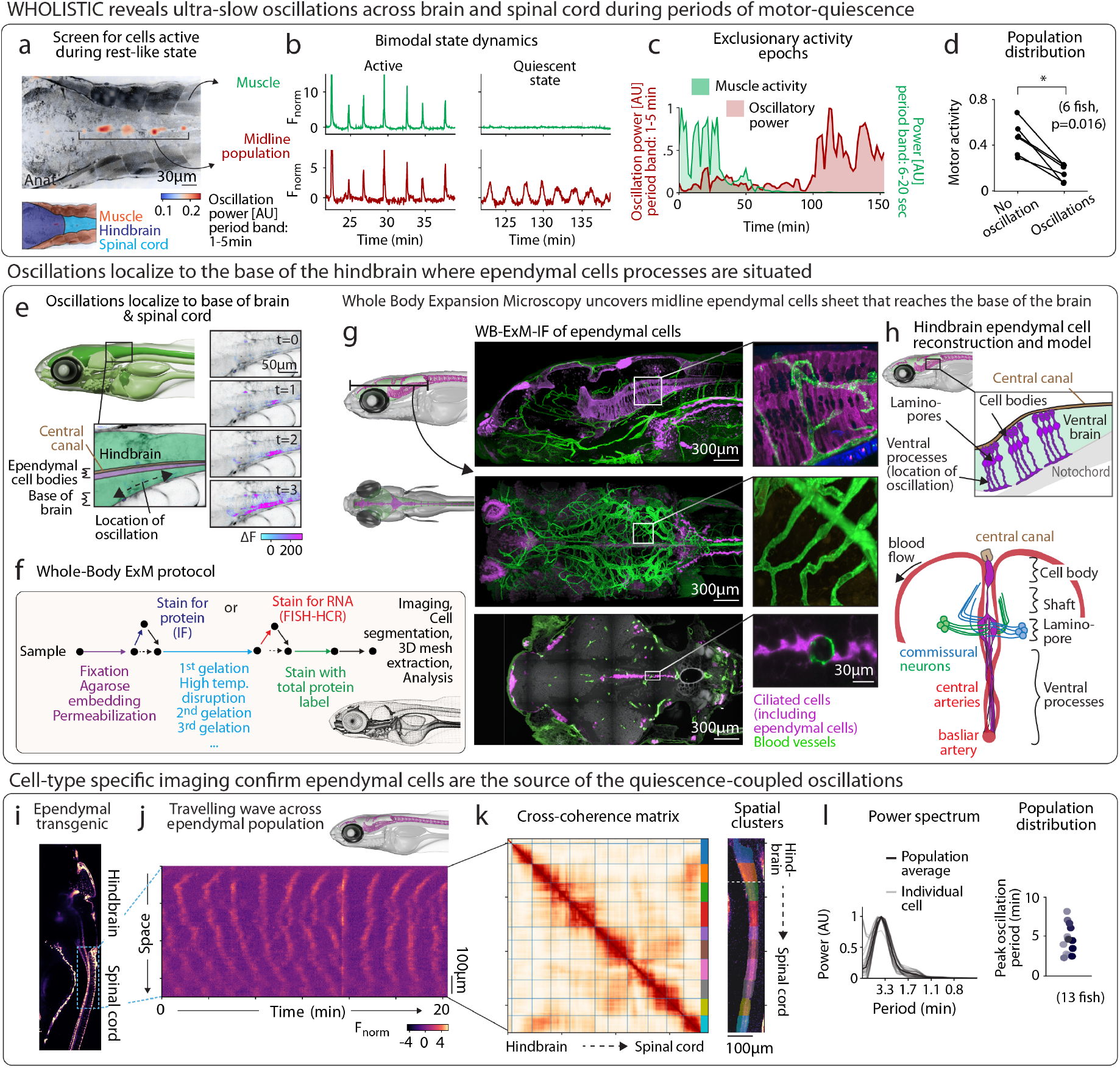
Discovery of motor-quiescence coupled ultraslow oscillations and cell-type of origin. **a**, Midline cells in the hindbrain and spinal cord display high-amplitude ultraslow oscillations during periods of extended motor quiescence. Heatmap showing spatial distribution of oscillatory power (periodicity band: 1-5 min), overlaid on anatomical reference (gray). **b**, Traces during the two activity states. *Top*: active period during which muscle and ependymal cells show coupled activity. *Bottom*: motor-quiescent period during which there is little motor activity and ependymal cells show oscillatory dynamics. **c**, Spectral power of oscillatory cells and muscle activity over a 3 hr window showing anticorrelated dynamics between muscle and ependymal cells (periodicity band: 1-5 min and 6-20 sec for midline population and muscles respectively). **d**, Quantification of motor activity during oscillatory and non-oscillatory periods. Average motor activity during ependymal oscillations pooled from fish exhibiting both motor states (6 fish, one sided Wilcoxon signed-rank test, p=0.016). **e**, Propagation of Ca^2+^ wave along the base of the hindbrain, shown through sequential time points of WHOLISTIC data. ΔF overlaid onto anatomical reference (gray). **f**, Outline of Whole-Body Expansion Microscopy protocol. **g**, Visualization of ependymal cells throughout the brain using WB-ExM-IF. Sagittal and dorsal section of double transgenic sample labeling the ventricular and vascular systems (*Tg(foxj1a:eGFP)* x *Tg(flk1:dsRED-CAAX)*), stained against eGFP (magenta) and dsRED (green), with total protein stain, Alexa488-NHS (gray) (WB-ExM-IF, 10 days post-fertilization, expanded ∼ 2×). **h**, Schematic of individual hindbrain ependymal cells, highlighting structural projections. **i**, Genetic strategy for imaging ependymal cells through the use of Foxj1a promoter, labeling all motile-ciliated cell including ependymal cells. Sagittal section of foxj1a-transgenic line (max. intensity projection). **j**, Kymograph of ependymal cell activity along the hindbrain and spinal cord showing local traveling waves over the course of ∼ 45 min. Data derived from cell type-specific transgenic (*Tg(foxj1a:GCaMP7f)*). In contrast to the periodic activity, an example of synchronous population activation is visible at around minute 12 **k**, Cross-coherence matrix derived from kymograph data in **j**, showing block-diagonal structure indicative of local coherence clusters. Coherence-based k-means clusters projected onto anatomical space, showing spatial continuity. **l**, *Left*: Periodogram of ependymal cell activity. Population mean (dark) and individual cells (light) from example specimen. *Right*: population distribution of peak oscillatory frequency.

#### Whole-Body Expansion Microscopy (WB-ExM) to map ultrastructure, RNA, and protein across the body

Despite the zebrafish’s relative transparency, the accumulation of refractive index variations across the body results in light scattering, making it difficult to visualize the fine anatomy of such midline structures. Existing clearing and expansion methods are either only applicable to younger samples (≤5 days post fertilization), achieve clearing at the expense of extensive proteolytic digestion^84,85^, or require extensive wash steps to clear adequately without expansion^86,87^. To overcome these limitations, we developed an advanced enzyme-free, rapid, and robust Whole-Body Expansion-Microscopy (WB-ExM) protocol^4,88^ (Fig. 3f) that enables high-quality clearing and up to 5× uniform expansion, along with the acquisition of molecular information at subcellular resolution (Extended Data Fig. 8a-h) throughout entire larval and juvenile zebrafish (Extended Data Fig. 8i), as well as mature *Danionella* (Extended Data Fig. 8j). This protocol combines high-temperature disruption with a multi-round embedding strategy, gradually reinforcing and expanding the sample to ensure homogeneous expansion (Methods). WB-ExM retains immunofluorescence signals (WB-ExM-IF) (Extended Data Fig. 8a,b, Supplementary Video 6, see Methods) and, with modifications, is compatible with in situ hybridization (WB-ExM-FISH) (Extended Data Fig. 8d-f, see Methods). In addition, the use of total protein stains yields rich anatomical information (WB-ExM-Histo) with which to contextualize the IF and FISH signals (Extended Data Fig. 8g,h, Supplementary Video 7)^89,90^. To quantitatively assess expansion quality across the sample, we developed PhotoMap, an unbiased method for extracting the deformation field of the gel (Extended Data Fig. 10). Briefly, a fluorescent gel (Extended Data Fig. 10a) is photobleached with a regular grid pattern using a two-photon microscope prior to expansion (Extended Data Fig. 10b), and re-imaged afterwards (Extended Data Fig. 10c). The resulting grid deformation can be readily visualized and quantified (Extended Data Fig. 10d-e). Expansion was found to be largely uniform (Extended Data Fig. 10f-g), except in regions of mineralizing bone, where tissue pinching was observed but remained spatially restricted (Extended Data Fig. 10c-d). These tools complement and bridge existing approaches based on histological, X-ray, expansion microscopy, and electron microscopy methods^46,84,85,91,92^ and serve as a high-throughput approach to obtaining more comprehensive anatomical and molecular data at the whole-body scale, providing a basis for establishing a foundational reference compendium. WB-ExM data provide a vital basis for interpreting, cross-referencing, and rigorously validating the findings obtained through WHOLISTIC.

We used WB-ExM-IF to characterize the fine morphological features of ependymal cells (Fig. 3g, Extended Data Fig. 7e-g) and found that this population of hindbrain ependymal cells, typically considered to be small cuboidal cells^93^, actually extends long and dense ventral projections (∼120 µm), forming a sheet-like structure that runs along the entire midline of the hindbrain, reaching the floor of the brain (Fig. 3g,h, Supplementary Video 8), thereby explaining the spatial distribution of the oscillatory signal. The morphology of these ependymal cells echoes a class of ependymal cells, including tanycytes, that retain morphological features of their progenitor, radial-glia cells^94,95,68^, and are widely present in the adult mammalian brain, suggesting an evolutionarily conserved lineage^96^. Of note, we uncovered that ∼25 µm below the central canal, the ependymal processes expand into a dense set of membranes that ensheathe the axonal crossings of the commissural fibers (Extended Data Fig. 7f), an apposition not previously described, suggesting that ependymal cells may support tangential migration of axons, as well as radial migration along them^97^. We also observed ependymal ensheathement of passing arteries Extended Data Fig. 7g), and evenly spaced lateral projections surrounding neuronal tracts exiting the spinal cord (Extended Data Fig. 7j,k), echoing those recently identified in mice^82^. Such findings, which are not readily observable in non-expanded samples, underscore the utility of WB-ExM in elucidating the cellular and extracellular contexts of functionally analyzed populations and contribute to the generation of new hypotheses. We independently verified these morphological findings by reconstructing hindbrain midline ependymal cells in a published EM zebrafish connectome^98^, confirming their ensheathment of commissural axons.

Cell-type specific imaging of ependymal cells recovered the motor quiescent-locked ultraslow oscillatory dynamics, confirming the ependymal-cell origin of the ultraslow oscillations (Fig. 3i-l). To understand the spatiotemporal structure of ependymal activity, we performed cross-coherence analysis to assess the presence of coupling between the oscillatory dynamics of cells (see Methods). This uncovered a spatial organization manifesting as local waves along the anterior-posterior axis of the fish, encompassing both the hindbrain and spinal cord (Fig. 3j), forming a block-like structure in the cross-coherence matrix with a length scale of ∼75 µm. To quantitatively evaluate this, we derived a method for utilizing coherence as a metric in k-means clustering (see Methods); the algorithm, blind to the anatomical location of the cells, recovered spatially continuous clusters (Fig. 3k). Ependymal cells are known to be interconnected via gap junctions^99,100^, which likely accounts for the localized propagation of the signal. Calcium within ependymal cells has been associated with f-actin-mediated cellular shrinkage, resulting in increased paracellular space between ependymal cells, thereby facilitating efflux through peri-neuronal routes^82^. Collectively, these findings support the hypothesis that these cells form a network extending across the brain and spinal cord, constituting an interconnected system capable of integrating and potentially modulating neural activity, vascular signals, and cerebrospinal fluid (CSF) composition. Moreover, it suggests that the observed state-dependent calcium oscillatory modes of ependymal cell activity may correspond to prolonged periods of altered CSF exchange during extended motor quiescence.

Taken together, these analyses illustrate the efficacy of WHOLISTIC in identifying state-dependent specialized cellular signals within densely labeled tissue and highlight a previously overlooked cellular population involved in a universally conserved tendency of organisms to undergo prolonged periods of motor quiescence. Whether guided by human hypotheses or machine learning techniques, WHOLISTIC offers the potential to reveal physiologically or behaviorally relevant cellular activities, which can subsequently be validated and followed up using cell type-specific transgenics and WB-ExM.

#### Combining WHOLISTIC with physiological challenges to rapidly uncover brain-body circuits

To demonstrate the power of WHOLISTIC for studying organism-wide responses to physiological stress, in addition to spontaneous activity (Fig. 2, Fig. 3), we examined how animals adapt to hypoxia – a condition of reduced oxygen availability that engages systemic responses across tissues and organs. Hypoxia is a fundamental physiological challenge that arises in contexts ranging from adaptation to high-altitudes, to pathological conditions such as stroke and heart failure^101,102^. It triggers a cascade of genetic, metabolic, and cellular adaptations, as well as interorgan communication and behavioral changes, to maintain oxygen homeostasis and prioritize energy allocation to critical tissues^103,104,105,106,107^. However, the dynamic interplay between single-cell activity changes and whole-organism physiology during hypoxia remains poorly understood, largely due to the technical challenges of simultaneously monitoring multiple tissues in real time.

To systematically evaluate the systemic effects of hypoxia across the body, animals were exposed to alternating periods of normoxia (normal O_2_ levels, 21%, ∼15 minutes) and hypoxia (reduced O_2_, 10% O_2_, ∼15 minutes) (Fig. 4a). We observed widespread changes in baseline levels of calcium, quantified by computing the difference in average baseline calcium levels between normoxic and hypoxic phases, referred to as the ‘oxygen modulation score’ (Fig. 4b, see Methods). Notably, gastrointestinal tissues demonstrated a systematic increase in baseline calcium levels during hypoxia (Fig. 4c), potentially attributable to a reduction in the calcium buffering capacity of the cells as a result of diminished oxygen availability^108^. Other organs also showed changes in baseline/low-frequency calcium levels, though at the average tissue level with less relative increase. Given that the arterial blood supply to the brain and gut would contain similar oxygen levels, we sought to understand these disparities in responses.

**Fig. 4:**
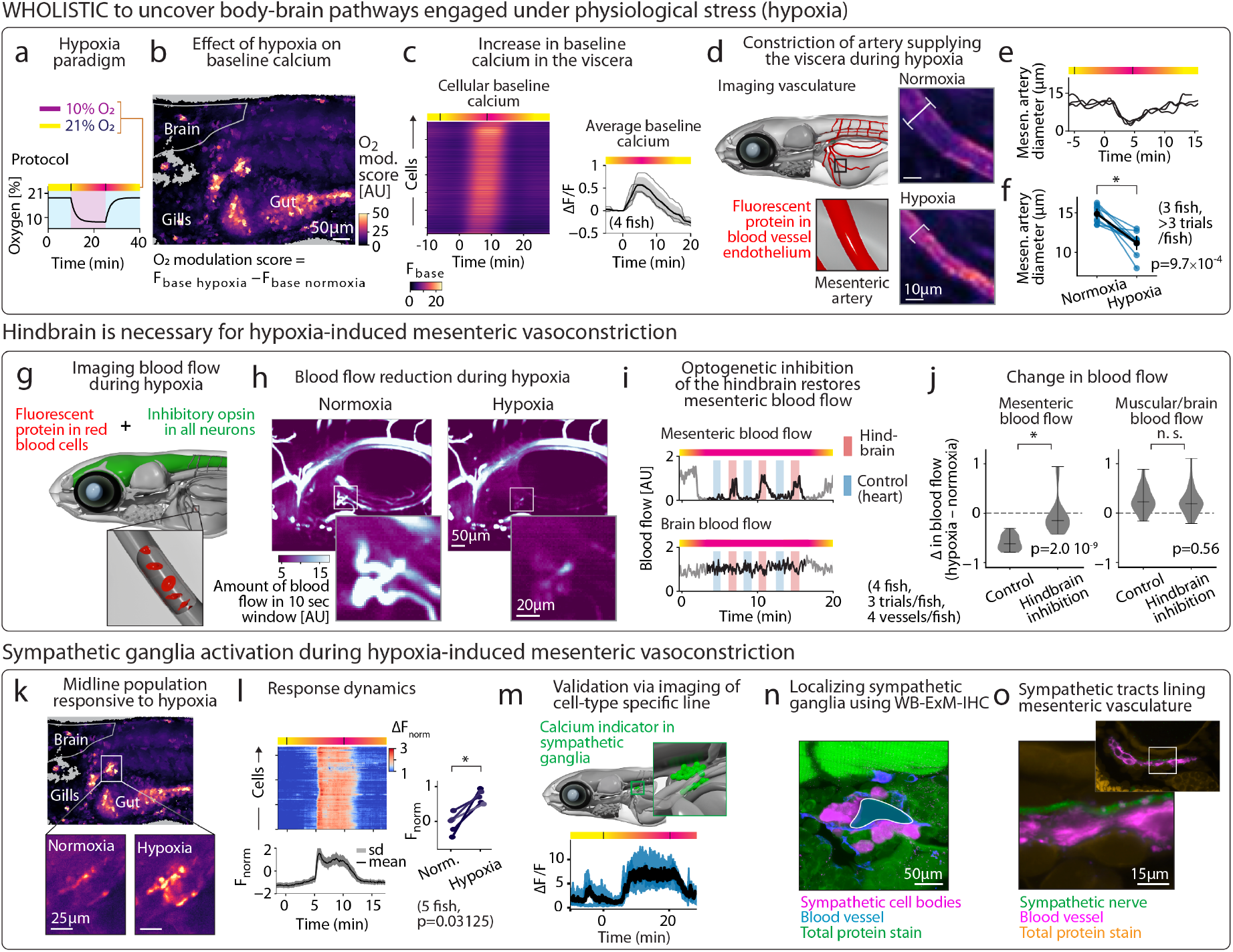
Uncovering of body-wide circuit engaged in response to stress. **a**, Hypoxia paradigm. Animals were subjected to cyclic oxygen levels between normoxia (21% O_2_) and hypoxia (10% O_2_). Black line: measurement of oxygen levels within the bath. **b**, Whole-body map of cellular oxygen modulation score (max. intensity projection). Regions including the gut, liver and kidney displayed increased baseline Ca^2+^ levels during hypoxia. **c**, *Left*: raster of baseline Ca^2+^ levels across visceral tissues over the course of the experiment, ordered by peak time. *Right*: Time course of average hepatic baseline calcium levels in response to hypoxia (dark: mean response, light: individual animals, N=4). **d**, *Left*: genetic strategy to image and estimate blood vessel diameter (*Tg(flk1:dsRED-CAAX)*). *Right*: mesenteric artery during normoxia (*top*) and hypoxia (*bottom*) showing narrower diameter (max. intensity projection). **e**, Temporal dynamics of mesenteric artery constriction. Diameter of mesenteric artery over the course of the experiment (3 trials for the same animal). **f**, Quantification of mesenteric artery diameter during normoxia and hypoxia (3 animals, ≥ 3 trials/animal, one sided Wilcoxon signed-rank test, p=0.00097). **g**, Genetic strategy for optogenetic silencing of the brain whilst imaging blood flow: a transgenic line expressing an inhibitory opsin in all neurons (*Tg(elavl3:gtACR2-EYFP)*) is crossed to a line labeling red blood cells (*Tg(gata1:dsRED)*). **h**, Mesenteric blood flow decreases during hypoxia. Sagittal section of animal, averaged during 10 sec, during normoxia (*left*) and hypoxia (*right*) showing drop in blood flow in mesenteric arteries. **i**, Optogenetic hindbrain inhibition restores blood flow to the gut. Temporal dynamics of blood flow in mesenteric (*top*) and cerebral arteries (*bottom*) in response to hypoxia and to optogenetic neural inhibition. Red: optogenetic inactivation of the hindbrain. Blue: control light stimulus on the heart. **j**, Quantification of blood flow to the mesentery and to the brain under normoxic and hypoxic conditions (4 fish, 3 trials/fish, independent t-test, hindbrain inhibition: p=2.0 *×* 10^−9^, control: p=0.56) **k**, Midline compact cell population shows acute and sustained response to hypoxia, likely being the sympathetic ganglion. *Top*: Same as **b** with location of population shown. Enlarged view of population during normoxia (*left*) and hypoxia (*right*). **l**, *Top left*: raster of functional tissue cluster that maps to midline population (*Bottom left*): mean and standard deviation of raster. *Right*: Average population calcium levels during hypoxia vs. normoxia (5 animals, one sided Wilcoxon signed-rank test, p=0.3125). **m**, Genetic strategy to specifically image the sympathetic ganglia (*Tg(th:gal4;UAS:GCaMP6f)*). Response dynamics of sympathetic ganglion to hypoxia. Individual cells (blue), population average (black). **n**, Localization of the sympathetic ganglia using WB-ExM-IF. Sagittal section of double transgenic sample labeling the autonomic nervous system and vascular systems (*Tg(phox2bb:eGFP)* x *Tg(flk1:dsRED-CAAX)*), stained against eGFP (magenta) and dsRED (blue), with total protein stain, Alexa488-NHS (green) (WB-ExM-IF, 10 days post-fertilization, expanded ∼ 2×) showing sympathetic neurons lining the cardinal vein. **o**, Visualization of juxtaposition of sympathetic fibers (green) along mesenteric artery (magenta). Same sample as in **n**. Vascular system (magenta), autonomic nervous system (green), total protein stain (dark gold).

While considering possible causes for the differential modulation of calcium in the visceral organs compared to the brain, we noticed that the mesenteric artery, the primary artery supplying the visceral organs, appeared constricted during episodes of hypoxia (in unregistered imaging data where tissue motion is visible). This suggests a reallocation of oxygen across the body by rerouting blood flow during physiological stress, a conserved response^106,109^ that has never been imaged in real time. To assess this arterial constriction in more detail, we imaged a transgenic line that selectively labels blood vessels, *Tg(flk1:dsRED-CAAX)* (Fig. 4d) and confirmed the constriction of the main mesenteric artery during hypoxia (Fig. 4e,f, see Methods). As the diameter of a blood vessel is an indirect indicator of blood flow, we imaged a transgenic line labeling red blood cells, *Tg(gata1:dsRED)* (Extended Data Fig. 9a), and found a near-complete cessation of blood flow to the gut during hypoxic conditions, which recovers upon returning to normoxic conditions (Extended Data Fig. 9b, Supplementary Video 9). In contrast, blood flow to the brain and muscle remained largely unaltered, potentially explaining the difference in baseline modulation between these tissues. Thus, consistent with mammals and certain aquatic animals^106,109,110^, WHOLISTIC shows that hypoxia induces a rapid redistribution of blood away from the gut, and reveals the full temporal dynamics of the surprisingly fast redirection of oxygen as well as its spatial distribution, and concurrent activity changes in other cells across the body.

To test whether the reduction in gastric blood flow was neurally regulated, we inhibited the nervous system using the anesthetic tricaine and observed that visceral blood flow was no longer diminished during hypoxia (Extended Data Fig. 9b), suggesting that the reduction in visceral blood flow is mediated by neurons, and showing that this brain-body circuit is already developed at 7 days post fertilization. To more directly test this, we optogenetically inhibited the hindbrain during hypoxia while measuring blood flow (Fig. 4g, Extended Data Fig. 9c). We found that inhibiting the hindbrain induced, within a few seconds, the return of blood flow to the gut (Fig. 4h-i). In spite of the hypoxic condition, near-normal blood flow was maintained for as long as the hindbrain was inhibited, with no corresponding effect observed when directing the stimulation light to a control location (Fig. 4j). Thus, hindbrain activity is necessary for the constriction of the mesenteric artery during hypoxia, which we speculate serves to redistribute oxygen, thereby preserving energy for the brain and other tissues whose activity must be prioritized, and may also be a control mechanism to decrease enteric cellular activity.

During hypoxia, we also observed a set of cells in the medial plane, below the notochord, that showed a systematic and sustained increase in activity during hypoxic periods (Fig. 4k,l). Given the location of these cells, and that sympathetic neurons are known to induce vasoconstriction via noradrenergic activation of vascular smooth muscle cells^111^, we hypothesized that this cluster might correspond to a sympathetic ganglion. Using cell type-specific imaging (*Tg(th:gal4; UAS:GCaMP6f)*)^112^, along with WB-ExM-IF and WB-ExM-FISH, we showed that this group of cells indeed corresponds to sympathetic neurons and that they send axons along the mesenteric artery (Fig. 4m-o, Extended Data Fig. 9d)). Together, these results establish a brain-to-mesenteric artery pathway already functional at 7 days post fertilization. Furthermore, they illustrate the efficacy of WHOLISTIC and its combination with optogenetic perturbations, as an investigative tool for studying whole-body physiology in the context of specific physiological stressors, demonstrating its capability to uncover and causally examine brain-body pathways mediated through specific cell types. This work, moreover, establishes the young zebrafish as a powerful model for studying the consequences of stress-induced blood shunting on enteric physiology, and more generally, organism-wide feedback loops engaged by stress.

## Discussion

The organism has long been acknowledged as a cohesive entity, comprised of a network of cells that communicate and coordinate across multiple scales^113^. The elaborate feedback loops embedded within this network enable animals to respond to, predict, and prepare for both internal and external challenges. While many regulatory pathways—such as those governing hunger, glucose balance, and immune responses—are well characterized^114^, the complete picture of organism-wide control systems remains a formidable challenge. Given the interdependence of these cellular interactions, considering the organism as an integrated whole is essential for full comprehension^2,17,1,115^.

Despite this acknowledged complexity, methodologies that probe vertebrate organisms in their entirety while retaining cellular-level access have remained elusive. This study introduces a novel approach to elucidating whole-body cellular dynamics through the imaging and analysis of genetically encoded sensors expressed ubiquitously across all cells, thereby offering unprecedented insights into the integration of cellular interactions and organism-wide control systems.

Future implementations of WHOLISTIC in freely behaving animals will enable the study of body-wide cellular control systems in naturalistic settings, rather than in an embedded preparation. This advancement will facilitate the investigation of the dynamic interplay between complex behavioral states and strategies, such as goal-driven navigation and sleep-wake transitions, and organism-wide physiology. A parallel route to such studies will occur through the creation of experimental systems that incorporate multimodal virtual-reality environments^116^ composed of, for example, visual, temperature, chemical, mechanical, and other types of stimuli which respond to an animal’s behavioral output. Combined with the further improvements in the ability to monitor and activate cellular signals across the entire body in closed-loop through advances in protein engineering and microscopy^18^, this will open up fundamentally new avenues of scientific inquiry.

The use of calcium sensors^32,38^ serves as an effective proxy for cellular activity due to calcium’s ubiquitous role in cellular processes^117^. The finding that most cells have calcium-activity dynamics within the range of standard sensors has enabled the extraction of a plethora of cellular dynamics, but this is still a limited view of the full dynamics within and between cells, which occur through signaling via a multitude of molecular, voltage, and mechanical pathways. Further measurements of additional molecules and physical cellular properties will provide deeper insights. Indeed, the WHOLISTIC methodology is extendable to co-imaging with other signals within the rapidly expanding repertoire of molecular, voltage, and mechanical sensors^42,41,118,119,39,120^, allowing access to a variety of additional information channels such as metabolic states and molecules like hormones, ATP, and neuromodulators that convey communication signals between cells across the organism.

Such additional measurements will enrich our data-driven understanding of organism-wide physiology, but due to the size and complexity of multicellular interactions will necessitate advanced analytical and modeling frameworks that should leverage advances in modern AI algorithms including Graph Neural Networks^121^, possibly in combination with large-scale biophysical models and model-free methods for predicting dynamics, such as Empirical Dynamic Modeling^122^. Furthermore, platform automation^123^ and AI-in-the-loop systems that combine measurements of whole-body cellular dynamics with online cell-targeted perturbation will be useful not only for reaching fundamental biological insights but also for prototyping future real-time interventions for medical conditions^124,123,125^.

The fact that all findings presented here — pertaining to different cell types, in different parts of the body, under different physiological assays and screens— initially derived from the same WHOLISTIC methodology underscores its wide-ranging applicability to fundamental biology and to medical research. The methodology can be used as a discovery tool, while validation and deeper insight can be gained using more specific molecular techniques, as performed here to understand phenomena like the brainstem’s control of blood flow redistribution during physiological stress and the molecular identification of the quiescence-related signals as originating from ependymal cells.

Cellular function is determined by a cell’s molecular and biophysical properties, its tissue environment, and connectivity to other cells^88,126^. The combination of organism-wide cellular activity imaging, the concept of ‘functional tissue compartments’ which merges function and anatomy, and whole body tissue expansion methods offers the integration of whole-body cellular activity measurements and the molecular, cellular, and connectivity properties of cells across the body. Developing accurate, body-wide registration methods to match the in-vivo to the ex-vivo expanded data cell-by-cell will enable analyses and insights at an enhanced level of depth and detail, including, for instance, matching sets of hormone-secreting neurons with neurons expressing the corresponding receptors^127^. Much information throughout the body flows through the nervous system, the central and peripheral nervous systems, the connectivity of which is becoming more readily accessible through light-based connectomics^128^ and improved machine learning algorithms for connectome reconstruction^129^. The current work will seed the creation of an atlas encompassing the mechanistic, molecular basis for the observed functional coupling at cellular detail across the whole organism.

Beyond enabling progress in fundamental biology, WHOLISTIC enables screens in disease models that allow for tracking the effects of pathologies and potential treatments at multiple spatial scales from cells to the whole body and at temporal scales from seconds to days. The majority of drug-screening workflows depend on observing a subset of drug-induced phenomenology such as molecular binding properties^130^, effects on individual cells or subset of tissues^131^, or influence on behavior^132,133^. The approach presented here offers a complementary lens through which to observe the impact of drugs and genetic interventions on the entire body, enabling the identification of unanticipated off-target consequences in organ systems not under direct study, as well as the discovery of unexpected benefits in tissues not previously considered. It would also reveal outcomes arising from system-wide interactions that could be missed when studying cells, tissues, or organs individually, rather than collectively within their native whole-body environment.

In conclusion, WHOLISTIC constitutes a bridge between cellular and organismal physiology, systems neuroscience and behavior, ushering in a new frontier in systems biology. By embracing the complexity of organisms while retaining the individual cell as a fundamental unit of analysis, this work affords a holistic yet mechanistic approach to unravel the complete cellular interplay governing health and disease in organism-wide networks.

## Methods

### Experimental model and subject details

#### Zebrafish husbandry

Zebrafish were reared at 28.5 °C in 14-10 h light-dark cycles (conductivity 1000 µS, adjusted via Instant Ocean Sea Salt (∼ 30 g/L), pH 7.0, adjusted using sodium bicarbonate (∼ 30 g/L))^134^. Zebrafish from 5 to 14 days post-fertilization were fed rotifers and used for experiments. All experiments complied with protocols approved by the Institutional Animal Care and Use Committee of Janelia Research Campus. Zebrafish sex cannot be determined until ∼4 weeks post-fertilization^135^, so the sex of the experimental animals was unknown.

#### *Danionella* husbandry

*Danionella cerebrum* were reared at 26.5°C in 14-10 h light-dark cycles (conductivity 450 µS, pH 7.5). Feeding protocols were adjusted according to age: 1) 5–28 days post-fertilization (dpf): rotifers were administered once daily, 2) 16–28 dpf: in addition to rotifers, GEMMA 75 was provided twice daily, 3) 29 dpf and beyond: the diet composed of GEMMA 75 twice daily and Artemia once daily. Adult *Danionella* were maintained in group housing with stock density ∼45 fish in 3.5 liter tanks (Tecniplast). Fish younger than 6 weeks old are sexually immature and could not be sexed; the sex of fish older than 6 weeks are mentioned in the main text. For egg collection, 10 cm long custom-made acrylic tubes were used. All experiments complied with protocols approved by the Institutional Animal Care and Use Committee of Janelia Research Campus.

#### Zebrafish transgenics and transgenesis

Transgenic zebrafish were maintained in *Casper* or *Nacre* background^136^. All lines were generated using the Tol2 system^137^ and genes codon optimized using CodonZ^138^. Codon-optimized GCaMP7f and jRGECO1b were synthesized (Twist) and used for subsequent cloning. For cloning, restriction digest cloning was used throughout and all genes (GCaMP7f, jRGECO1b, tTA) were cloned with a preceding Kozak sequence and followed by an SV40 poly(A) signal sequence. Plasmids (150 ng/µl), along with Tol2 transposase mRNA (50 ng/µl), were co-injected (0.5 nl total injected volume) into one-cell stage embryos. Embryos were screened at 7 days post-fertilization for expression and positive embryos were reared to maturity. At maturity, these adults were individually screened for dense expression in progeny, and the best founders were retained.

The ubi:tTA and TRE elements were obtained from multiple plasmids, gift from Daniel Feliciano and Isabel Espinosa-Medina. These included a ubiquitous promoter containing vector, or p5E-ubi^33^ (Addgene #27320), a vector containing the tTA advanced Tet-off transcriptional activator from pTet-Off Advanced Vector (Takara, 631070) inserted into the multiple-cloning site (MCS) of the pME entry vector^139^, and a vector containing the tetracycline responsive element promoter p5E-TRE^140^. To generate the *Tg(ubi:tTA;TRE:jRGECO1b)* animals, the ubi:tTA;TRE elements were cloned and a codon-optimized jRGECO1b sequence placed downstream. To generate the *Tg(ubi:tTA);Tg(TRE:GCaMP7f)* animals, plasmids containing ubi:tTA and TRE:GCaMP7f were independently cloned and co-injected at equimolarity. To generate the *Tg(foxj1a:GCaMP7f)* animals, the Foxj1a promoter was cloned (Addgene plasmid #163829^100^) and a codon-optimized GCaMP7f sequence placed downstream using restriction digest cloning. To generate the *Tg(elavl3:gtACR2-eYFP)* transgenics, the promoter was cloned using a known Elavl3 promoter sequence^141^ and gtACR2-eYFP sequence placed downstream^142^. To generate the *Tg(β-actin2:mCherry-CAAX; myl7:GFP)* transgenic line the *β*-actin2 promoter was cloned (Addgene plasmid #82583,^143^) and mCherry-CAAX sequence placed downstream. For optogenetic activation of motor vagal neurons, the transgenic lines *Tg(VAChTa:Gal4)*^144^ and *Tg(UAS:CoChR-eGFP)*^*jf* 44 48^ were utilized. To record activity of the sympathetic ganglia, the transgenic lines *Tg(th:Gal4)*^112^ and *Tg(UAS:GCaMP6f)*^*jf* 46 48^ were crossed and imaged. To quantify blood vessel diameter and track blood flow, the transgenic lines *Tg(flk1:dsRED-CAAX)*^145^ and *Tg(gata1:dsRED)*^146^ were imaged. To check the colocalization of cell located at the meningeal border of the brain with neurons, astrocytes and basement membrane, the transgenic lines *Tg(elavl3:H2B-jRGECO1a)*^**?**^, *Tg(gfap:jRGECO1b)*^48^ and *TgBAC(lamC1:lamC1-sfGFP)*^147^ were respectively used. Additional lines used for expansion microscopy were: *Tg(isl1CREST-hsp70l:mRFP)*^148^, *Tg(phox2bb:eGFP)*^149^ and *Tg(foxj1a:eGFP)*^150^.

#### *Danionella cerebrum* transgenics and transgenesis

To generate pigmentless *Danionella cerebrum* mutants, we employed the CRISPR-Cas9 genome editing technique to disrupt the function of the *mitfa* gene, following established protocols^26^ and utilizing a *mitfa*-targeting gRNA (sequence: CAGCATTATACACTAAGAGT). Mutations were confirmed via PCR amplification and subsequent sequencing, resulting in the establishment of *Danionella cerebrum mitfa*(-/-) colonies.

To generate the *Danionella cerebrum* WBI transgenic line, *Tg(ubb*^*R*^*:jGCaMP8m)*, a Tol2 vector was constructed containing the ubb^*R*^ promoter^151,152^, kindly provided by Dr. Balciunas and Dr. Lazutka. Following the promoter, a zebrafish codon-optimized jGCaMP8m sequence^152^ and the SV40 polyadenylation signal were arranged sequentially. This plasmid (25 ng/µl), along with Tol2 transposase mRNA (20 ng/µl), was co-injected (0.5 nl total injected volume) into one-cell stage *mitfa*(-/-) embryos. Embryos at 3 days post-fertilization were screened, and those exhibiting strong jGCaMP8m expression were reared to maturity. Upon reaching adulthood, these fish were screened collectively, and the most robust founders were selected for further study.

### Experimental procedures

#### Sample preparation for functional imaging

Prior to imaging, zebrafish and *Danionella cerebrum* samples were embedded on their right side in a drop of 2% low-melting point agarose (Sigma, A9414) in a glass-bottom Petri dish (Mattek, P35G-1.5-14-C, 35 mm). The right pectoral fin was moved away from the flank of the fish to point either tangentially or anteriorly such as to prevent it from covering visceral organs. Following agarose solidification, the dish was filled with E3 fish water. Samples were left to rest and settle for 30 min prior to the imaging session in order to minimize drift during the experiment. Cutaneous respiration is considered to be able to meet the oxygen needs of young zebrafish^**?**^ and thus agarose was not removed from the gills, nor mouth, and no oxygen perfusion is required.

#### Microscope and data acquisition

Functional data were acquired using a Nikon spinning-disk inverted confocal microscope (CSU-W1) with a 20× 0.75 NA air objective (field of view of 800 µm x 600 µm, working distance 1 mm) or a 40× 1.15 NA water immersion objective. For dual-color acquisition of GCaMP7f and mCherry signals, excitation lasers 488 nm and 594 nm were used (3-4% and 3-5% power). Emitted light was split using a 560 LP dichroic and passed through emission filters (525/36, 610 LP) before reaching the cameras (Hamatsu ORCA-Fusion BT). For jRGECO1b imaging, a 561 nm laser line was employed with a 610/75 nm emission filter. Camera exposure times were set between 120 ms and 200 ms. A piezo motor was employed to acquire fast z-stacks with a typical inter-plane interval of 7-9 µm, resulting in a volumetric scan rate of ∼0.3 Hz. Smaller z-steps were used for more targeted investigations (e.g., during vascular imaging).

#### Ketamine treatment

Animals were imaged for a 25 min baseline period, after which ketamine was manually added to the imaging dish to achieve a final concentration of 100 µg/ml (400 µM).

#### Tricaine treatment

Animals were treated with 750 µM tricaine (MS-222, Sigma, E10521-10G) diluted in E3 water. The samples were incubated with the drug for 15 min prior to and throughout the duration of the experiments.

#### Cold stimulus delivery

Fish water was cooled to 10°C and 350 µl of cold water was manually added to the imaging dish after 5 minutes of baseline imaging. To rule out any neural influence or motion, fish were anesthetized prior to and during the experiment with tricaine (MS-222, Sigma, E10521-10G, 750 µM).

#### Hypoxia treatment

Oxygen levels were programmatically varied using an Okolab O_2_ controller module, controlled via the Nikon spinning-disk microscope software (Elements). The module varies O_2_ levels by altering the ratio of N_2_ and O_2_ mixed and the oxygen levels at the chamber inlet are recorded. A baseline period (21% O_2_) of at least 5 min was recorded, after which oxygen levels were lowered to 10% for 10 to 20 min, after which levels were returned to 21%. Oxygen levels were measured in the bath using an oxygen micro-optode (Unisense, O_2_ MicroOptode) Fig. 4a. We note that hypoxia is expected to cause changes in acid-base balance within cells^**?**^ . As the aim was to study the physiological consequences of hypoxia, we did not attempt to control tissue pH.

#### Optogenetics

A commercial integrated DMD module within the Nikon spinning-disk confocal microscope was used for all optogenetic experiments in conjunction with a 488 nm LED. The duty cycle of the DMD was set to 10%, and LED power set to ∼ 8-10%. A dual-color reference image stack was acquired at the beginning of the experiment to get the anatomical location of the cells expressing CoChR-eGFP. Based on this 3D volume, ROIs to be illuminated were defined in 3D via the graphical user interface and programmatically stored. For optogenetic activation of motor vagal cells (Fig. 2), stimulus intensity was initially calibrated by collecting a dose-response curve with progressively increasing stimulus intensity until a reliable response was observed in the neurons being stimulated (∼8% LED power). Once calibrated, the experimental protocol was programmed using Nikon software (Elements) with inter-stimulus intervals of 10 seconds, with ROIs selected randomly (choose-without-repeat) or sequentially. ROIs were illuminated for varying durations ranging from 10 to 500 ms during dose-response experiments, and for all other experiments for one frame (∼180 ms). The data and metadata were extracted using the Python package ND2. For each trial type (i.e., specific ROI (region of interest) stimulated) the difference in activity pre- and post-stimulus was extracted and averaged over a ∼5 sec window (2 frames). To display the data within a single graph, the averaged stimulation-induced maps were combined, with each pixel colored according to the ROI inducing the most change in fluorescence, alpha-weighted by the magnitude of change with alpha set to 0 when the change was below noise level (estimated by the average change induced by the control ROI). For optogenetic inhibition of the hindbrain (Fig. 4), the same stimulus intensity was used as in the optogenetic activation experiment and was confirmed to result in loss of all motor output. After 5 min of onset of lower oxygen levels (10%), an ROI over the hindbrain or the heart (control) was illuminated in alternation for one minute followed by one minute of no illumination. After three alternating trials of each location, oxygen levels were returned to 21%.

## Data Processing Workflow

### Registration

The registration pipeline is based on local iterative motion estimation.

#### Iterative patch-wise optical flow

We refer to the static image as ‘template image’, and refer to the data that we aim to transform as ‘moving image’. The key part of the registration algorithm is the iterative patch-wise optical flow module. This module estimates motion using a modified version of the optical flow algorithm^47^. It aims to minimize the intensity difference between the template and the moving image while encouraging the motion field to vary smoothly over space. We formulate an optimization problem defined by the following loss function (for clarity, the one-dimensional version is described; in practice, this is extended to three dimensions):

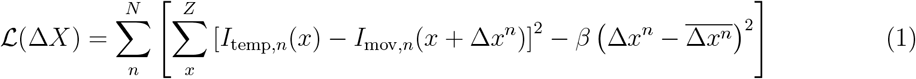

where Δ*X* ∈ *R*^*N*^ is the motion field to be estimated, Δ*x*^*n*^ is the n^th^ element of Δ*X, N* is the total number of patches the image is split into, *Z* is the number of pixels per patch, Δ*x*^*n*^ is the estimate of the spatial motion for patch *n* and 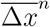 is the average Δ*x*^*n*^ for the neighboring patches of *n*.

This original optimization problem is non-convex and cannot be efficiently solved. Therefore, we re-formulate the problem into one that can be solved iteratively through a set of convex problems using a modified version of the optical flow algorithm^47^:

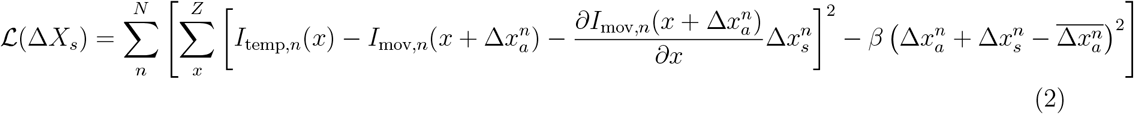

Computing spatial intensity gradients can be prone to noise if done pixel-wise; to overcome this, a locally averaged estimate of the spatial intensity gradient is used. The image is divided into patches of ([2*r* + 1] *×* [2*r* + 1]) pixels and the average spatial gradient is computed (*r* = 5 pixels). The accumulated motion 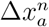 is estimated and optimized per patch. Once convergence is reached, the motion between patch centers is linearly interpolated. The estimated motion is also desired to be somewhat smooth over space. Therefore, the loss function is augmented with a penalty term weighted by *β*, which penalizes the difference between each motion estimate of a patch’s motion and the average of the immediately adjacent patches. This individual optimization step is a quadratic function and thus has a closed-form solution. Once a motion step Δ*X*_*s*_ is computed, the accumulated motion is updated and linearly interpolated between patch centers, then the accumulated motion is applied to the moving frame, and this is iterated until the loss ceases to decrease beyond a user-defined threshold or reaches the max iteration number of each pyramid layer *N*_*iter*_. To take into account that it is the accumulated motion which ought to be spatially smooth, the terms weighted by *β* are the accumulated patch and patch neighborhood motion. Furthermore, to enable the parallelization of calculations, the gradients of the previous time steps are used for the neighborhood motion estimation.

#### Image processing scheme and registration reliability mask

First, one needs to define the template and the moving image. For short datasets, the first frame serves as the template, and each subsequent frame is treated as the moving image. This setup allows for the alignment of all frames one by one to the initial frame. For longer-duration datasets, where bleaching occurs, a floating template is introduced along with an initialized motion field. The floating template is defined as the median of the last *N*_template_ selected motion-corrected frames, while the initial motion field is chosen as the one closest to the center of these *N*_template_ motion fields.

To enhance the algorithm’s robustness to noise, the ‘foreground’ in both the template and the current frame are defined. This step excludes regions containing moving immune cells, which appear as small, bright, connected components which otherwise introduce distortions into the estimated motion field. After pre-processing the data, pyramid downsampling is performed, a technique that reduces image resolution by iteratively smoothing and subsampling the image. This reduces the size of the template, moving image, and the initialized motion field to 1/2^*L*^ of their original dimensions in the x and y directions, where *L* is the number of levels. The original size along the z direction is maintained. Based on specified patch size parameters, iterative patchwise optical flow registration is performed on the downsampled data, which gives a rough estimate of the motion field. After this, the data is upsampled and the motion field further refined, thereby enhancing its accuracy. Once fit, using the refined motion field, the calcium channel is corrected by relocating the moved cells to their original positions as seen in the first frame. This systematic approach provides robust frame alignment and motion correction, accommodating dynamic cell movements and variations in image intensity throughout the functional recording. Finally, to filter out any data that is either not sufficiently well or unreliably registered a registration reliability mask is computed. To identify unreliable textures, characterized by low spatial intensity gradients, the amplitude of the spatial gradient of the template image is computed and standardized using z-scores. Regions exhibiting a score below 0 have little texture and are flagged as potentially unreliable; in addition the mean squared error is computed for each patch across time, and all patches whose error is above a user-set threshold are discarded from downstream analysis.

#### Parameters and Implementation

The algorithm was executed with *β* = 0.01 (smoothness parameters), *r* = 5 (motion correction patch size, measured in pixels), *N*_*L*_ = 4 (number of pyramid layers used for motion correction), *N*_template_ = 5 (number of frames incorporated in the floating template), *N*_*moving*_ = 5 (number of frames to consider when initializing motion), *N*_*iter*_ = 10 (max iteration number of each pyramid layer). The code was developed and run in MATLAB and is available on GitHub (https://github.com/Weizheng96/WBI-registration).

#### Validation

To validate the registration at single-cell resolution, the transgenic line used for the registration, pancellular membrane marker (*Tg(β-actin2:mCherry-CAAX)*) was crossed to a sparse transgenic line, (*Tg(phox2bb:eGFP)*), to use the later as held-out ground truth. Dual-color data was acquired for one hour. Using the reference channel alone, the motion field was estimated and applied it to the held-out green (sparse) channel. In both unregistered and registered data, individual cells within the sparse held-out channel were manually located in the ‘template’ (first time point) and 20 and 60 min into the recording within Fiji software. Displacement distance was subsequently extracted for individual cells.

### Registration - method comparison

#### Synthetic dataset generation

To benchmark registration performance under controlled but challenging conditions, we created a library of three–dimensional simulated recordings in which signal-to-noise ratio (SNR), motion smoothness and displacement amplitude were varied independently.

#### Seed volume

For each *z*–slice we selected an anatomically rich 256*×*256px region and concatenated 13 consecutive slices to form a 256 *×* 256 *×* 13 voxel reference stack (*I*_ref_).

#### Motion field synthesis

A zero-mean, unit-variance 3-D Gaussian random field *G*(*x, y, z*) was convolved with an isotropic Gaussian kernel of standard deviation *σ* = *rigid score* (equivalent to the `FilterSigma` parameter). The smoothed field was normalised to unit peak magnitude and scaled by an *amplitude* factor *A* to yield the final displacement field

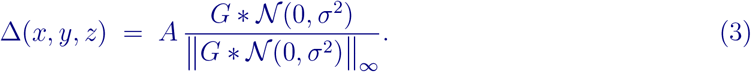

Larger rigid scores therefore produce broader, more spatially coherent motion, whereas increasing *A* raises the maximum voxel displacement.

#### Noise injection

Additive white Gaussian noise *η* ∼ 𝒩 (0, 1) was applied identically to the reference and the warped (moving) image: Nine noise levels (1.4^0^ … 1.4^9^), sixteen rigid scores (5–20 px), and ten amplitudes (1–10 px) were explored. When one parameter was swept the remaining two were fixed at *noise level*=5, *rigid score*=20 px, and *amplitude*=5 px. Eight stochastic realisations were generated for every parameter combination and averaged to obtain a robust estimate.

#### Performance evaluation

For every simulated pair we measured two mean–squared errors (MSE):

1. **Image fidelity:** MSE between *I*_ref_ and the registered image *Î*.
2. **Motion fidelity:** MSE between the ground-truth displacement field Δ and the field recovered by the algorithm 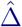.

Only voxels inside a validity mask (`validMap`) contributed to these metrics. The mask excluded (i) a 10-px border in all directions to avoid extrapolation artefacts, (ii) low-texture regions (empirically defined by local gradient magnitude *<*!1, SD of the whole volume), and (iii) trivially small motions (|Δ| ≤ 1 px) that would otherwise bias global averages.

#### Benchmark algorithms

The proposed method was compared against four widely used non-rigid registration frameworks, each tuned for volumetric calcium-imaging data.

#### Suite2p (MATLAB GPU port)

Multi-plane registration was enabled; the field of view was partitioned into 40 *×* 40 XY blocks with 2-px overlap. Bidirectional and plane-to-plane alignment were activated.

#### NoRMCorre

Parameters were set to `grid_size=[32, 32, 1], mot_uf=[4, 4, 1], max_shift=[15, 15, 5], max_dev=[3, 3, 1], and overlap_pre=[4, 4, 1]` .

#### Demons (SimpleITK)

Intensity histograms were matched (1024 bins, 16 match points, background threshold = mean). The algorithm ran for 200 iterations with Gaussian-smoothed updates (*σ* = 0.5).

#### B-spline FFD (SimpleITK)

A 24 *×* 24 *×* 13 control-point grid was optimised with L-BFGS-B (tol= 10^−5^, 100 iterations, 5 corrections, 1000 evaluations; cost-function convergence factor = 10^7^). Correlation similarity and linear interpolation were used; the final transform was converted to a dense displacement field.

#### Cellular segmentation

The registered data are segmented on a cellular scale using Voluseg, a pipeline that implements locally constrained non-negative matrix factorization (NMF) to transform neighboring correlated pixels into functional segments^48^ (https://github.com/mikarubi/voluseg/). In brief, an intensity-based brain mask is created and divides the volume into spatially contiguous three-dimensional blocks, which overlap slightly to capture cells on the borders, and cell detection is run in parallel on these through the use of Spark, a distributed cluster-computing framework^153^. The algorithm fits the following model:

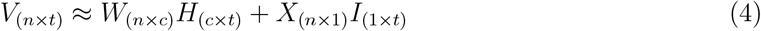

where *V* is the full spatiotemporal fluorescence matrix for each block, *W* and *H* are, respectively, the spatial location and time series of segmented cells, and *X* and *I* are rank-1 spatiotemporal models of the background signal. The timeseries are then normalized (mean-subtracted, and divided by the standard deviation of the timeseries). This resultant normalized timeseries is denoted F_*norm*_

#### Denoising

Synchronous Ca^2+^ bursts arising from muscle activation result in significant fluorescence. Camera pixels that ought only receive light from non-muscle cells can, if situated in close proximity to muscle tissue, capture some scattered photons from muscle cells, which confounds downstream analysis. While the contribution of a constant baseline fluorescence from muscle is effectively mitigated by mean subtraction of the data, fluctuations in such fluorescence remain problematic.

To address this challenge, the timepoints of muscle activity are located and nearby highly correlated cells identified. These potentially contaminated data points are masked, and linear interpolation is applied to the time series. This method may omit activity genuinely linked to muscle activations but is a conservative approach that could be refined in future work, or two-photon microscopy can be used instead of one-photon confocal to avoid out-of-point excitation. More specifically, the functional tissue compartments (FTC) corresponding to muscle are labeled, separating (owing to per-plane variations in contamination) and averaging them by imaging plane to form ‘muscle group’ timeseries. Muscle activation timepoints are determined using the probabilistic oasis model from Suite2p^154^ with parameters (window = maxmin, win baseline = 120s, sig baseline = 2). Activation timepoints were almost always isolated, i.e. neighboring timepoints contained no muscle activation by virtue of swimming being sparse in time in these experiments. Activation events with a probability over 0.6 are deemed real, and correlated cells above 0.35 close to the muscle group (within one plane above or below the muscle group) are masked at these timepoints, and the signal is then linearly interpolated between neighboring time points.

### Resolvable frequency estimation via Fourier Ring Correlation (FRC)

To quantify the resolvable spatial frequencies across the sample, we computed the Fourier Ring Correlation (FRC) for imaged volumes.

The FRC is the real-valued, normalized cross-power:

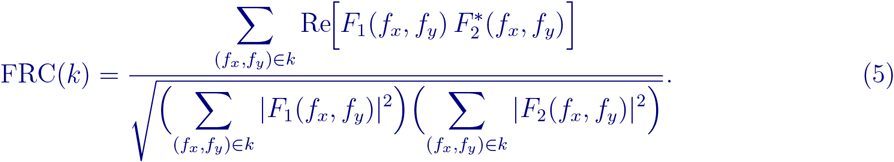

Here *k* denotes a radial shell in Fourier space, and *F*_1_(*f*_*x*_, *f*_*y*_) and *F*_2_(*f*_*x*_, *f*_*y*_) are the complex Fourier coefficients of two independent images. Writing each coefficient in polar form 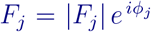 and using Re 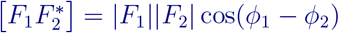, we obtain:

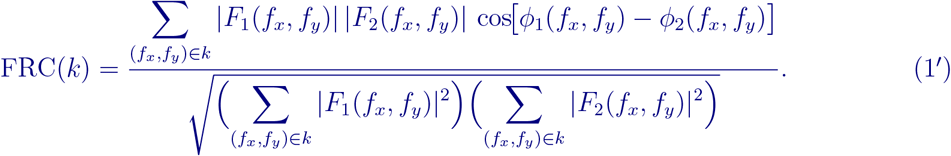

This expression highlights that the FRC is the weighted mean cosine of the phase differences in shell *k*, with weights given by the product of Fourier magnitudes, normalized so that −1 ≤ FRC ≤ 1. FRC maps were computed for volumes acquired dorsally and sagittally at 0.3 *×* 0.3 *×* 1 *µ*m resolution. To ensure that the results were not limited by the true spatial frequency content of the specimen, we used a transgenic line labeling all cellular membranes, guaranteeing high-frequency content throughout the sample. The analysis used a patch size of 20 *µ*m with 75% overlap. The spatial frequency at which the FRC fell below the standard 1/7 threshold was taken as the resolution cut-off, following established practice^155^, and was mapped across the sample.

### Coherence-based spectral clustering for identification of Functional Tissue Compartments

To cluster the data, Spectral Clustering^49,156^ was performed using a new coherence-based distance measure to define the adjacency graph. The similarity function was defined to be: 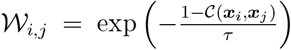 where 𝒞(***x***_*j*_, ***x***_*j*_)(*ω*) is the coherence between unit ***x***_*i*_ and ***x***_*j*_. This can be demonstrated to be a valid semi-positive definite kernel. Coherence is estimated using Welch’s method, that is interpreting it as the windowed magnitude-squared coherence estimator for stationary signals:

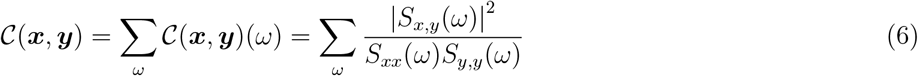

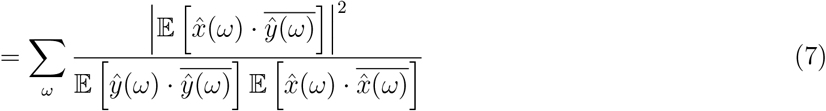

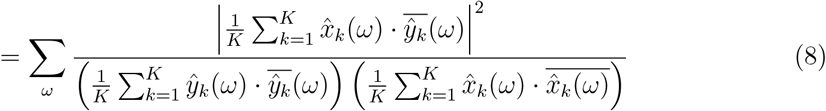

where *K* is the number of time windows.

This definition of coherence weights each 𝒞(*ω*) at each frequency irrespective of the power in that frequency band, this can be detrimental as bands of low power can be dominated by noise. To avoid this, the measure is normalized by the total power across all frequency bands:

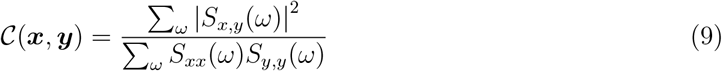

which remains bounded between 0 and 1. Having defined this adjacency graph, standard Spectral Clustering is performed. The Laplacian of the adjacency matrix is computed: *L*_norm_ = *I* − *D*^−1^*W* where *D* is a diagonal matrix and *d*_*i,i*_ = ∑_*j*_ *W*_*i,j*_. The lowest N eigenvectors of the matrix are computed and used as an encoding basis, and finally runs the k-means algorithm searching for N clusters. The k-means clusters are initialized with k-means++^157^, an algorithm for choosing good initializations of the cluster centroids. The model was fit with: *N*_clusters_ = 400, *τ* = 0.3, for coherence estimation, a time window of ∼ 8 minutes was used, with 80% overlap and a Hanning tapering window.

### Hierarchical clustering and lag-regression model

Spectral clusters mapping to muscle were identified manually based on their anatomy and timeseries and the mean activity of each cluster was computed. Hierarchical clustering was performed on the cluster means using the linkage function from the Python SciPy package, utilizing correlation as a metric and Ward linkage. A dendrogram was plotted, with muscle groups with greater than 0.5 correlation color-coded in similar shades, which we denote hyper-clusters, and shown in anatomical space with the same color scheme. To confirm the greater synchrony of the cervical-epaxial muscle with the ventral abdominal muscle, rather than with the neighboring hypaxial muscle, the cluster corresponding to the cervical-epaxial muscle was identified in different samples and cross-correlation between the activity of hypaxial and abdominal muscle computed, and a one-sided Wilcoxon signed-rank test used.

The mean activity of the muscle hyper-clusters identified in hierarchical cluster analysis was used as the basis for the regressors in the lag-regression model. A lag-regression model is a predictive model for time-series data in which a regression equation is used to predict the current value of the dependent variable, *y*(*t*), based on on both the current and past (i.e.: lagged) values of an explanatory variable, *x*(*t*), *x*(*t* − 1), …, *x*(*t* − *L*) where *L* is the maximum number of lags. This is the equivalent to assuming that the observed dependent variable arises from the convolution of the independent variable with a learnt kernel, which is parametrized by a set of weights, one for each lag:

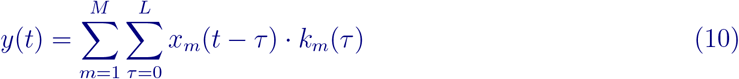

where *M* is the number of regressors/independent variables and *L* is the maximum number of lags.

The mean activity of the muscle hyper-clusters were converted into a weighted binary trace by identifying the onset of motor contraction and weighting them by the power of the contraction. Contraction onset and offset were identified by finding positive deflections above the noise floor and the time at which the curve returned to the noise floor; power was defined by the area under the curve of these two time points. For each cell *y*_*i*_ ∈ *Y*, a lag regression model is fit, 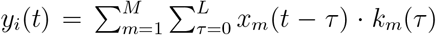 where *M* is the number of muscle regressors included (4 to 6), *L* the number of time lags included (60 seconds), *y*_*i*_(*t*) and *x*_*m*_(*t*) is the activity of a cell *i* and muscle regressor *m* at time *t*. The model is fit by finding the least squares solution using the numpy.linalg.solve function. The identified kernel values are 3-fold cross-validated and the average *R*^2^ value on held-out data is reported.

### Kymograph analysis

Peak oscillation frequency was defined as the frequency with largest power excluding the zero-frequency power. To compute kymographs of ependymal cell activity from imaging data acquired from *Tg(foxj1a:GCaMP7f)* imaging, the ventral and dorsal boundaries of the brain were first anatomically identified. Line integrals perpendicular to the ventral boundary were computed, resulting in a 1*×*N vector with N being the number of spatial bins used. When repeated over frames, this results in a N*×*T matrix. The data were denoised through spatial smoothing using a Gaussian kernel with a standard deviation of 4 pixels and temporally filtered with a bandpass Butterworth filter of order two, employing cutoff periodicities of 0.5 to 9 minutes.

### Coherence k-means algorithm

Derivation of the algorithm used in Fig. 3k. The k-means algorithm alternates between computing cluster centroids and identifying the clusters to which data points belong. We aim to use coherence as a measure of similarity. We define the following: *M* : # of latent clusters, *K*: # of sub-samples used to average over to calculate coherence, *T* : # of total time points, *L*: # of time points per sub-sample, ***x***_*i*_ ∈ *R*^*T*^ : sample *i*, ***x***_*ik*_ ∈ *R*^*L*^: sub-sample *k* of sample *i* of length *L*, ***µ***_*m*_ ∈ ℝ^*L*^: centroid of cluster *m, 𝒞*(*x, y*)(*ω*): coherence between *x* and *y* at frequency *ω*.

We wish to find a centroid ***µ***_*m*_ such that the sum of the coherences of points within that cluster ∑_*i*_ 𝒞(***x***_*i*_, ***µ***_*m*_) is maximal. This quantity is invariant to an arbitrary real scaling of 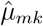, so the following constraint is added: 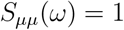, i.e.: 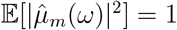 that is 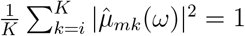.

Thus, we aim to maximize:

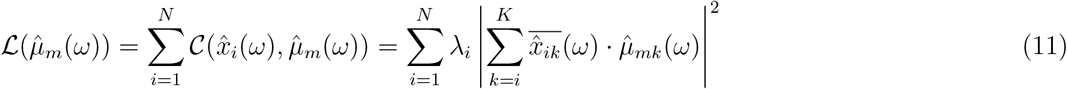

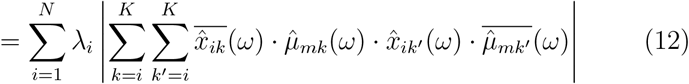

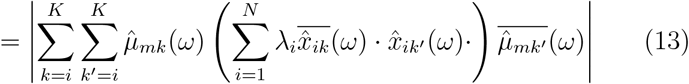

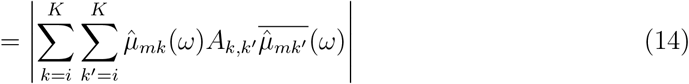

writing 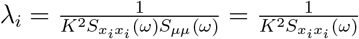

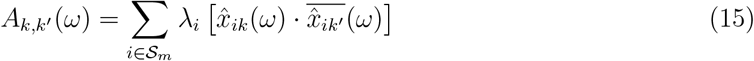

We note that 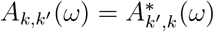 so the elements form a Hermitian matrix *A*(*ω*).

Vectorizing *µ*_*m*_, the objective can be written to maximize as:

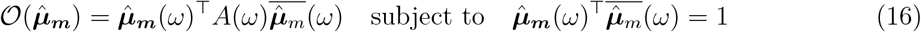

from which it follows that 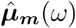 is the leading eigenvector of *A*(*ω*).

To initialize the group labels, k-means++^157^ is first run, an algorithm for choosing the initial values. Cluster centroids are calculated as above and cluster allocation reassigns the cluster ID based on coherence. The process is iterated until the convergence of the labels.

### Characterizing hypoxia-induced physiological changes

#### Estimation of blood vessel diameter

To measure blood vessel diameter, a transgenic line labeling vascular endothelium (*Tg(flk1:dsRED-CAAX)*) was imaged. To ensure accurate width estimation, volumetric stacks were acquired with 2 µm z-spacing spanning the entire width of the mesenteric blood vessel and maximum intensity projection of each stack was computed. A line crossing the middle of the vessel orthogonally is defined and the kymograph computed using Fiji’s kymograph function resulting in a matrix with T rows, where T is the number of time points. To denoise the data, the matrix was row-wise median filtered with window size of 5 pixels. The matrix was thresholded at the midrange value. The largest connected component is identified using the scipy-ndimage label function, which corresponds to the main blood vessel. The width of the mask at each row was computed, and the resulting time series was smoothed over time using a rolling mean with a window length of 10 time points. Finally, the width was scaled by the resolution of the data to have units of microns. Baseline and hypoxic-state vessel widths were defined as the mean width over the one-minute interval preceding the onset of the gas switch or at steady-state hypoxia (10 min following the gas switch from 21% to 10% O_2_) respectively.

#### Estimation of blood flow

To measure blood flow, a transgenic line labeling red blood cells (*Tg(gata1:dsRED)*) was imaged. For faster imaging, a single plane was acquired at 10 Hz. The plane was selected to cover the mesenteric artery, which runs parallel to the body. The frame rate was too low to track individual red blood cells; instead, local bulk blood flow was estimated by quantifying changes in fluorescence induced by red blood cells moving in and out of any specific ROI. ROIs over arteries feeding the brain, muscle, and mesentery were manually defined. The absolute value temporal difference of the mean was computed for each of these ROIs. As passing red blood cells result in differing changes in magnitude depending on whether the entire or a fraction of the cell was in the ROI, the timeseries was clipped between 0 and 3*×* the median value of the beginning of the timeseries (normoxic period, 2 min). The timeseries was finally normalized by dividing by the standard deviation of this baseline period. The baseline and hypoxic-state blood flow were defined as the mean blood flow over the one minute interval preceding the onset of the gas switch or at steady state hypoxia (10 min following the gas switch from 21% to 10% O_2_) respectively. Change in blood flow was defined as the difference between these values and baseline blood flow (mean blood flow prior to gas switch, ∼2 min). To compare the effects of optogenetic inhibition of the hindbrain and control illumination of the heart, mean blood flow over each 1 min stimulation period was computed for each condition (hindbrain or control). Change in blood flow was defined as the difference between these values and baseline blood flow (mean blood flow prior to gas switch, ∼2 min).

#### Baseline oxygen modulation score

To quantify the effect of hypoxia on baseline calcium levels, activity traces were passed through a min-max filter using centered windows of length 3 min, and were subsequently smoothed using a Gaussian kernel with a standard deviation of 15 sec. The mean baseline activity during normoxia and hypoxia over a 3 min window (taken at the end of the normoxic/hypoxic phases to ensure steady-state oxygen levels had been reached) was computed on a per-cell basis and subtracted.

### Whole-Body Expansion Microscopy

A comprehensive, step-by-step protocol is provided in the Supplementary notes **??**.

### Whole-Body Expansion Microscopy combined with immunofluorescence (WB-ExM-IF)

#### Fixation, IF and agarose-embedding

Whole larval zebrafish were fixed with 4% PFA overnight at 4°C on a shaker, then washed with 1*×* PBS 4x 15 min. Fixed fish were permeabilized for 5 hr with 0.5% Triton X-100 in 1*×* PBS (PBST-0.5) at room temperature (RT) with gentle agitation. Permeabilization must be included for all specimens, whether they will be stained with antibodies or not. For IF, specimens were blocked for 3 hr with blocking buffer (5% goat serum, 0.5% Triton X-100, 0.1% Na Azide in 1*×*PBS) at RT with gentle agitation. Blocked specimens were incubated with primary antibodies diluted 1:100 in blocking buffer for 3 days at RT, followed by washing with PBST-0.5 3*×* 2 hr. Specimens were then incubated with secondary antibodies diluted 1:100 in blocking buffer for 2 days at RT, followed by washing with PBST-0.5 3*×* 2 hr. Samples were embedded in a thin layer of 1% low-melting temperature agarose to ensure desired sample orientation. Antibodies used: rabbit anti-eGFP (Invitrogen, A11122), chicken anti-RFP (synaptic systems, 409006), goat anti-rabbit Atto647N (Sigma, 40839), donkey anti-chicken Alexa568 (Invitrogen, A78950).

#### Protein anchoring

Specimens were incubated in Acryloyl-X SE at 20 ug/mL in 1*×* PBS (anchoring solution) for 1 hr at RT followed by washing in 1*×* PBS. Anchoring solution is prepared just before use from a 10 mg/mL stock solution dissolved in anhydrous DMSO. Samples were washed in 1*×* PBS 3*×* 5 min.

#### Gelation and digestion

Specimens were incubated in the first gelation solution (10% Acrylamide, 0.5 M sodium acrylate, 0.1% bis-acrylamide, 0.01% 4HT, 0.2% TEMED and 0.2% APS, 1*×* PBS) on ice 3*×* 10 min with gentle agitation. Gelation chambers were constructed using an uncharged glass slide as the bottom piece and side walls consisting of 11 layers of Scotch tape (∼ 600 µm thick to approximate the thickness of the agarose block) serving as spacers. Coverslips were placed on top of the chamber. The chambers were filled with first gelation solution by pipetting in solution from the open side. Fully assembled chambers were carefully placed in a humidified incubator for 2 hr at 37°C for gelation. After gelation, the chambers were carefully disassembled and extra gel was trimmed around the samples using a scalpel, leaving a ∼ 2 mm margin around the fish. Gelled specimens were treated with disruption buffer (DB; 5% SDS, 50 mM Tris pH 7.5, 200 mM NaCl in H2O) at 100°C overnight in Eppendorf tubes, then washed with 1*×* PBS 3*×* 20 min.

#### Re-embedding and staining

Gelled and disrupted specimens were incubated in the second monomer solution (10% Acrylamide, 0.5 M sodium acrylate, 0.02% bis-acrylamide, 0.01% 4HT, 0.2% TEMED and 0.2% APS, 1*×* PBS; this is the same as the first gelation solution except with bisacrylamide reduced from 0.1% to 0.02%) on ice for 3*×* 10 min with gentle agitation. Gels imbued with second gelation solution were placed on uncharged glass slides with side spacers as before composed this time of 20 layers of Scotch tape (∼ 1.2 mm). A coverslip was positioned on top as the chamber top piece. The space surrounding the first gel was filled with additional second gelation solution. Fully assembled chambers were carefully placed in a humidified incubator for 2 hr at 37ºC. After gelation, the chambers were gently disassembled, and excess gel was trimmed around the embedded fish with a scalpel, leaving a ∼ 1 mm margin. Gelled specimens were stained with Alexa 488-NHS ester (Thermo Fisher Scientific, A20000) and/or Atto647N-maleimide (AAT Bioquest, 2857) dyes. Samples were incubated in dye solution (1:1,000 in PBS from 10 mg/ml stocks dissolved in anhydrous DMSO (Thermo Fisher, D12345)) for 1 hr at RT with shaking, then washed 3*×* 1 hr with 1*×* PBS. The re-embedding process can be repeated up to four times resulting in ∼5 *×* expansion.

#### Imaging, tile stitching and data registration

Samples were mounted onto a glass slide using polylysine and attached to a custom built sample holder. Specimens were imaged on a Zeiss Z1 light-sheet microscope (20x NA 1.0 water immersion objective), immersed in 1*×* PBS. Stitching and registration were done using the Imaris stitching and registration software using the default parameters.

#### Online data

An example dataset can be viewed online at https://neuroglancer-demo.appspot.com/#!gs://flyem-user-links/short/2025-01-03.165221.071748.json. This is the sample shown in Fig. 8b : double transgenic zebrafish animal labeling the ventricular and vascular systems (*Tg(foxj1a:eGFP)* x *Tg(flk1:dsRED-CAAX)*), stained against eGFP (magenta) and dsRED (green) (10 days post-fertilization, expanded ∼2×).

### Whole-Body Expansion Microscopy combined with in situ hybridization (WB-ExM-FISH)

All solutions were made using molecular grade (RNase-free) water and reagents. A comprehensive, step-by-step protocol is provided in the Supplementary notes **??**.

#### Anchoring stock solutions

Melphalan and Acryloyl-X SE (AcX) were prepared as 2.5 mg/mL and 10 mg/mL stocks, respectively, in anhydrous DMSO and stored desiccated. Melphalan-X was prepared by combining melphalan and AcX stocks at a ratio of 4:1 to yield a final concentration of 2 mg/mL each and incubating at RT overnight with shaking. Melphalan-X was aliquoted and stored desiccated.

#### Fixation and permeabilization

Whole larval zebrafish were fixed with 4% PFA overnight at 4°C on a shaker, then washed with PBS 4*×* 15 min in 1*×* PBS. Fixed fish were permeabilized for 1 hr with 0.5% Triton X-100 in 1*×* PBS (PBST-0.5) at RT with gentle agitation.

#### RNA and protein anchoring

Fixed and permeabilized fish were washed with MOPS buffer (20 mM, pH 7.7) for 30 min at RT. Specimens were next treated with 1 mg/mL Melphalan-X supplemented with 0.1 mg/mL extra AcX diluted freshly into MOPS buffer overnight at 37°C, followed by washing in MOPS buffer (2*×* 5 min) and then PBS (2*×* 5 min). Anchored specimens were then mounted on poly-L-lysine-coated coverslips with the fish positioned on its side.

#### Gelation and digestion

Mounted specimens were incubated in the first gelation solution and gelled as in the protein method variant above. Briefly, this includes incubating in complete gelation solution, assembling the gelation chamber, gelling at 37°C, disassembling the chamber and trimming the gel to leave ∼ 2 mm margin around the specimen. The trimmed gel was then incubated in the first digestion solution (500 mM NaCl, 0.3% SDS, 50 mM Tris-HCl pH 8.0, 1 mM EDTA with Proteinase K diluted 1:50 from 800 U/mL stock) for 4 hr at 50°C, followed by incubation in the second digestion solution (50 mM NaCl, 1% SDS, 50 mM Tris-HCl pH 8.0, 1 mM EDTA with Proteinase K diluted 1:50 from 800 U/mL stock) overnight at 50°C. Digested specimens were washed 4*×* 15 min in 1*×* PBS.

#### Re-embedding

Digested specimens were re-gelled as in the protein method above. Gelled specimens were recovered into 1*×* PBS and trimmed, leaving ∼ 1 mm margin around the specimen.

#### Probe Hybridization and Hybridization Chain Reaction (HCR)

Gels were incubated in hybridization buffer (Molecular Instruments, https://www.molecularinstruments.com/hcr-rnafish-products) for 30 min at 37°C, followed by incubation in primary probes designed by Molecular Instruments (6 µl in 600 µl hybridization buffer, 10 nM final concentration) overnight at 37°C. Following hybridization, gels were washed with pre-warmed (37°C) probe wash buffer (Molecular Instruments) 3*×* 30 min, then washed with pre-warmed (37°C) 1*×* PBS 3*×* 1 hr, and once overnight at RT. Gels were next incubated in amplification buffer (Molecular Instruments, https://www.molecularinstruments.com/hcr-rnafish-products) for at least 30 min at RT. Hairpins were diluted 1:50 in amplification buffer and snap cooled by heating to 95°C for 90 s followed by cooling at RT for 30 min. Gels were incubated for 4 hr at RT in the dark in hairpin/amplification buffer mix. Amplified gels were washed in 5x SSCT (5x SSC, 0.1% Tween) 2*×* 20min at RT, then in 0.5x SSCT (0.5x SSC, 0.1% Tween) 2*×* 40min at RT and finally equilibrated in 1*×* PBS for 2*×* expansion. Probes used: phox2bb_*B*_3 (lot # RTG113), Th_*B*_1 (lot # RTB474), hsd31_*B*_5 (lot # RTG103), calca_*B*_2 (lot # RTG105), pomca_*B*_5 (lot # RTG108).

#### Stripping and re-probing

For multi-round imaging, HCR amplification products and probes were stripped by digestion with DNase followed by re-probing and imaging as described for the first round, above.

#### Imaging, tile stitching and data registration

Samples were processed the same way as WB-ExM-IF samples.

### PhotoMap

To quantify deformation introduced by expansion in an unbiased manner, we developed PhotoMap, a method that measures the gel’s deformation field from a pre-imposed reference pattern. A regular grid is photobleached into the sample prior to expansion and re-imaged afterwards, enabling straightforward visualization and quantification of spatial distortions. A comprehensive step-by-step protocol is provided in the Supplementary Notes (**??**).

#### Fluorescent gel formation

A fluorescent monomer was created by conjugating AF488-NHS (20 mg/ml, 31 mM) to 3-aminopropyl methacrylamide (10 mg/ml, 56 mM) in the presence of triethylamine (10%, 720 mM). The reagents the incubated overnight at room temperature, protected from light. Gel preparation followed the protocol described in **??**, except that 1:100 of the fluorescent monomer was added to the first monomer solution to render the gel intrinsically fluorescent. In brief, samples were fixed, agarose-mounted, chemically anchored, and polymerized, and then kept unexpanded in the gelation chamber for photobleaching.

#### Photomap grid formation through photobleaching

Using a large field of view 2 photon microscope, a 50 *×* 50 *×* 50 µm grid was photobleached into the gel prior to expansion. Custom code was written to drive the microscope via ScanImage^**?**^ . Regions corresponding to the eyes, which contain light-absorbing pigments, were excluded from the photobleaching pattern to prevent excessive local heating. After photobleaching, the sample was disrupted overnight, expanded, and re-imaged using the same microscopee at 1 *×* 1 *×* 5 µm resolution.

#### Photomap deformation field analysis

To extract the resulting grid intersections from post-expansion images, we used a two-step procedure combining zero-mean normalized cross-correlation (ZNCC) with peak detection.

#### Template matching via zero-mean normalized cross-correlation (ZNCC)

We first manually selected a representative intersection from the raw image and used this as a template *T* ∈ ℝ^*P ×Q*^. The full image *I* ∈ ℝ^*N ×M*^ was then scanned using ZNCC, defined as:

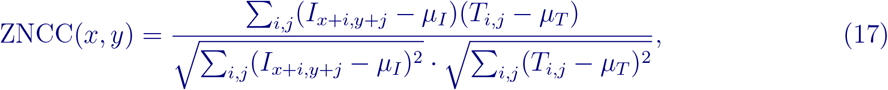

where *µ*_*I*_ and *µ*_*T*_ are the local mean intensities of the image and the template, respectively, over the region of interest. This was efficiently computed via FFT-based cross-correlation and using sliding-windows. To enhance local contrast and robustness to drift in the global dynamic range, the ZNCC map was Z-score normalized over a *k × k* neighborhood.

#### Local peak detection

The resulting ZNCC map was passed through a local top-percentile filter, keeping the brightest 10% of pixels per patch (80 *×* 80 px). A Laplacian-of-Gaussian (LoG) filter was then applied to enhance bright-on-dark blobs corresponding to grid intersections. Peak centers were finally extracted using *peak_local_max* function from *scikit-image*, with minimum distance (50 px) and relative threshold parameters empirically tuned to match the grid geometry.

#### Quantifying local grid distortion

To assess spatial distortions in the photobleached grid post-expansion, two geometric features were computed for each grid point:

- **Length deviation:** The mean relative deviation of edge lengths (horizontal and vertical) from an ideal spacing *L* = 90 pixels, computed as 1 − ||vec (*v*) |/*L*|.
- **Angle deviation:** The cosine of the angle between adjacent horizontal and vertical vectors at each point, ideally zero for orthogonal axes (i.e., deviation from 90°).

These values were computed across the entire sample. To visualize and compare the spatial distortion statistics, kernel density estimates (KDEs) of each component (length and angle) were computed separately for on-sample and off-sample points.

### ExM data modeling and quantification

#### Building 3D ExM model

To build the main whole-fish model, WB-ExM-Histo zebrafish datasets were acquired (total protein stain Alexa488-NHS) and imaged on a Zeiss Z1 light-sheet microscope (20x NA 1.0 water immersion objective). The organs were manually segmented within the Amira software and the meshes imported in the software Blender. To incorporate cellular populations that have too complex morphology or spatial distribution for manual annotation, and that can be molecularly defined, such as ependymal cells and the vasculature, WB-ExM-IF data was utilized. To build meshes, WB-ExM-IF data was up-sampled in z by a factor of 2, converted to 8 bit format, denoised (Fiji’s Remove Outliers function), thresholded (Fiji’s Threshold function, Otsu algorithm), opened in Fiji’s 3D viewer as a surface and exported as an stl binary file, ready to be imported into Blender.

#### Quantification of SNR in ExM data

Scattering across the sample was quantified by comparing SNR within muscle fibers between the near and far-side of the sample, defined as the average difference between minimum and maximum intensity signals across muscle sarcomeres.

#### Quantification of enteric motor vagus innervation

To estimate motor vagus innervation density along the gastrointestinal tract, an animal expressing RFP under the Islet-1 promoter (*Tg(isl1CREST-hsp70l:mRFP)*) were stained and expanded using WB-ExM-IF. The gastrointestinal tract was manually segmented using the Amira software. The IF signal was averaged radially resulting in a 1-d histogram of average innervation density as a function of position along the gastrointestinal tract.

#### Tracing of motor nerves

A high-resolution confocal stack of a 4 days post fertilization (*Tg(VAChTa:Gal4;UAS:CoChR-eGFP)*) was acquired using a spinning-disk confocal microscope (0.1625 µm x 0.1625 µm x 1 µm voxel size). Nerve bundles and muscle outlines were traced manually using Amira, and the resulting tracts were imported into Blender.

#### EM tracing of ependymal cells

Ependymal cells were identified are readily identifiable within the EM volume^98^ based on their distinctive location and morphology as visualized in the WB-ExM-IF data. The published dataset includes both raw data and the segmentation of cells into subcellular fragments. Ependymal cells were traced by agglomerating cellular fragments using the Neuroglancer web interface. Exemplar commissural axons were also traced, as well as the neighboring vasculature.

## Supporting information

Supplemental data and protocols

## Code and data availability

The code for registration is available on GitHub in python and MATLAB at https://github.com/vruetten/wholistic_registration and https://github.com/Weizheng96/WHOLISTIC-registration, respectively, and segmentation code is available at https://github.com/mikarubi/voluseg/. The 3D geometric body model is available on GitHub https://github.com/vruetten/wholebodymodel. An example WB-ExM-IHC dataset can be viewed online at https://neuroglancer-demo.appspot.com/#!gs://flyem-user-links/short/WHOLISTIC_ExM_fox1ja_flk1_10dpf.json. The raw data is available at https://s3.janelia.org/ahrens-lab/ruettenv/ExM/. Raw WHOLISTIC data will be made available at https://s3.janelia.org/ahrens-lab/ruettenv/WHOLISTIC/. The plasmid map for *Tg(ubb*^*R*^*:jGCaMP8m)* can be accessed on Addgene (# 232485).

The paper has an accompanying website on which the data can be browsed: https://wholistic.janelia.org/.

## Acknowledgments

We are grateful for the valuable support and contributions from many individuals. For computational assistance, we thank Talley Lambert for adapting his ND2 Python package and Greg Fleishman for advice on registration. For optical guidance, we are grateful to the Janelia’s light-microscopy core team, especially Michael DeSantis, and Janelia’s Experimentation and Technology team and Janelia’s Scientific Computing Team, especially Jody Clements and Konrad Rokicki. We thank Janelia’s Anibody Project Team for support and discussions in building the 3D anatomical model. We are grateful to Janelia’s vivarium staff, especially Jared Rouchard and his team. We thank the Janelia Visiting Scientist Program for their support. We thank Vivek Jayaraman, David Prober, Ron Vale, Kristian Herrera, Ken Harris, Erik Snapp, Boaz Mohar, Isabel Espinosa-Medina, Helen Farrants, Rylan Schaeffer, Rob Johnson, Anoj Ilanges, Yin Liu, Emmanuel Marquez Legorreta, Dhruv Zocchi, Arco Bast, Pavlo Bulanchuk, Cyrus Mostajeran, John Rallis, Alex Chen, Will Dorrell, Monika Makurath, Alexandre Benedetto, and Dan Cortes for discussions and for feedback on the manuscript. We thank Isabel Espinosa-Medina and Sujatha Narayan who provided guidance on transgenic methods. We thank the Ahrens and Sahani lab for ongoing feedback. This research was supported by the Howard Hughes Medical Institute (MA), the Gatsby Charitable Foundation (MS), NIH grant RF1MH125933 (MR), NSF grant 2207891 (MR).

## Author contributions

Conceptualization: VMSR, MBA and MS. Methodology, data collection and data analysis: VMSR. Registration and segmentation: WZ, VMSR, YC, MR and GY. Building 3D model and rendering: IS. Expansion microscopy: VMSR, ME, AH, YH, KC, GI, AP, AD and PT. Genetics: VMSR, CG, ALL, CS, MK, MR, BJ, YW, PJK. Conceptualizing and testing of hypoxia findings: VMSR, SLG, FE, MCF. Writing, review and editing: VMSR, MBA and BM. Supervision and funding: MBA and MS.

