## Supplemental data and protocols for "Imaging cellular activity simultaneously across all organs of a vertebrate reveals body-wide circuits"

### Extended Data Figures

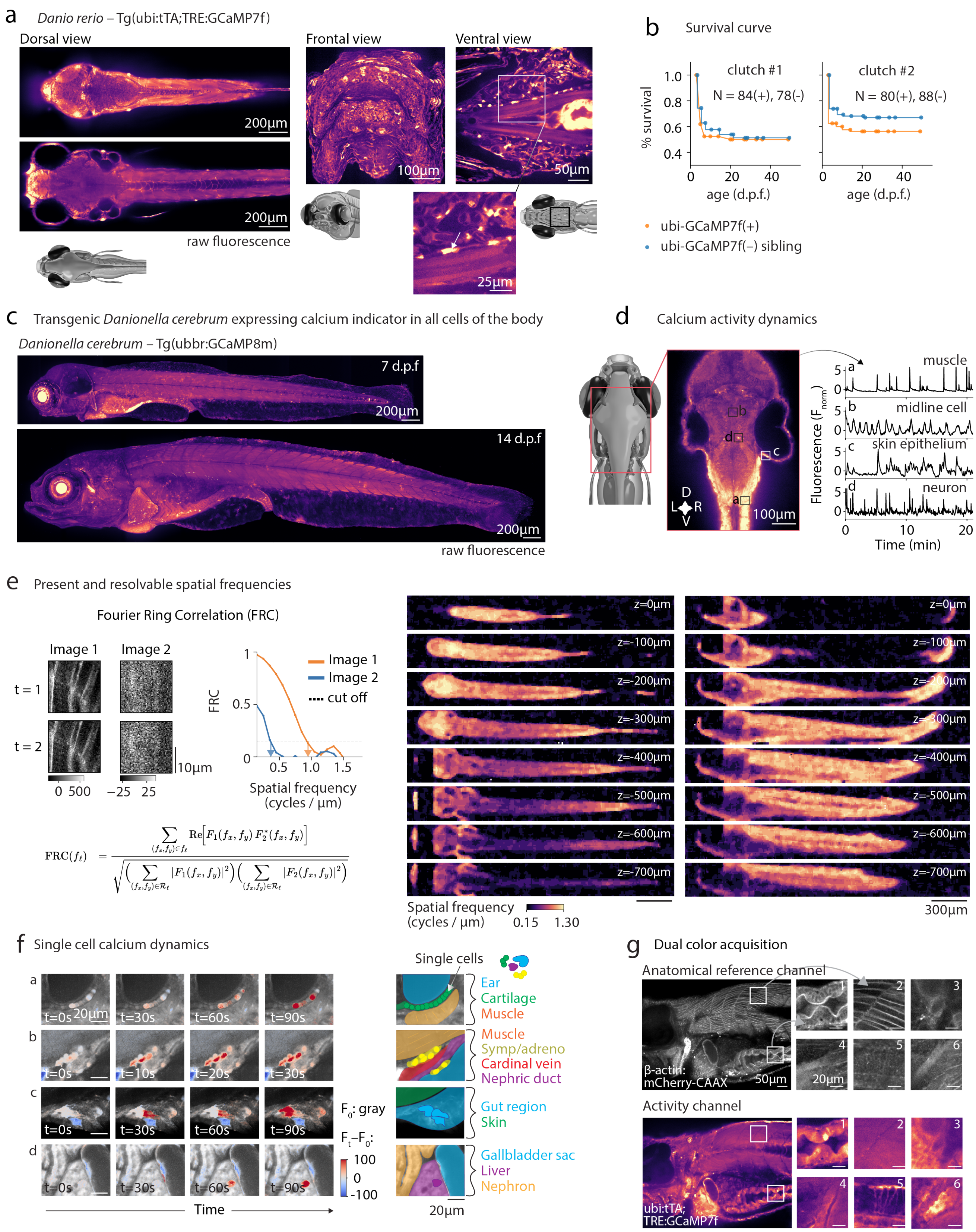

$$\mathrm{FRC}(f_\ell) = \frac{\sum_{(f_x,f_y)\in f_\ell} \mathrm{Re}\left[F_1(f_x,f_y)\,F_2^*(f_x,f_y)\right]}{\sqrt{\left(\sum_{(f_x,f_y)\in\mathcal{R}_\ell} |F_1(f_x,f_y)|^2\right)\left(\sum_{(f_x,f_y)\in\mathcal{R}_\ell} |F_2(f_x,f_y)|^2\right)}}$$

**Extended Data Fig. 1: WHOLISTIC transgenics, workflow and registration.**

#### Extended Data Fig. 1: WHOLISTIC transgenics and workflow.

**a**, Complementary views of WHOLISTIC zebrafish transgenic. *Left to right*: 1) dorsal view (two planes at different depths), 2) frontal view (maximum projection), 3) ventral views of gill regions, with enlarged view containing unidentified cell types.

**b**, Survival curves for zebrafish pancellular GCaMP7f transgenic line. Embryos were segregated into GCaMP positive embryos (N = 84 and 80, represented by the orange line) and GCaMP negative siblings (N = 78 and 88 indicated by the orange line).

**c**, WHOLISTIC for *Danionella* species. Pancellular GCaMP transgenic line. Sagittal view of *Tg(ubb*$^{R}$*:jGCaMP8m) Danionella cerebrum* (7 and 14 days post fertilization), imaged using spinning-disk confocal microscopy.

**d**, Calcium activity dynamics from *Danionella cerebrum* WHOLISTIC line. *Left*: dorsal view of brain. *Right*: example time-series from 1) neuropil, 2) skin epithelium, 3) hindbrain neuron, 4) muscle.

**e**, Present and resolvable frequencies across the sample. Left: schematic of the Fourier Ring Correlation (FRC) method for quantifying the reliably resolvable spatial frequencies within an image. Two independent images of the same object are acquired, their 2D Fourier transforms computed, and the phase correlation calculated, radially averaged, and normalized. The spatial frequency at which the correlation falls below 1/7 is taken as the resolution cut-off[1]. Right: spatial map of the highest spatial frequency at which the FRC remains above the 1/7 threshold. High spatial frequencies are detectable across most of the body, with localized reductions in regions between the ear and in deep pharyngeal areas between the gills.

**f**, Single-cell dynamics across various body regions. Sequential frames displaying fluorescence intensity variations (blue-red) superimposed on anatomical reference (gray). Specific images depict *top to bottom*: 1) the ear with surrounding active chondrocytes; 2) the midline cardinal vein surrounded by active cells; 3) the ventral fin containing large active skin epithelial cells; 4) the liver displaying active hepatic cells.

**g**, Volumetric tissue registration via dual-color imaging using double transgenic line (*Tg(ubi:tTA; TRE:GCaMP7f)*; *Tg(β-actin:mCherry-CAAX)*) line. *Top*: anatomical reference channel showing membrane labeling. *Bottom*: activity channel. Insets showing enlarged views of: (1) gut, (2) muscle, (3) gallbladder sphincter, (4) blood vessel, (5) spinal cord, and (6) neural tract.

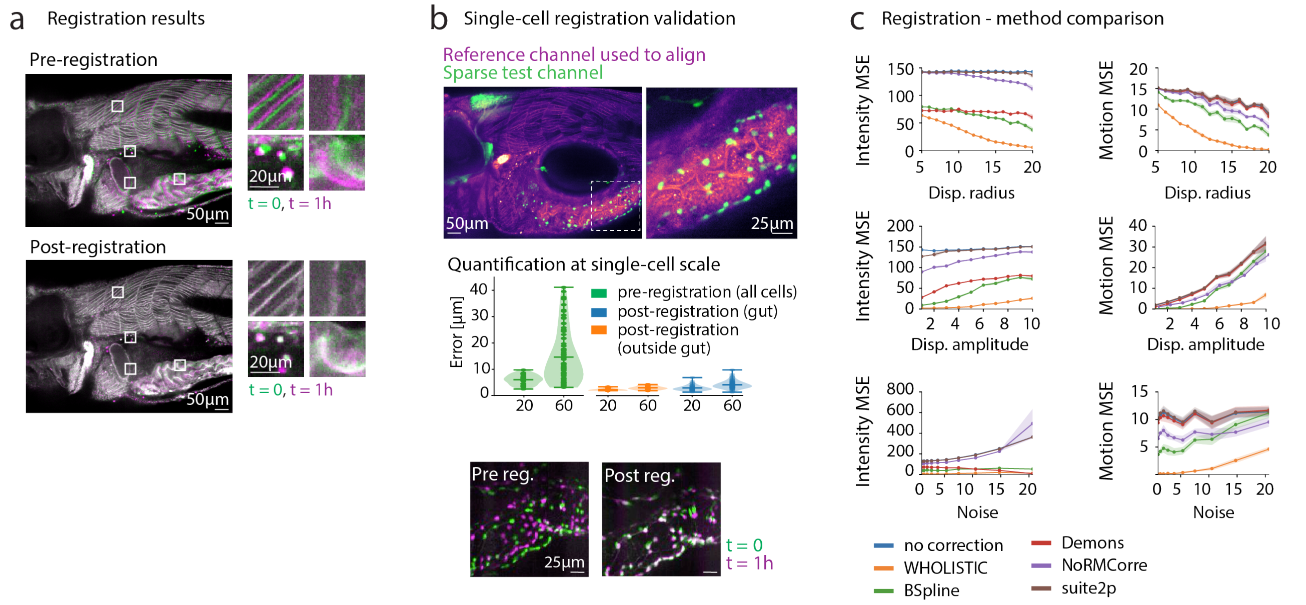

#### Extended Data Fig. 2: WHOLISTIC registration.

**a**, Registration results. Overlay of data from the start of an experiment (green) and 1 hour into recording (magenta). *Top left*: pre-registration overlay showing significant drift. *Bottom left*: post-registration overlay showing alignment correction in the same regions and at the same timepoints. *Right*: enlarged views of muscle, gallbladder, skin, and gut regions pre- and post-registration.

**b**, Validation of registration results. *Top left*: double transgenic line used for validation (*Tg(β-actin:mCherry-CAAX; phox2bb:eGFP)*) with inset showing enlarged view of the gut – the most challenging region to register. *Top right*: overlay of sparse cell line at the start of the experiment (green) and 1 hour in (magenta), pre-registration (left) and post-registration (right). *Bottom*: violin plots of cellular displacement after 20 and 60 minutes: pre-registration displacement (green), post-registration displacement in gut (blue), and in the rest of the viscera and brain (orange).

**c**, Registration method comparison. Synthetic datasets were generated by applying known motion fields to experimental images, varying displacement radius, amplitude, and noise to test robustness. Each dataset was registered with five methods: WHOLISTIC, B-spline, Demons, NoRMCorre, and Suite2p. Mean squared error (MSE) was computed for both the image intensity and the recovered motion field, enabling direct comparison to ground truth.

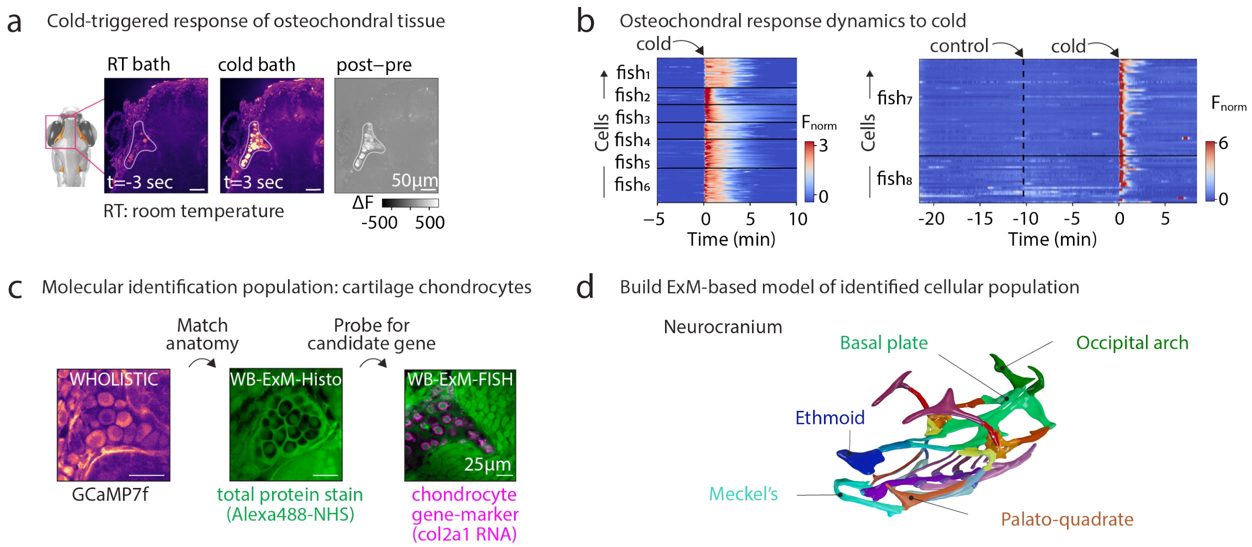

#### Extended Data Fig. 3: WHOLISTIC screening for cellular responses to stimuli.

**a**, WHOLISTIC screen for cold-responsive cells. Following a 5-minute period of baseline recording, 10°C water is introduced to the imaging chamber. Transverse section of the anterior portion of the cranium shown pre- and post-exposure to cold, and ΔF. Cold-responsive population outlined in gray.
**b**, Left: raster of $Ca^{2+}$ levels in osteochondral cells aligned to cold stimulus onset, showing time-locked responses in 6 animals. Right: control assay - raster of $Ca^{2+}$ levels in osteochondral cells showing little to no time-locked responses to control stimulus (room temperature water) in 2 animals and responses to cold stimulus.
**c**, Cell type identification and modeling via Whole-Body ExM. Matching cellular morphology between in vivo and ExM data. *Left to right*: 1) view of cold-responsive region in WHOLISTIC in vivo data, 2) corresponding region in WB-ExM-Histo data matched by location, cell morphology, and spatial distribution, using Alexa488-NHS (green), 3) corresponding region in WB-ExM-FISH sample stained against chondrocytic gene-marker, col2a1 (magenta).
**d**, Reconstruction of the neurocranium - 3D model of young zebrafish cartilage generated from ExM data, annotated based on literature.

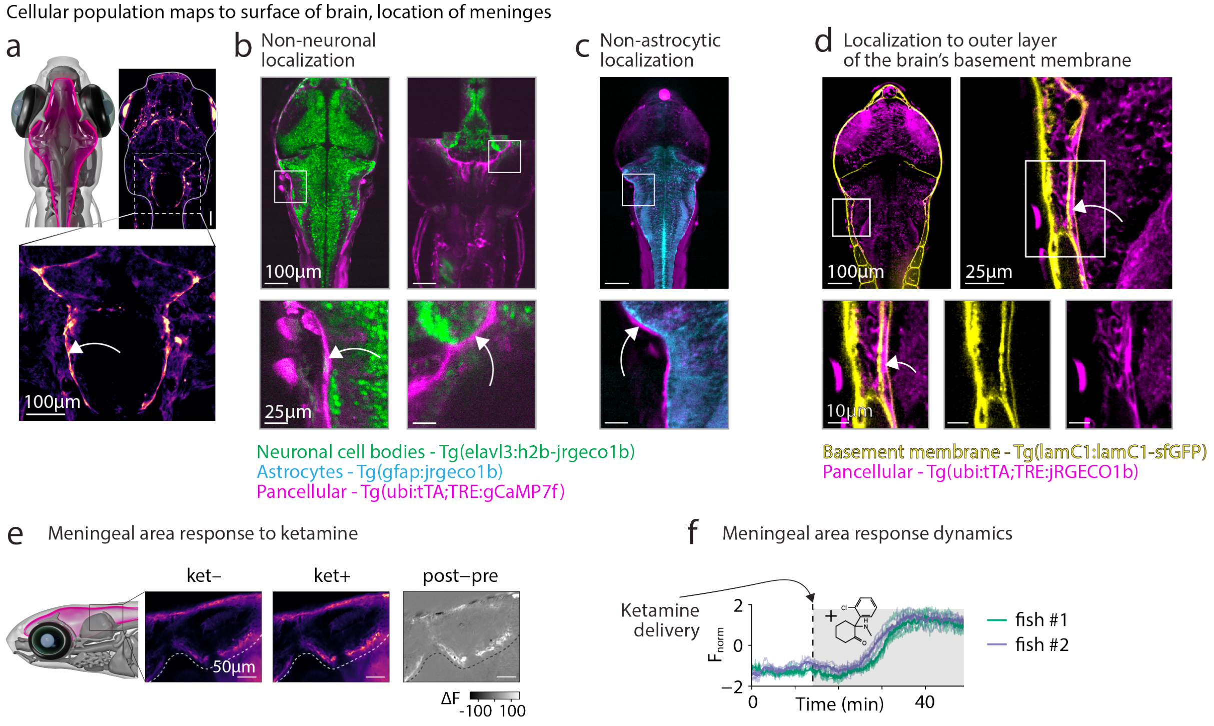

**Extended Data Fig. 4: WHOLISTIC screening for cellular responses to drugs.**
**a**, Maximum intensity projection showing a brightly fluorescent cellular layer surrounding the brain (top). Enlarged view of the hindbrain (bottom).
**b**, Border cells (visible among magenta cells) do not colocalize to neurons (green). Image of double transgenic line at dorsal (left) and ventral (right) planes (*Tg(elavl3:H2B-jRGECO1b; ubi:tTA;TRE:gCAMP7f)*) with inset showing enlarged view of border region. Contrast of magneta adjusted to highlight border cells.
**c**, Border cells (visible among magenta cells) do not colocalize to astrocytes (blue). Image of double transgenic line (*Tg(gfap:jRGECO1b; ubi:tTA;TRE:gCAMP7f)*) showing non-colocalization of border cells and astrocytes, with inset showing enlarged view of border region. Contrast of magenta channel adjusted to highlight border cells.
**d**, Border cells (visible among magenta cells) localize to the outer layer of the brain's basement membrane (yellow). Image of double transgenic line (*Tg(lamC1:lamC1-sfGFP; ubi:tTA;TRE:jREGCO1b)*) showing intercalation of border cell and brain basement membrane, with inset showing enlarged view of border region. Contrast of magenta channel adjusted to highlight border cells.
**e**, Meningeal area response to ketamine exposure. Animals were exposed to ketamine following a 25 min period of baseline imaging. *Top*: Sagittal section showing pre- and post-exposure to ketamine, and $\Delta$F.
**f**, Activity traces from meningeal regions; population average in darker shade.

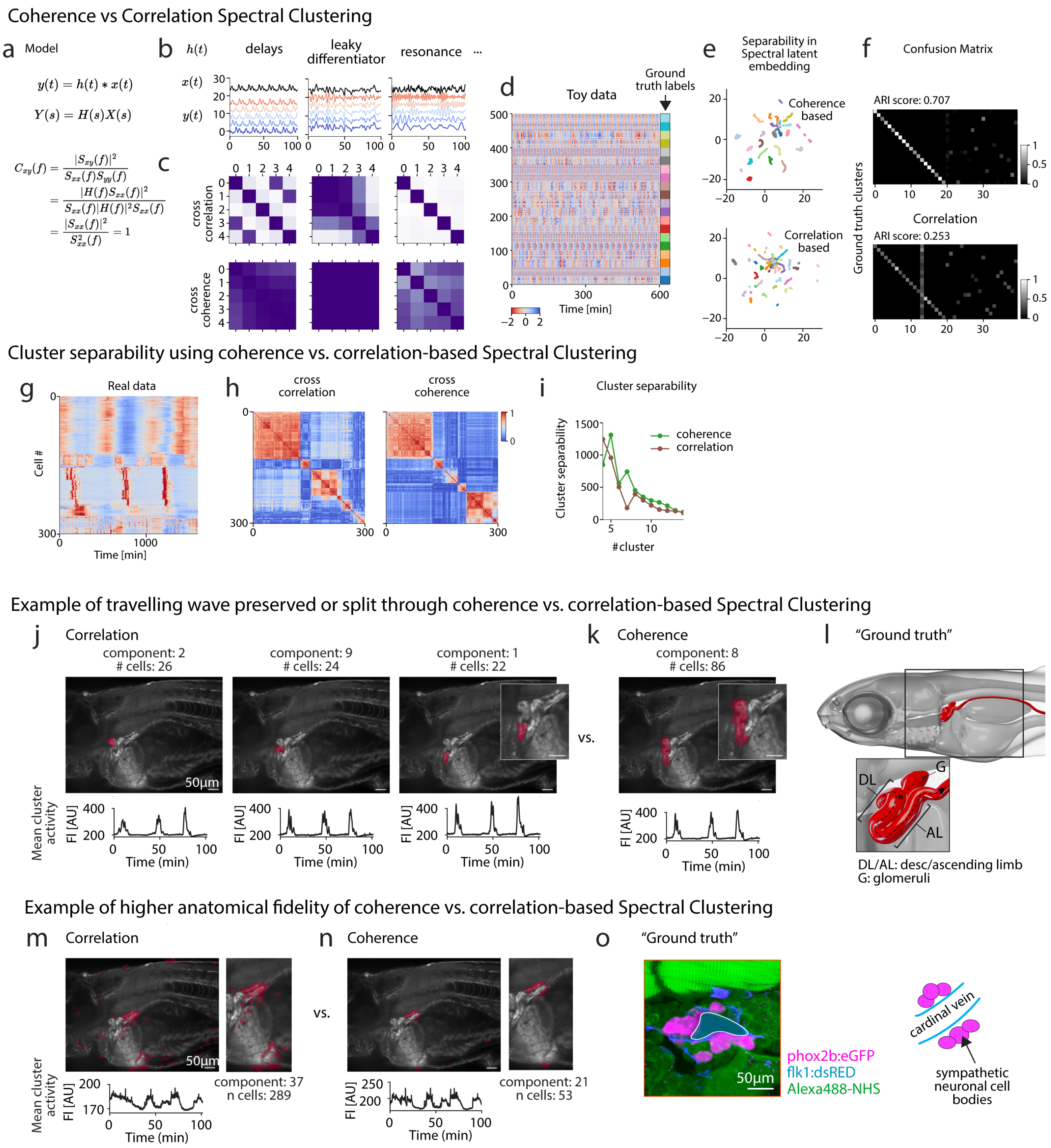

**Extended Data Fig. 4: Compactness of coherence vs. correlation-based spectral clustering.**

#### Extended Data Fig. 4: Compactness of coherence vs. correlation-based spectral clustering.

**a**, Any two signals $y(t)$, $x(t)$ which are linearly filtered version of each other will have a coherence of one. Definition of coherence.
**b**, Example of time-series which are coherent but not necessarily correlated. Unit $y(t)$s are filtered versions of a unit $y(t)$, illustration delay lines (left), leaky differentiators (center) and resonance (left).
**c**, Cross-coherence vs cross-correlation matrix of signals in b. Coherence better captures the relationship between these timeseries.
**d**, Toy data constructed as in b). The data consists of 20 clusters, which consist of noisy coherent timeseries. Left: ground truth labels of cluster identity.
**e**, Spectral latent embedding. Embedding of timeseries using coherence-based embedding clustering clustering is running using coherence-based clustering (top) or correlation-based clustering. **f**, Confusion matrix. Results of coherence vs correlation-based clustering on toy data fit with 40 clusters. Values correspond to fraction of cells that were correctly assigned. ARI (adjusted random score): 0.707 for coherence, and 0.253 for correlation.
**g**, Activity data matrix showing varying dynamics, sorted using RasterMap.
**h**, Cross-correlation and cross-coherence matrix sorted through hierarchical clustering.
**i**, Cluster separability is similar between coherence and correlation-based spectral clustering. **j**, Comparison of coherence vs. correlation-based spectral clustering. Using correlation as a metric, the descending limb of the kidney is clustered into three distinct clusters. Anatomical footprint of clusters (red) superimposed on anatomical reference (gray). Mean cluster activity trace (bottom).
**h**, Using coherence as a metric, the descending nephron is grouped into a single cluster. Anatomical footprint of clusters (red) superimposed on anatomical reference (gray). Mean cluster activity trace (bottom).
**i**, Model of zebrafish nephron based on WB-ExM-Histo data (same as in Fig. 2).
**j**, Anatomical fidelity of the clusters. The clusters using coherence tend to be more compact allowing for easier anatomical identification. Using correlation as a metric, a group of cells around the central vein showing bistable dynamics are grouped together; however, the cluster is contaminated by other cell types. Anatomical footprint of clusters (red) superimposed on anatomical reference (gray). Mean cluster activity trace (bottom).
**k**, Using coherence as a metric, a group of cells around the central vein are systematically pulled out. Anatomical footprint of clusters (red) superimposed on anatomical reference (gray). Mean cluster activity trace (bottom).
**l**, The functional tissue compartment identified in **d** and **e**, likely maps to sympathetic ganglion which is indeed compactly located around the central vein. Expansion microscopy of double transgenic fish labeling the autonomic nervous system (*Tg(phox2bb:eGFP)*, magenta), the vasculature (*Tg(flk1:dsRED)*, blue), with WB-ExM-Histo (Alexa488-NHS, green).

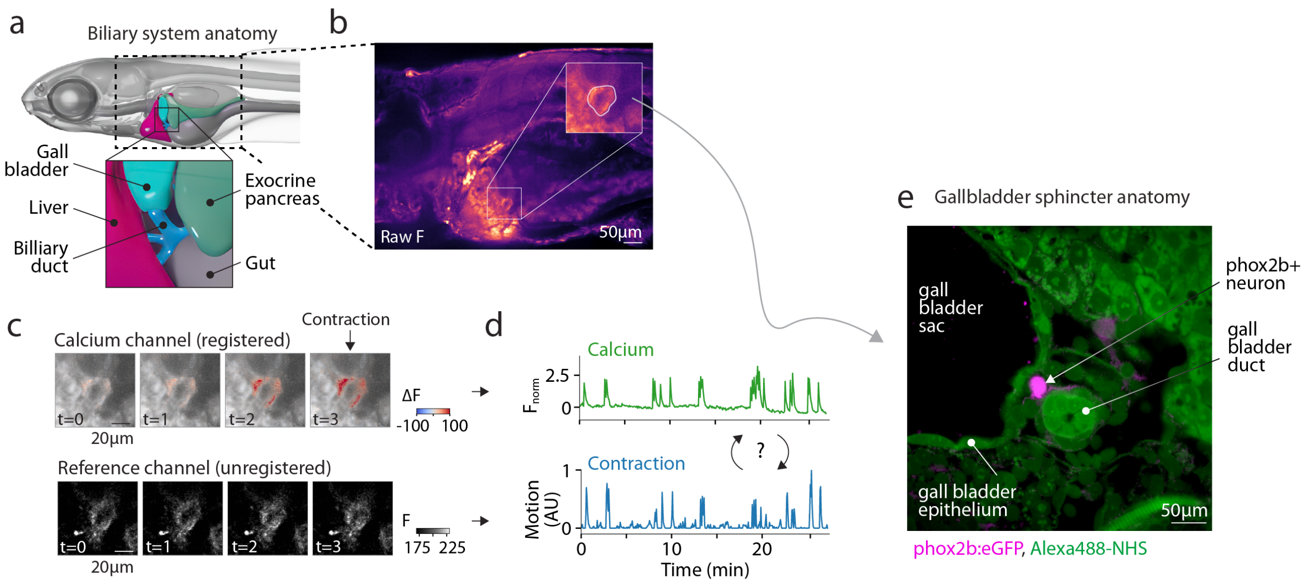

**Extended Data Fig. 5: Example of functional tissue compartments identified through WHOLISTIC coherence-based spectral clustering: biliary system.**
**a**, Contractile biliary sphincter with contraction-locked calcium activity bursts. ExM-based reconstruction of the hepato-biliary duct system.
**b**, Sagittal view showing a sphincter at the base of the gallbladder sac, with an enlarged view and outline of the associated functional tissue compartment.
**c**, Calcium activity concurrent with sphincter contraction. *Top*: Registered $\Delta F$ (red) on anatomical reference (gray). *Lower*: Unregistered reference showing sphincter constriction.
**d**, *Top*: average $Ca^{2+}$ activity trace from the sphincter functional tissue compartment. *Bottom*: average motion magnitude in the same region computed from the fitted motion field of the area.
**e**, Visualization of gallbladder sphincter using WB-ExM. View of biliary system with gallbladder sac, epithelia and duct visible, along with local post-ganglionic vagal neuron. WB-ExM of transgenic fish labeling the autonomic nervous system (*Tg(phox2bb:eGFP)*, magenta, total protein stain Alexa488-NHS, green).

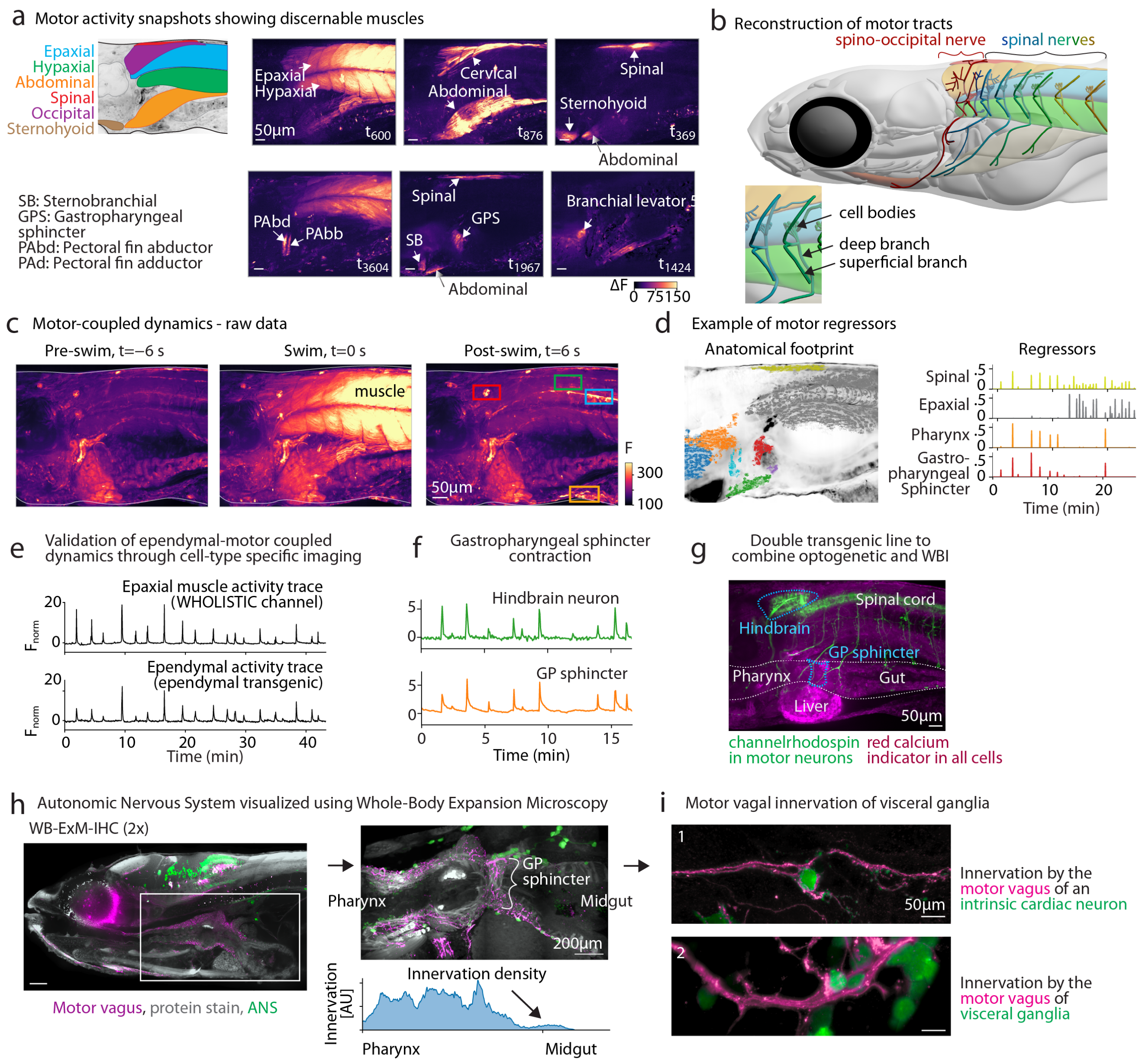

**Extended Data Fig. 6: Body-wide motor coupled dynamics.**

#### Extended Data Fig. 6: Body-wide motor coupled dynamics.

**a**, Distinguishable muscle groups. Example timepoints highlighting the contraction of various muscle groups. *Left to right*: 1) Outline of muscles, 2) epaxial and hypaxial, 3) cervical portion of the epaxial muscle and abdominal muscle (same as in Fig. 2), 4) spinal muscle (not yet reported in the literature) and sternohyoid muscle, 5) fin abductor and adductor muscles, 6) gastropharyngeal sphincter muscle, sternobranchial muscle, 7) branchial levator muscle.
**b**, Enlarged view of Fig. 2d showing reconstruction of spino-occipital nerves (red) innervating anterior portion of the epaxial muscle and spinal nerves innervating the remaining epaxial and hypaxial muscles. Each group of cell bodies project both deep and superficial axonal tracts.
**c**, Full field of view of data shown in Fig. 2h.
**d**, Spatial footprint and average time series of muscle regressors used.
**e**, Trace of average ependymal cell activity along the hindbrain and spinal cord extracted from *Tg(foxj1a:GCaMP7f)* channel and concurrent average epaxial activity extracted from *Tg(ubi:tTA;TRE:jRGECO1b)*.
**f**, Trace of gastropharyngeal mean sphincter activity used with the lag-regression model and example hindbrain neuron identified as correlated.
**g**, Double transgenic fish for optogenetic activation experiments. A transgenic line expressing channelrhodopsin in motor neurons is crossed to a red pancellular calcium indicator (*Tg(VAChTa:CoChR-eGFP)*; *Tg(ubi:tTA;TRE:jRGECO1b)*). Maximum projection of the green channel (green), overlaid on midline plane of red channel for anatomical reference (magenta).
**h**, *Left*: sagittal section of double transgenic animal labeling the motor vagus and autonomic nervous system (*Tg(isl1CREST-hsp70l:mRFP)* x *Tg(phox2bb:eGFP)*), stained against RFP (magenta) and eGFP (green) with total protein stain (Alexa488-NHS, gray) (WB-ExM-IF, expanded ∼2×). *Top right*: enlarged view of the anterior gastrointestinal tract. *Bottom right*: innervation density along the anterior-posterior axis showing dense innervation up to, but not beyond, the anterior gut.
**i**, WB-ExM-IF highlights innervation of visceral post-ganglionic neurons by motor vagus. *Top*: innervation of intrinsic cardiac neurons by motor vagus. *Bottom*: innervation of enteric neuron by motor vagus.

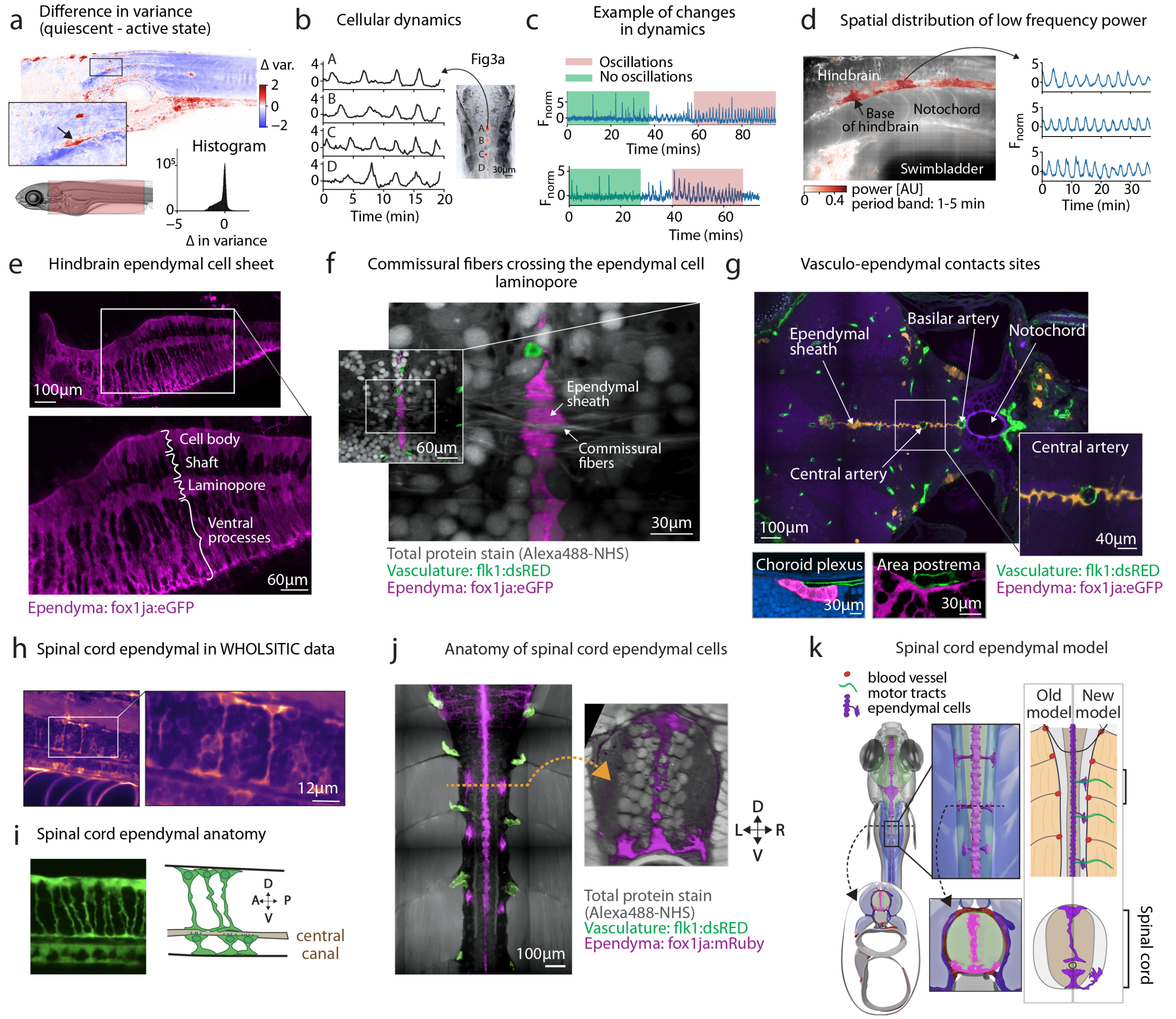

**Extended Data Fig. 7: Motor-quiescence coupled ultraslow oscillations and cell-type of origin.**

#### Extended Data Fig. 7: Motor-quiescence coupled ultraslow oscillations and cell-type of origin.

**a**, Ultra-slow oscillations during periods of prolonged quiescence. *Top*: maximum projection of difference in activity variance across the body between periods of quiescence and rest. Enlarged view of ventral hindbrain. *Bottom*: histogram of change in variance of cells.

**b**, Example of ultra-slow dynamics at different regions along the hindbrain and spinal cord. Regions show similar frequency, however, appear phase shifted. (Same animal as in Fig. 3a).

**c**, Example of changes in cellular dynamics observed with cells initially without oscillations (green phase) gradually displaying slow oscillations (magenta phase).

**d**, Spatial distribution of low frequency power during periods of quiescence mostly localizes to the base of the hindbrain and spinal cord.

**e**, *Top*: hindbrain ependymal cell midline sheet viewed sagittally. *Bottom*: enlarged view annotated with location of cell body layer, shaft, laminopore and ventral processes layer. Sagittal section of transgenic sample labeling the ventricular system (*Tg(foxj1a:eGFP), magenta*, labelled and expanded using WB-ExM-IF.

**f**, Visualization of commissural fibers crossing the midline at the level of the laminopore. Total protein stain (Alexa488-NHS, gray), vasculature (*Tg(flk1:dsRED-CAAX)*, green), ependyma (*Tg(foxj1a:eGFP)*, magenta).

**g**, Visualization of vasculo-ependymal cell contact sites with annotations. Same sample as in Fig. 3g. Total protein stain (Alexa488-NHS, dark blue), vasculature (*Tg(flk1:dsRED-CAAX)*, green), ependyma (*Tg(foxj1a:eGFP)*, gold top, magenta bottom).

**h**, Appearance of spinal cord ependymal cells in WHOLISTIC data. Sagittal view of the spinal cord of *Tg(ubi:tTA;TRE:GCaMP7f)* line.

**i**, Appearance of spinal cord ependymal cells in transgenic line labeling motile ciliated cells. *Left*: sagittal view of the spinal cord of *Tg(foxj1a:eGFP* line imaged using spinning disk confocal microscopy. *Right*: graphical representation of the dorsal and ventral ependymal cell population.

**j**, Visualization of spinal cord ependymal cells in WB-ExM-IF data. Same sample as **f**. *Left*: dorsal view. *Right*: coronal view highlighting bilateral projections in between somites, location of spinal nerve exit sites. Total protein stain (Alexa488-NHS, gray), vasculature (*Tg(flk1:dsRED-CAAX)*, green), ependyma (*Tg(foxj1a:eGFP)*, magenta).

**k**, Graphical model of spinal cord ependymal cells.

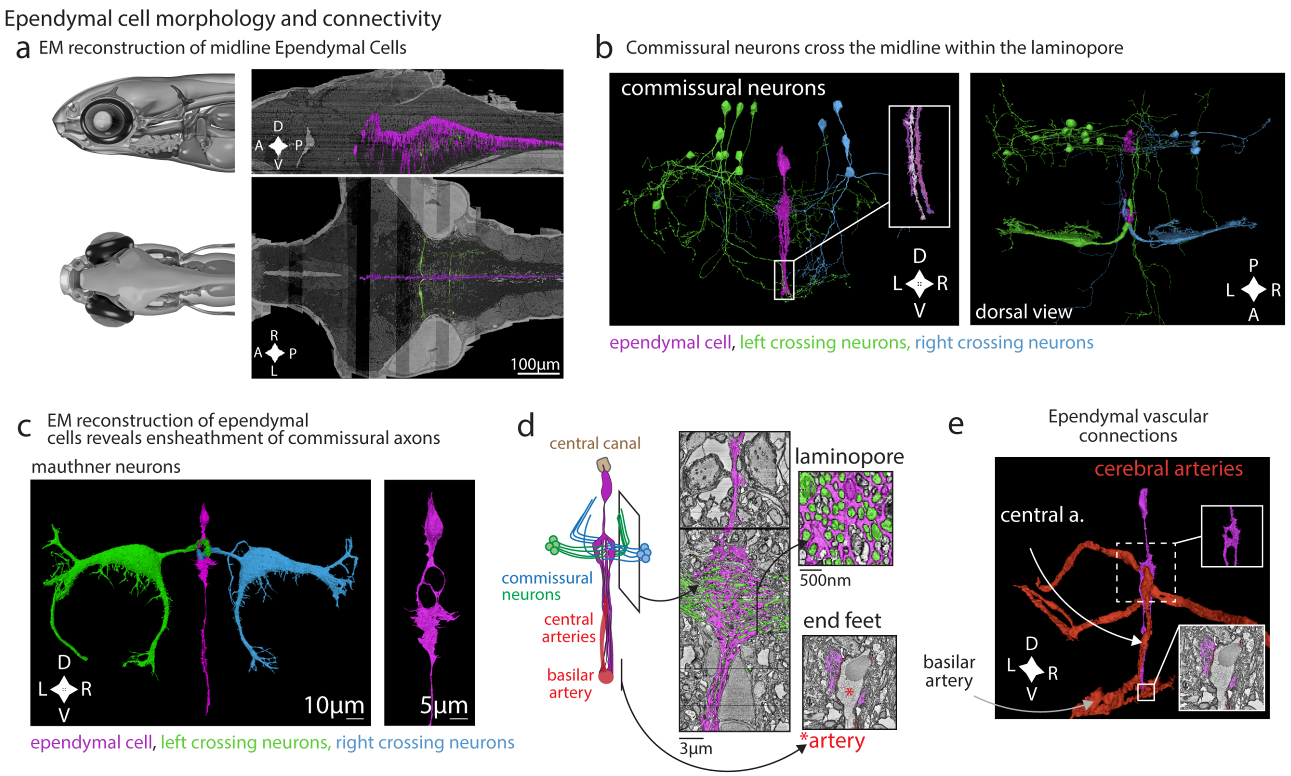

#### Extended Data Fig. 8: Ependymal cell morphology and connectivity reconstructed from EM data.

**a**, Midline ependymal cells within EM data. EM volume with subset of hindbrain ventral ependymal cells labeled (magenta). Ependymal cells form a thin dense sheath.

**b**, Reconstruction of example ependymal cells with commissural neurons crossing at the level of the laminopore in coronal view (left) and dorsal view (right). Midline ependymal cell (magenta), left commissural neurons (green), right commissural neurons (blue).

**c**, Reconstruction of example ependymal cells with Mauthner neurons crossing at the level of the laminopore. Coronal view of EM reconstruction. Midline ependymal cell (magenta), left Mauthner neuron (green), right Mauthner neuron (blue).

**d**, Left: schematic of midline ependymal cell anatomy. Right: EM data with ependymal cell (magenta) and ensheathed commissural axons (green) at the level of the laminopore.

**e**, Reconstruction of example ependymal cells along with cerebral vasculature. The basilar artery, located at the base of the brain, branches into central arteries that course up the midline, some of which are ensheathed by ependymal cell laminopores. Inset: EM data with ependymal cell (magenta) appositioning against the basilar artery.

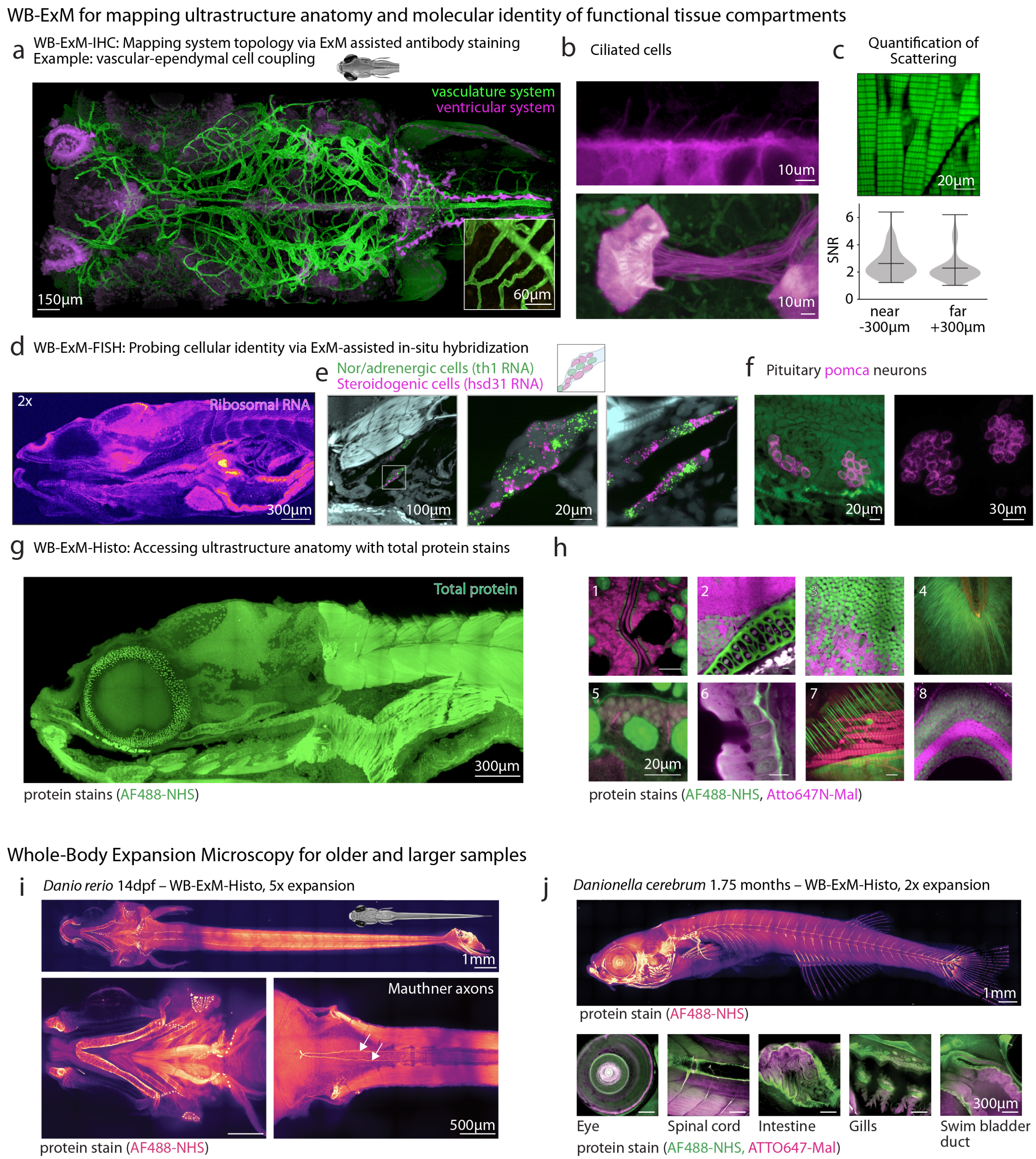

**Extended Data Fig. 8: Whole-Body Expansion Microscopy (WB-ExM) to map ultrastructure, RNA and protein across the body.**

#### Extended Data Fig. 8: Whole-Body Expansion Microscopy (WB-ExM) to map ultrastructure, RNA and protein across the body.

**a**, Enlarged view of Fig. 3g. Dorsal section of double transgenic sample labeling the ventricular and vascular systems (*Tg(foxj1a:eGFP)* x *Tg(flk1:dsRED-CAAX)*), stained against eGFP (magenta) and dsRED (green) (10 days post-fertilization, expanded ∼2×).

**b**, Enlarged view of a cell from sample **a** in the nephron epithelium projecting cilia into the nephric lumen.

**c**, Quantification of clearing quality of WB-ExM sample. *Top*: Image of muscle sarcomeres in WB-ExM sample stained with total protein stain (Alexa488-NHS). *Bottom*: Distribution of signal to noise ratio (peak to trough) in sarcomeres at far and near side of the sample.

**d**, Whole-Body Expansion Microscopy combined with fluorescent in situ hybridization (WB-ExM-FISH) enables RNA-based cell identification across tissue. Sagittal section of an expanded sample stained against ribosomal RNA, demonstrating full probe access to all tissues (8 days post-fertilization, ∼2× expanded).

**e**, Localization of nor/adrenergic cells and steroidogenic cells using WB-ExM-FISH. *Left*: sagittal section of an expanded sample (2×) stained against *th* (green) and *hsd31* (magenta) RNA and with total protein stain (Alexa488-NHS, cyan). *Right*: enlarged view showing intermingled cellular populations.

**f**, Localization of pituitary pomc neurons using WB-ExM-FISH, cells situated at the base of the brain that are typically hard to optically access. *Left*: coronal view of expanded sample (2×) stained against *pomca* (magenta) RNA and with total protein stain (Alexa488-NHS, green), showing two distinct populations. *Right*: rendering of volume.

**g**, Whole-Body Expansion Microscopy combined with total protein stains (WB-ExM-Histo) to provide contextual information.

**h**, Higher magnification views of expanded sample: 1) enteric goblet cell, 2) osteochondral tissue at the base of the brain, 3) midbrain neurons, 4) tail fin, 5) epithelial cell of nephron with primary cilia visible, 6) intestinal epithelium with basement membrane, 7) collagen fibers, 8) multilayered retina. Total protein stains: Alexa488-NHS (green) and Atto647N-Maleimide (magenta).

**i**, Whole-Body ExM for older samples. WB-ExM-Histo of 14 days post fertilization zebrafish after 4 gelation rounds leading to ∼5× expansion, stained with total protein stain Alexa488-NHS.

**j**, Whole-Body ExM for *Danionella cerebrum*. WB-ExM-Histo of 1.75 months *Danionella cerebrum* stained with total protein stain Alexa488-NHS (green) and Atto647N-Maleimide (magenta). *Top*: maximum intensity projection of entire sample. *Bottom, left to right*: enlarged views of same sample showing 1) eye, 2) spinal cord, 3) intestine, 4) gills, 5) swim bladder duct.

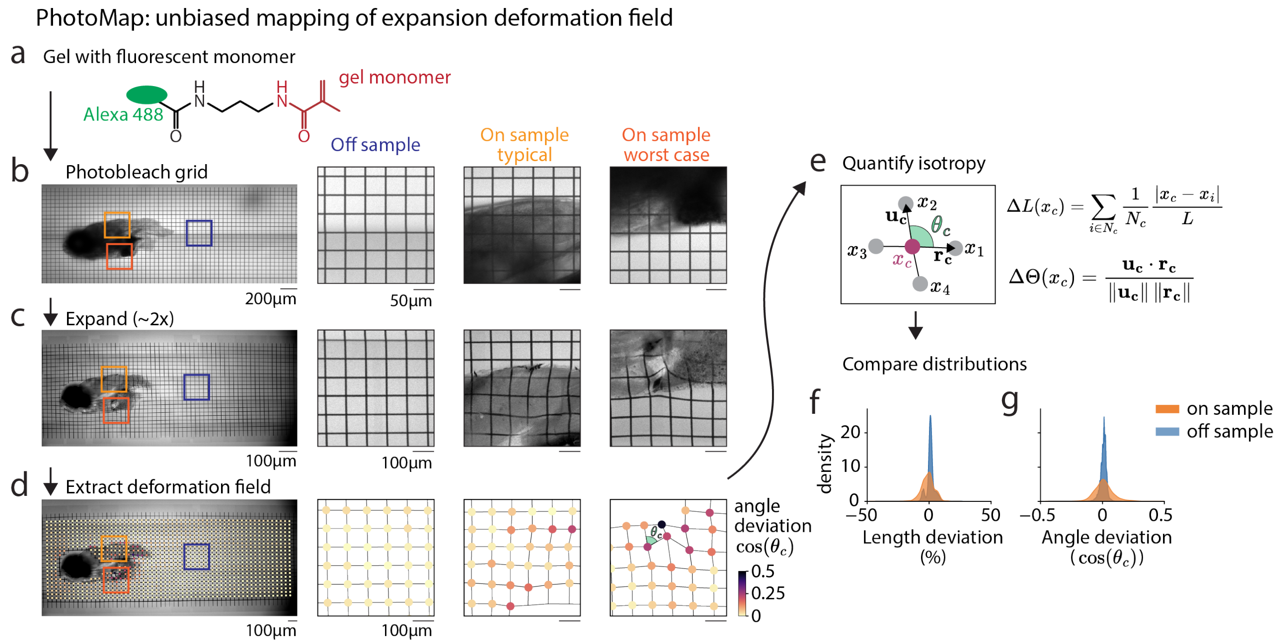

$$\Delta L(x_c) = \sum_{i \in N_c} \frac{1}{N_c} \frac{|x_c - x_i|}{L}$$

$$\Delta\Theta(x_c) = \frac{\mathbf{u_c} \cdot \mathbf{r_c}}{\|\mathbf{u_c}\| \, \|\mathbf{r_c}\|}$$

**Extended Data Fig. 10: PhotoMap: Unbiased mapping of expansion deformation field.**

**a**, Schematic of fluorescent monomer. A fluorescent dye is covalently conjugated to the gel monomer, rendering the gel intrinsically fluorescent.

**b**, Grid photobleaching. Using a two-photon microscope, a three-dimensional grid ($50 \times 50 \times 50$ $\mu$m) was photobleached into both the sample and the surrounding gel. *Left*: whole-gel image with the grid visible. *Right*: example crops showing regions containing no sample, brain tissue, and the cleithrum, the earliest mineralizing bone in zebrafish[?] .

**c**, Visualization of deformation after expansion. Same views as in **b** imaged after tissue disruption and gel expansion.

**d**, Extraction of the deformation field. Grid intersections were detected throughout the sample. Same views as in **b**, with intersection points color-coded by their deviation from the ideal 90° intersection angle.

**e**, Quantification of isotropy. For each grid intersection, the mean deviation from the target length $L$ to its four nearest neighbors was measured, along with the deviation from the ideal 90° intersection angle.

**f**, Deformation field comparison. Distributions of length deviation (*left*) and angle deviation (*right*) measured in off-sample (blue) and on-sample (orange) regions. On-sample distributions are broader but remain within a relatively narrow range.

**a**, Schematic of fluorescent monomer. A fluorescent dye is conjugated to the gel monomer to make the gel intrinsically fluorescent.

**b**, Grid photobleaching. Using a 2 photo microscope a 3D grid is photobleached into the sample (50 x 50 x 50 µm) both outside and on the sample. Left: image of entire gel with grid visible. Right: three example crops showing area containing no sample, area containging brain, area containing the cleithrum bone, the earliest bone to mineralize in zebrafish[?] .

**c**, Expansion of grid allows for easy visualization of deformation. Same views as in b) imaged after tissue disruption and expansion.

**d**, Extraction of deformation field. Grid intersections are located throughout the sample. Same views as in b) with identified intersection points are color coded according to the angle deviation from the ideal 90° angle.

**e**, Graphic illustration quantification of isotropy. For each intersection point, the average deviation from the ideal length $L$ from its surrounding neighbors is measured, as well as deviation from 90° angle.

**f**, Deformation field comparison. Distribution of length deviation (left) and angle deviation (right) off sample (blue) and on sample (orange). The on sample distribution is broader but remains in a relatively constrained range.

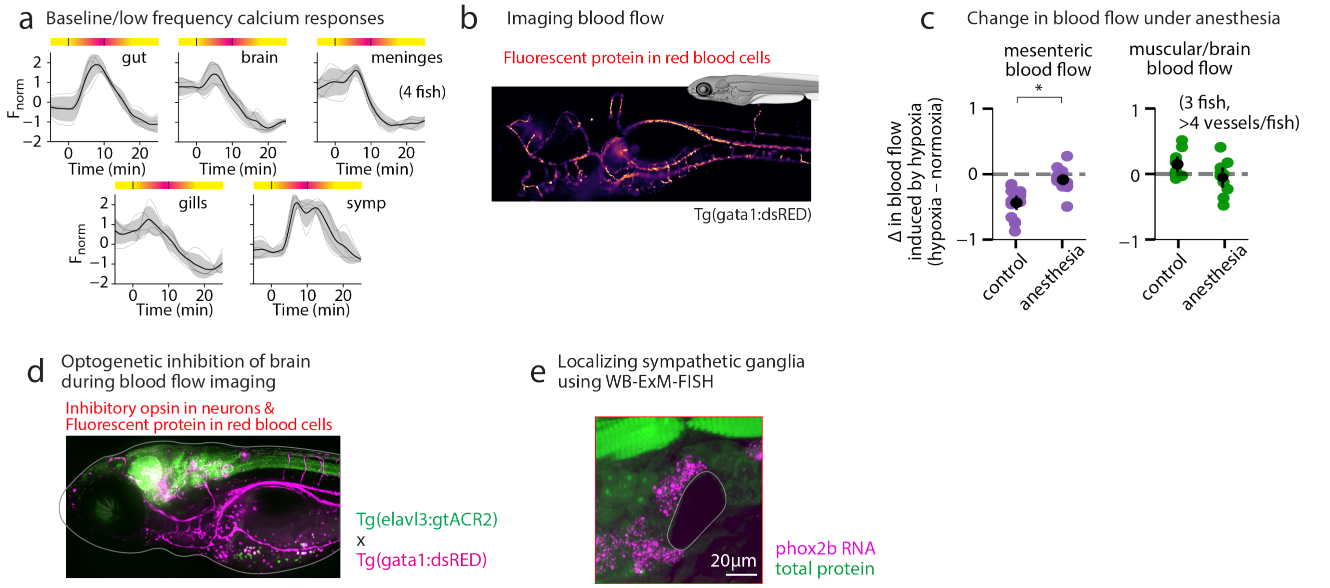

**Extended Data Fig. 9: Body-wide circuit engaged in response to hypoxic stress.** **a**, Hypoxia induces baseline changes in calcium levels in multiple organs. Average organ baseline calcium levels (dark: mean response, light: individual animals, shaded gray: standard deviation, N=4). **b**, Hypoxia induces reduction of blood flow to the gut. Genetic strategy for monitoring blood flow during normoxia and hypoxia using a transgenic line labeling red blood cells (*Tg(gata1:dsRED)*).
**c**, Quantitative comparison of blood flow changes in brain/muscle (green) and visceral organs (purple) during hypoxia, with and without anesthesia.
**d**, Double transgenic fish for optogenetic inhibition experiments. A transgenic line expressing gtACR2 (*Tg(elavl3:gtACR2)*), an inhibitory opsin, in all neurons is crossed a transgenic line labeling red blood cells (*Tg(gata1:dsRED)*). Maximum projection of the green channel (green), overlaid on midline plane of red channel (magenta).
**e**, Localization of the sympathetic ganglia using WB-ExM-FISH. Sagittal section of animal probed for phox2b mRNA (magenta), stained with total protein stain, Alexa488-NHS (green). A cluster of phox2b+ cells localizes below the spinal cord, surrounding the cardinal vein. (WB-ExM-FISH, 7 days post-fertilization, expanded ∼2×)

### Supplementary movies

##### Supplementary Video 1: WHOLISTIC imaging - sagittal view.

Sagittal view of *Tg(ubi:tTA; TRE:GCaMP7f)* line with major organs indicated, imaged with 20× 0.75 NA air objective (sped up ∼50×, raw data).

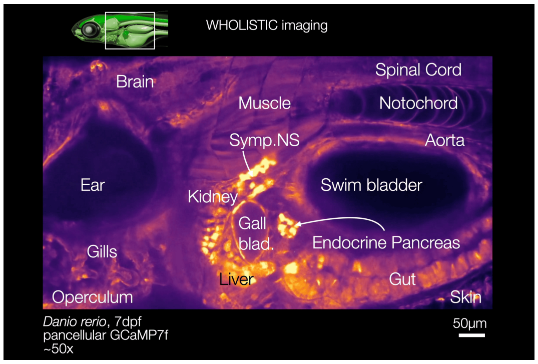

**Supplementary Video 2: WHOLISTIC imaging - dorsal view.**

Dorsal view of *Tg(ubi:tTA; TRE:GCaMP7f)* line with major organs indicated, imaged with 10× 0.45 NA air objective (sped up ∼50×, raw data).

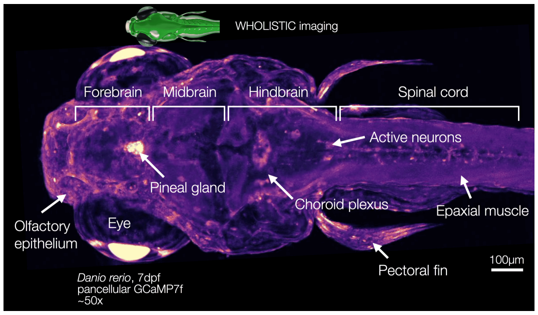

**Supplementary Video 3: WHOLISTIC imaging of enteric activity highlighting visceral motion.**

Sagittal view of *Tg(ubi:tTA; TRE:GCaMP7f)* line with major organs indicated, imaged with 40× 1.15 NA water objective (sped up ∼50×, raw data).

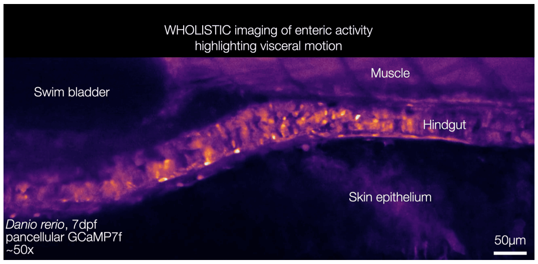

**Supplementary Video 4: Registration results.**

Sagittal view of registration channel derived from *Tg(β-actin:mCherry-CAAX)* line, imaged with 20× 0.75 NA air objective. *Left*: data pre-registration. *Right*: data post-registration.

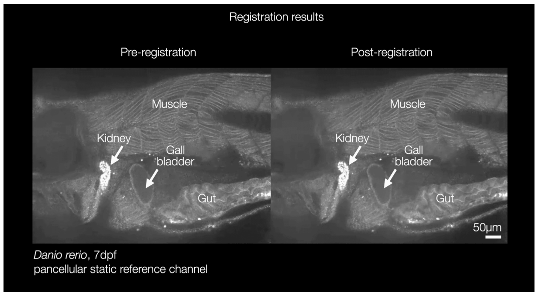

**Supplementary Video 5: WHOLISTIC imaging of cold response.**

Dorsal view of *Tg(ubi:tTA; TRE:GCaMP7f)* line, imaged with 40× 1.15 NA WI objective, in response to the delivery of cold water (sped up ∼50×, raw data). Onset of cold water indicated on the top right.

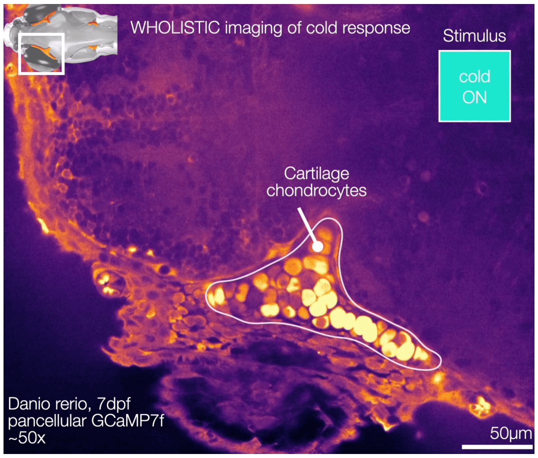

**Supplementary Video 6: Whole-Body Expansion Microscopy.**

Visualization of transgenic animal labeling the autonomic nervous system (*Tg(phox2bb:eGFP)*), stained against eGFP (magenta) with total protein stain (Alexa488-NHS, green) (WB-ExM-IF, expanded ∼2×).

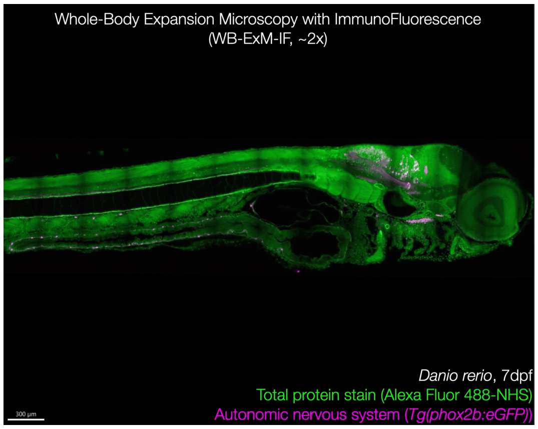

**Supplementary Video 7: 3D geometric model of young zebrafish.**

3D model derived from data generated by Whole-Body Expansion Microscopy data containing the major organs of the young zebrafish. Model build and rendered using the software Blender.

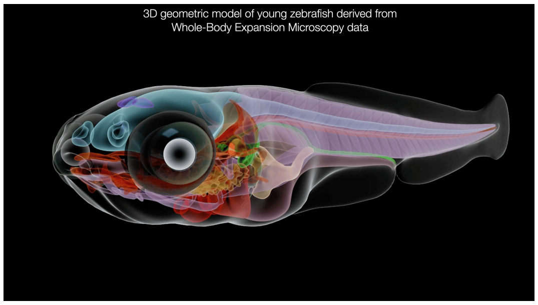

**Supplementary Video 8: Visualization of hindbrain ependymal sheet using Whole-Body Expansion Microscopy with immunofluorescence.**

Visualization of ventral ependymal sheath using WB-ExM-IF. Sagittal view of transgenic sample labeling the ventricular system (*Tg(foxj1a:eGFP)*), stained against eGFP (magenta) (WB-ExM-IF, expanded ∼2×).

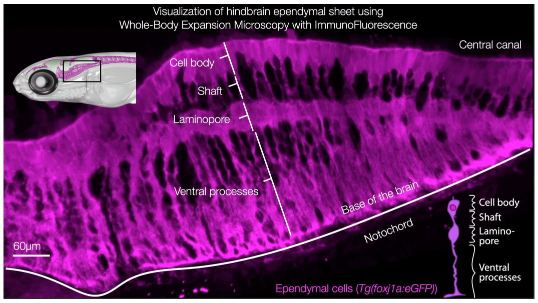

**Supplementary Video 9: Hypoxia induced blood flow redirection.**

Blood flow imaged during normoxia (*left)* and hypoxia (*right*). Sagittal section of a transgenic animal with fluorescently labeled red blood cells (*Tg(gata1:dsRED)*), imaged with 20× 0.75 NA air objective (sped up ∼1.5×, raw data), showing a reduction in mesenteric blood flow upon exposure to hypoxia. This reduction is readily evident in the vascular supply to the endocrine pancreas (indicated in box).

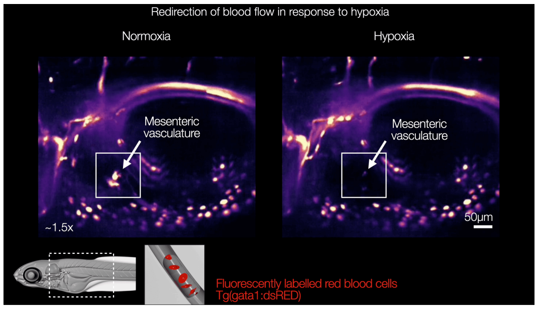

### Supplementary notes

#### Notes on imaging and tissue identification

We explored alternative imaging approaches, including the imaging of larger areas and perspectives from different angles, such as dorsal, ventral, and head-on views (Extended Data Fig. 1a,c,d). The identification of all areas was achieved by comparing them with established anatomical references, including histological and electron microscopy databases[2,3,4,5,6]. A minority, approximately 10% of other regions exhibit characteristic textures but remain more difficult to classify (Extended Data Fig. 1a). The ventral view capturing the gills exemplifies a region where it remains challenging to identify tissue boundaries and determine the identity of individual cells. We note that the eye, ear, and swim bladder introduce scattering, resulting in partial occlusion of the underlying structures, including the forebrain, thalamus, and part of the nephric duct.

#### Notes on coherence-based spectral clustering

Spectral clustering is a well-established method for clustering data[7,8]. The method requires one to define a similarity function between data points, which is often set to the Euclidean distance. However, such a metric is not necessarily a meaningful distance to use between two time series. Correlation would be a better distance measure, but fails to encode that time-lagged activity traces should ideally be considered as close, which results in the breaking up of such groups of traces into multiple clusters. Therefore, we aim to define a distance based on coherence (which is small when coherence is high and vice versa) that can be utilized in spectral clustering.

Background. In spectral clustering, one first defines an adjacency graph. A common choice of similarity function is the Gaussian similarity function defined between $x_i$ and $x_j$ as:

$$\mathcal{W}_{i,j} = \exp\left(-\frac{|x_i - x_j|_2^2}{\tau}\right) \tag{18}$$

where $\tau$ defines the range of distances of interest, or width of the neighborhood.

One then builds the normalized graph Laplacian of the adjacency matrix:

$$L = D - W \tag{19}$$

$$L_{\text{norm}} = I - D^{-1}W \tag{20}$$

where $D$ is a diagonal matrix and $d_{i,i} = \sum_j W_{i,j}$. The lowest N eigenvectors of the matrix are then computed and used as the encoding basis set, and finally the k-means algorithm is run and outputs N clusters.

For intuition, we can interpret $L$ as being a transition function of the consensus dynamics of the system. Assuming rows of $W$ already sum to 1 for simplification:

$$\frac{dx}{dt} = -Lx \tag{21}$$

$$\frac{dx}{dt} = -(I - W)x = -x + \sum_j W_{i,j}x \tag{22}$$

where the fixed point is:

$$x = \sum_j W_{i,j}x \tag{23}$$

which is the weighted average of $x$ (also showing that it has to be stable). The eigenvalues thus tell us how quickly consensus, or information, will be distributed. It can be shown that the graph has as many 0 eigenvalues as disconnected components; indeed, consensus would never occur in disconnected graph. The smaller the eigenvalues the slower the diffusion within that mode, and the similarity of the weights of the eigenvectors indicate which entries are similar. Indeed, for any vector $f$:

$$f'Lf = f'Df - f'Wf \tag{24}$$
$$= \sum_{i=1}^{n} d_i f_i^2 - \sum_{i,j=1}^{n} f_i f_j w_{ij} \tag{25}$$
$$= \frac{1}{2}\left(\sum_{i=1}^{n} d_i f_i^2 - 2\sum_{i,j=1}^{n} f_i f_j w_{ij} + \sum_{j=1}^{n} d_j f_j^2\right) \tag{26}$$
$$= \frac{1}{2}\sum_{i,j=1}^{n} w_{ij}(f_i - f_j)^2. \tag{27}$$

which is minimized if for large similarity $w_{ij}$, $f_i$ and $f_j$ are close.

Derivation. We first can show that if one wanted to use correlation as similarity metric, we can define a distance which depends on correlation (small when correlation is high and vice-versa) and which can be rewritten as the Euclidean distance between vectors. This then ensures that we can use this defined distance in a Gaussian kernel and that it remains positive semi-definite.

The correlation between vectors $\boldsymbol{x} \in R^N$ and $\boldsymbol{y} \in R^N$ being defined as:

$$\mathcal{R}(\boldsymbol{x}, \boldsymbol{y}) = \frac{\sum_i x_i \cdot y_i}{\sqrt{\sum_i x_i^2}\sqrt{\sum_i y_i^2}} \tag{28}$$

We assume already pre-normalized vector:

$$x_i = \frac{x_i}{\sqrt{\sum_i x_i^2}} \tag{29}$$

Then:

$$|\boldsymbol{x} - \boldsymbol{y}|_2^2 = 2\sum_i (1 - x_i \cdot y_i) \tag{30}$$
$$= 2(1 - \sum_i (x_i \cdot y_i)) \tag{31}$$
$$= 2(1 - \mathcal{R}(\boldsymbol{x}, \boldsymbol{y})) \tag{32}$$

Thus, the Euclidean distance between normalized vector defines a distance based on correlation.

In the same spirit, one can define a distance based on coherence and show that it can be written as a Euclidean distance.

By definition, coherence is defined as:

$$\mathcal{C}(\boldsymbol{x}, \boldsymbol{y})(\omega) = \frac{|\sum_k \hat{x}_k(\omega)\overline{\hat{y}_k(\omega)}|^2}{\left(\sum_k \hat{x}_k(\omega)\overline{\hat{x}_k(\omega)}\right)\left(\sum_k \hat{y}_k(\omega)\overline{\hat{y}_k(\omega)}\right)} \tag{33}$$

We assume an already normalized spectrum to not carry the normalization term:

$$\sum_k \hat{x}_k(\omega)\overline{\hat{x}_k(\omega)} = 1 \tag{34}$$

Coherence can then be re-written as:

$$\begin{aligned}
\mathcal{C}(\boldsymbol{x}, \boldsymbol{y})(\omega) &= \left(\sum_k \hat{x}_k(\omega)\overline{\hat{y}_k(\omega)}\right)\left(\sum_l \hat{y}_l(\omega)\overline{\hat{x}_l(\omega)}\right) && (35)\\
&= \left(\sum_k \sum_l \hat{x}_k(\omega)\overline{\hat{y}_k(\omega)}\hat{y}_l(\omega)\overline{\hat{x}_l(\omega)}\right) && (36)\\
&= \operatorname{vec}\left(\hat{X}(\omega)\overline{\hat{X}(\omega)}^\top\right)^\top \operatorname{vec}\left(\hat{Y}(\omega)\overline{\hat{Y}(\omega)}^\top\right) && (37)
\end{aligned}$$

where $\hat{X}(\omega) \in R^k$ is a vector of all of the $\hat{x}_k(\omega)$. Thus, we define $Z(\omega) = \operatorname{vec}\left(\hat{X}(\omega)\overline{\hat{X}(\omega)}^\top\right)$ and $W(\omega) = \operatorname{vec}\left(\hat{Y}(\omega)\overline{\hat{Y}(\omega)}^\top\right)$

Thereby, we can define a distance based on coherence which can be written as a Euclidean distance between two vectors, thus guaranteeing that one can use this distance and have a Gram matrix that remains psd.

$$\begin{aligned}
|Z(\omega) - W(\omega)|_2^2 &= \sum_k \sum_l (\hat{x}_k(\omega)\overline{\hat{x}_l(\omega)})^2 + \sum_k \sum_l (\hat{y}_k(\omega)\overline{\hat{y}_l(\omega)})^2 - 2\mathcal{C}(\boldsymbol{x}, \boldsymbol{y}) && (38)\\
&= 2(1 - \mathcal{C}(\boldsymbol{x}, \boldsymbol{y})) && (39)
\end{aligned}$$

Normalization. Typically coherence is averaged over frequencies:

$$\mathcal{C}(\boldsymbol{x}, \boldsymbol{y}) = \sum_{\omega} \mathcal{C}(\boldsymbol{x}, \boldsymbol{y})(\omega) \tag{40}$$

$$= \sum_{\omega} \frac{|S_{x,y}(\omega)|^2}{S_{xx}(\omega) S_{y,y}(\omega)} \tag{41}$$

This definition weights each $\mathcal{C}(\omega)$ at each frequency irrespective of the power in that frequency band, this can be detrimental as bands of low power can be dominated by noise. To avoid this, one can normalize by the total power in all frequency bands:

$$\mathcal{C}(\boldsymbol{x}, \boldsymbol{y}) = \frac{\sum_{\omega} |S_{x,y}(\omega)|^2}{\sum_{\omega} S_{xx}(\omega) S_{y,y}(\omega)} \tag{42}$$

which remains bounded between 0 and 1. In other words, one can normalize the entire vector $\hat{X} \in R^{K\Omega}$:

$$\hat{x}_k(\omega) \to \frac{\hat{x}_k(\omega)}{\sqrt{\sum_{\omega} \sum_k \hat{x}_k(\omega) \overline{\hat{x}_k(\omega)}}} \tag{43}$$

Comparing the use of correlation vs. coherence as metric in Spectral Clustering. As a motivating example as to the use of coherence vs. correlation as a metric, we run the Spectral Clustering algorithm with each metric on a single plane of WHOLISTIC imaging data keeping all the other parameters constant ($\tau$, the length-scale and $N$, the number of eigenmodes/clusters). We find that coherence-based spectral clustering groups together more compactly tissues: for instance, the proximal part of the nephric tube is grouped into one cluster despite their being significant time delay between dorsal and ventral portions as well as them exhibiting slightly differing waveforms (Extended Data Fig. 4b). In contrast, correlation-based spectral clustering systematically separates them at the cost of finding other clusters in the data (Extended Data Fig. 4a). This, in part, contributed to other clusters being more mixed (Extended Data Fig. 4d-f). We note that these result depend on the number of clusters asked, and when larger numbers of clusters are searched for, both methods break up the nephric tube into disjoint sets, as indeed cells within a segment of kidney are more coherent within than outside of that segment. In passing, we note that the histogram of coherence coefficients is less uniformly distributed than those of correlation which can guide the choice of $\tau$. Nonetheless both have a local peak at around 0.7 justifying comparing both methods with the same value of $\tau$ which we set to 0.3.

#### Notes on functional tissue compartments and screening for cellular responses

We further illustrate some functional tissue compartments identified through the WHOLISTIC workflow and how the motion flow field can be used itself to identify contractile tissue, using the biliary system as an example. While the majority of cells within the epithelium lining the gallbladder sac exhibited activity that appeared largely uncoordinated, a distinct ring-shaped area at the base of the gallbladder sac demonstrated coordinated calcium bursts (Extended Data Fig. 5a-c). By extracting the equivalent region in the original unregistered imaging data, it was found that these calcium bursts occur in synchrony with the constriction of the ring, suggesting the presence of a sphincter (Extended Data Fig. 5e). Subsequent localization of this region within Whole-Body Expansion Microscopy data revealed that this area aligns with the tripartite ductal system connecting the gallbladder, liver, and pancreas to the gastrointestinal tract. This finding posits that the identified contractile sphincter is homologous to the sphincter of Oddi[9], which has not been described in fish; in mammals, it regulates hepato-biliary and pancreatic flow into the gastrointestinal tract. Collectively, these findings underscore the capability of the WHOLISTIC workflow to identify small, localized functional units, such as the biliary sphincter, and to correlate their calcium dynamics with the underlying tissue contraction.

WHOLISTIC identifies new cellular response properties to stimuli. To illustrate the integration of WHOLISTIC and WB-ExM as a screening method for uncovering novel cellular properties of molecularly defined cell types, we conducted a WHOLISTIC screen to identify cells that are responsive to thermal changes, specifically a reduction in temperature (Extended Data Fig. 3). Young zebrafish were subjected to cool water (10°C) while calcium dynamics were monitored throughout the organism (see Methods). A specific cellular population exhibited robust, high-amplitude responses (Extended Data Fig. 3a-b). We matched the cells in WHOLISTIC data to the corresponding region in WB-ExM volumes through their characteristic morphology and anatomical distribution (Extended Data Fig. 3c, *left*), which showed that these cells localize to osteochondral tissue (Extended Data Fig. 3c, *center*)[10]. Finally, using WB-ExM-FISH, we established that these cells express col2a1, a marker for chondrocytes (Extended Data Fig. 3c, *right*)[11], establishing that chondrocytes are thermally responsive, displaying a rapid cold-triggered rise in intracellular calcium, a seconds-timescale responsiveness that has not been previously reported. Chondrocytes are known to maintain the extracellular matrix via calcium-dependent intracellular pathways[12,13] and express a diverse array of membrane channels that transduce external stimuli into intracellular signals[14], providing a possible mechanism for thermal responsiveness. For completeness, using WB-ExM data, we reconstructed the full skull of young zebrafish, which is the most challenging tissue in the specimen to expand (Extended Data Fig. 3d, see Methods). This example illustrates how combining WHOLISTIC and WB-ExM allows for the integration of functional properties, molecular cellular identity, and macroscopic anatomy, facilitating the discovery of previously unknown cellular properties.

WHOLISTIC identifies new cellular response properties to drugs. As a demonstration of using WHOLISTIC for drug screening, we performed a WHOLISTIC screen for cells that respond to ketamine, a dissociative anesthetic (Extended Data Fig. 4). Quantifying the strongest responding cells by ranking $F_{norm}$ increases, we identified a functional compartment that aligns with the outer perimeter of the brain, likely corresponding to the brain meninges (Extended Data Fig. 4a), which is responsive to ketamine (Extended Data Fig. 4b-c). While the effects of ketamine on neurons and glia are documented[15], the role of the meninges is less studied; meninges have not yet been functionally imaged in any species in vivo[16] and remain largely uncharacterized in zebrafish. With no existing selective driver line[17], they are difficult to segment based on anatomy due to the thinness of the tissue. This highlights how WHOLISTIC activity analysis is able to delineate functional compartments that are otherwise challenging to label – i.e., identify, within the many cells in the body, small groups of cells with specific dynamics that would otherwise remain undetected.

#### Notes on Whole-Body Expansion Microscopy

Expansion Microscopy (ExM) is a form of super-resolution microscopy. Unlike standard super-resolution methods, which either focus on altering the hardware or software, ExM focuses on altering the object to be imaged itself, making it physically larger[18], resulting in both high quality clearing and better effective resolution.

Gel engineering. We found, as previously reported[19], that having a harsh digestion with proteinase K (proK) could allow for more homogeneous expansion; however, expansion quality and antibody signal retention was variable between specimens, limiting the reliability and utility of the method. As a result, we turned to high temperature, high detergent disruption, in which heat is used to denature proteins, as reported in[20]. We reasoned that such hydrolysis-based disruption would result in chemical cleavage of proteins that would be less subject to crowding or to differences in local tissue environment as compared to enzymatic digestion. This approach indeed improved the robustness of digestion; however, as we began working with older samples, propagating cracks re-appeared. Over the course of embryological development, tissue heterogeneity throughout the animals dramatically increases, with for instance the rapid development of cartilage beginning at 6 days post fertilization, which progressively expands and ossifies. After trying a variety of gel recipes to address this, we found that an intermediate monomer concentration gel recipe was least prone to crack formation, and that by re-embedding immediately after disruption, without reducing salt concentration or expanding more than $\sim 1.5\times$, strongly reduced subsequent crack formation and that any cracks that did form (typically in cartilage) did not propagate into neighboring tissues (Extended Data Fig. 8i). An intermediate monomer concentration may produce a better match to tissue elasticity, while the modest first expansion enables more uniform access to the next round of polymers, thereby making the polymer density more homogeneous throughout irrespective of variation in tissue structure. This allows the expansion of zebrafish up to 14 dpf as well as mature *Danionella* (Extended Data Fig. 8i-j).

Immunofluorescence optimization. We found that strongly permeabilizing the fish was critical to avoid the skin impeding antibody access to the tissue, as well as increasing antibody (AB) concentration, incubation time, and temperature in the case of high-abundance targets. Increasing antibody concentration comes at the risk of higher background signal; however, a titration curve of antibody concentration vs. signal showed that the gain in signal far outweighs small increases in background and that at 1:100 we are still not reaching epitope saturation. With long incubations in rich serum, bacterial growth is more likely to occur leading to nonspecific background staining; this can be resolved by processing the samples with minimal delay and adding sodium azide from the point of fixation onward.

Sample mounting and overcoming edge effects. Since the head of the sample is generally wider than its body, placing samples on their sides often results in slanted samples. Despite this not being a problem for local measurements, this can result in having to acquire a much larger field of view and render sample alignment to a reference highly laborious, requiring hours of manual work. We found that fixed samples can be embedded in low melting temperature agarose (0.7%) where it is easy to optimally orient the sample, stabilizing it in place. Following high-temperature denaturation, agarose pre-embedding did not appear to hinder the ExM expansion process. This process results in more standardized samples. Another related issue was the changes in refractive index that can occur at the edges of the gel as a result of boundary conditions in the gelation process. We resolved this by increasing the agarose gel thickness, ensuring that the sample was embedded away from either surface of the gel preventing edge effects.

Detailed Whole-Body Expansion Microscopy (WB-ExM) protocols. WB-ExM protocols were formatted using the protocols.io web platform. After publication, the protocols will be made accessible at protocols.io.

dr8d59s6

### Whole-Body Expansion Microscopy with Immunofluorescence and Histological Stains (WB-ExM-IF & WB-ExM-Histo) V.(dr8d59s6)

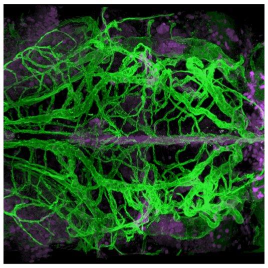

RESERVED DOI:
10.17504/protocols.io.dm6gp9wxjvzp/v1

Virginia M S Ruetten[1], Amy Hu[2], Mark Eddison[2], Kari Close[2], Yisheng He[2], Misha B. Ahrens[2], Paul Tillberg[2]

[1]Janelia Research Campus, HHMI - Gatsby Computational Neuroscience Unit; [2]HHMI/Janelia Research Campus

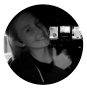

Virginia M S Ruetten
Janelia Research Campus, HHMI - Gatsby Computational Neurosc...

**Protocol Info:** Virginia M S Ruetten, Amy Hu, Mark Eddison, Kari Close, Yisheng He, Misha B. Ahrens, Paul Tillberg . Whole-Body Expansion Microscopy with Immunofluorescence and Histological Stains (WB-ExM-IF & WB-ExM-Histo). **protocols.io** **https://protocols.io/view/whole-body-expansion-microscopy-with-immunofluores-dr8d59s6**

**Created:** November 22, 2024

**Last Modified:** August 20, 2025

**Protocol Integer ID:** 112613

**Keywords:** Expansion Microscopy, Zebrafish, immunohistochemistry, clearing

#### Abstract

The interpretation of Whole Body Imaging (WBI) data necessitates comprehensive anatomical knowledge to accurately determine cell-type identity; however, resources pertaining to the internal anatomy of larval zebrafish are limited. To mitigate this gap, we established an advanced Whole Body Expansion-Microscopy (WB-ExM) protocol, facilitating the acquisition of molecular and cell-type information at subcellular resolution throughout the entire organism. While high-quality resources are available for the larval zebrafish brain[1,2,3] there remains a deficiency in materials concerning its visceral anatomy. In order to supplement existing histological, X-ray, and Expansion Microscopy methods aimed at mapping body-wide anatomy[4,5,6,7] and to develop a technique that yields high-fidelity molecular and anatomical data with enhanced processing speeds and compatibility with older samples, we devised an enzyme-free, rapid, and robust whole-body expansion microscopy method[8,9], effective on larvae up to at least 14 days post-fertilization. This method involves the use of high-temperature (100°C) chemical hydrolysis to uniformly soften tissues, embedding within a medium-density gel with reduced protein-gel anchoring, and repeated embedding post-digestion with moderate ~1.5x expansion factors in each cycle, culminating in a robust and uniform expansion even of challenging structures such as cartilage embedded in soft tissue. This protocol results in excellent optical clearing, retains high levels of antibody signals.
A demo dataset can be viewed **here**. Dorsal view of a double transgenic zebrafish animal labeling the ventricular and vascular systems (*Tg(foxj1a:eGFP)* x *Tg(flk1:dsRed-CAAX)*), stained against eGFP (magenta) and dsRed (green) (10 days post-fertilization, expanded ~2×).

1. Kunst, M. *et al.* A Cellular-Resolution Atlas of the Larval Zebrafish Brain. *Neuron* 103, 21-38.e5 (2019).
2. Randlett, O. *et al.* Whole-brain activity mapping onto a zebrafish brain atlas. *Nat Methods* 12, 1039–1046 (2015).
3. Tabor, K. M. *et al.* Brain-wide cellular resolution imaging of Cre transgenic zebrafish lines for functional circuit-mapping.
4. Copper, J. E. *et al.* Comparative analysis of fixation and embedding techniques for optimized histological preparation of zebrafish. *Comparative Biochemistry and Physiology Part C: Toxicology & Pharmacology* 208, 38–46 (2018).
5. Ding, Y. *et al.* Computational 3D histological phenotyping of whole zebrafish by X-ray histotomography. *eLife* 8, e44898 (2019).
6. Steib, E. *et al.* TissUExM enables quantitative ultrastructural analysis in whole vertebrate embryos by expansion microscopy. *Cell Reports Methods* 2, 100311 (2022).
7. Sim, J. *et al.* Nanoscale resolution imaging of the whole mouse embryos and larval zebrafish using expansion microscopy. Preprint at https://doi.org/10.1101/2021.05.18.443629 (2021).
8. Chen, F., Tillberg, P. W. & Boyden, E. S. Expansion microscopy.
9. Wang, Y. *et al.* EASI-FISH for thick tissue defines lateral hypothalamus spatio-molecular organization. *Cell* 184, 6361-6377.e24 (2021).

#### Image Attribution

Virginia M. S. Ruetten

#### Protocol materials

- CS-8R Coverslips, 0.15 mm (0.006 in), 8 mm diameter, pkg of 100 **Multi Channel Systems MCS GmbH Catalog #**640701
- Z1 Sample Holder **Janelia Research Campus**
- Alexa Fluor™ 488 NHS Ester (Succinimidyl Ester) **Thermo Fisher Scientific Catalog #**A20000
- ATTO 647N maleimide **AAT Bioquest Catalog #**2857
- DMSO, Anhydrous **Thermo Fisher Catalog #**D12345
- Acryloyl-X, SE (6-((acryloyl)amino)hexanoic acid, succinimidyl ester) **Thermo Fisher Scientific Catalog #**A20770
- DMSO, Anhydrous **Thermo Fisher Catalog #**D12345
- PBS - Phosphate-Buffered Saline (10X) pH 7.4, RNase-free **Thermo Fisher Scientific Catalog #**AM9625
- Pierce™ 16% Formaldehyde (w/v), Methanol-free **Thermo Scientific Catalog #**28906
- UltraPure™ DNase/RNase-Free Distilled Water **Thermo Fisher Scientific Catalog #**10977023
- GFP Polyclonal Antibody **Invitrogen - Thermo Fisher Catalog #**A-11122
- anti RFP antibody **Synaptic Systems Catalog #**409 006
- Anti-Rabbit-IgG - Atto 647N **Merck MilliporeSigma (Sigma-Aldrich) Catalog #**40839-1ML-F
- Donkey anti-Chicken IgY (H L) Highly Cross Adsorbed Secondary Antibody, Alexa Fluor™ 568 **Thermo Fisher Scientific Catalog #**A78950
- Corning® 25×25 mm Square #2 Cover Glass **Corning Catalog #**2855-25
- Poly-L-lysine hydrobromide **Merck MilliporeSigma (Sigma-Aldrich) Catalog #**Poly-L-lysine hydrobromide
- Photo Flo 200 Solution **Electron Microscopy Sciences Catalog #**74257
- Press-to-Seal™ Silicone Isolator with Adhesive, eight wells, 9 mm diameter, 0.5 mm deep **Thermo Fisher Scientific Catalog #**P24743
- Fisherbrand™ Superfrost™ Disposable Microscope Slides **Fisher Scientific Catalog #**Catalog No.12-550-123
- Alexa Fluor™ 488 NHS Ester (Succinimidyl Ester) **Thermo Fisher Scientific Catalog #**A20000
- ATTO 647N maleimide **AAT Bioquest Catalog #**2857
- NaCl (5 M), RNase-free **Thermo Fisher Scientific Catalog #**AM9760G
- UltraPure™ SDS Solution, 10% **Thermo Fisher Scientific Catalog #**15553027
- UltraPure™ DNase/RNase-Free Distilled Water **Thermo Fisher Scientific Catalog #**10977023
- PBS, pH 7.4 **Thermo Fisher Catalog #**10010001
- Triton™ X-100 **Merck MilliporeSigma (Sigma-Aldrich) Catalog #**X100-5ML
- Sodium Azide **Merck MilliporeSigma (Sigma-Aldrich) Catalog #**S2002-100G
- Normal Goat Serum **Jackson ImmunoResearch Laboratories, Inc. Catalog #**005-000-121

- Agarose, low gelling temperature **Merck MilliporeSigma (Sigma-Aldrich) Catalog #**A9414-100G
- PBS - Phosphate-Buffered Saline (10X) pH 7.4, RNase-free **Thermo Fisher Scientific Catalog #**AM9625
- Fisherbrand™ Superfrost™ Disposable Microscope Slides **Fisher Scientific Catalog #**Catalog No.12-550-123
- Press-to-Seal™ Silicone Isolator with Adhesive, eight wells, 9 mm diameter, 0.5 mm deep **Thermo Fisher Scientific Catalog #**P24743
- Hydrogen peroxide solution **Merck MilliporeSigma (Sigma-Aldrich) Catalog #**H1009-500ML
- PBS, pH 7.4 **Thermo Fisher Catalog #**10010001
- Triton™ X-100 **Merck MilliporeSigma (Sigma-Aldrich) Catalog #**X100-5ML
- Ammonium persulfate (APS) **Merck MilliporeSigma (Sigma-Aldrich) Catalog #**A3678-100G
- 40% Acrylamide Solution **Bio-Rad Laboratories Catalog #**1610140
- 2% bis-acrylamide solution **Bio-Rad Laboratories Catalog #**1610142
- Acrylic acid **Merck MilliporeSigma (Sigma-Aldrich) Catalog #**147230-5G
- UltraPure™ DNase/RNase-Free Distilled Water **Thermo Fisher Scientific Catalog #**10977023
- Sodium Hydroxide Solution (10N/Certified) **Fisher Scientific Catalog #**SS255-1
- 4-Hydroxy-TEMPO (4HT) **Merck MilliporeSigma (Sigma-Aldrich) Catalog #**4-Hydroxy-TEMPO
- N,N,N′,N′-Tetramethylethylenediamine (TEMED) **Merck MilliporeSigma (Sigma-Aldrich) Catalog #**T7024-25ML
- Acrylic acid **Merck MilliporeSigma (Sigma-Aldrich) Catalog #**147230-5G
- Sodium Hydroxide Solution (10N/Certified) **Fisher Scientific Catalog #**SS255-1
- UltraPure™ DNase/RNase-Free Distilled Water **Thermo Fisher Scientific Catalog #**10977023
- 2% bis-acrylamide solution **Bio-Rad Laboratories Catalog #**1610142
- Acrylic acid **Merck MilliporeSigma (Sigma-Aldrich) Catalog #**147230-5G
- UltraPure™ DNase/RNase-Free Distilled Water **Thermo Fisher Scientific Catalog #**10977023
- 40% Acrylamide Solution **Bio-Rad Laboratories Catalog #**1610140
- Sodium Hydroxide Solution (10N/Certified) **Fisher Scientific Catalog #**SS255-1
- Ammonium persulfate (APS) **Merck MilliporeSigma (Sigma-Aldrich) Catalog #**A3678-100G
- 4-Hydroxy-TEMPO (4HT) **Merck MilliporeSigma (Sigma-Aldrich) Catalog #**4-Hydroxy-TEMPO
- N,N,N′,N′-Tetramethylethylenediamine (TEMED) **Merck MilliporeSigma (Sigma-Aldrich) Catalog #**T7024-25ML
- Hydrogen peroxide solution **Merck MilliporeSigma (Sigma-Aldrich) Catalog #**H1009-500ML

#### Before start

Make sure you have all the reagents at hand.

#### Reagents

1 **4% PFA**

Cannot be prepared in advance.

Take a new ampule of 16% PFA (10ml). Aliquot it into 1 ml aliquots. Store at -80ºC. On the day take one aliquot out and thaw.

To make a total volume of 4 ml of 4% PFA: mix 1ml of 16% PFA, 0.4 ml of 10x PBS and 2.6 ml of distilled H20. Use within the next two days and store in a 4ºC fridge.

> Note
>
> PFA deteriorates even if stored at -80ºC. Avoid reusing aliquoted samples.
> When PFA is stored in sealed ampules, it is protected from atmospheric oxygen and moisture. This isolation prevents oxidative degradation

**Reagents**

- PBS - Phosphate-Buffered Saline (10X) pH 7.4, RNase-free **Thermo Fisher Scientific Catalog** #AM9625
- Pierce™ 16% Formaldehyde (w/v), Methanol-free **Thermo Scientific Catalog** #28906
- UltraPure™ DNase/RNase-Free Distilled Water **Thermo Fisher Scientific Catalog** #10977023

1 **Blocking Buffer**

Cannot be prepared in advance.

5% goat serum, 0.5% Triton-100 in 1x PBS, 0.1% Na Azide.

Keep in fridge at 4ºC

4 °C

To make 7.5 ml of Blocking Buffer

- 0.375 ml of goat serum
- 0.75 ml of PBS-T-5-NaAz-1 [PBS + Triton (5%) + NaAzide (1%)]
- 6.375 ml of 1x PBS

> Note
>
> Notes on Goat Serum:
> This comes in a sealed vial. Do not open the seal but use a syringe to extract medium. The medium is nutrient rich and so easily can become contaminated. Do not use for more than 3 months. Reconstitute with distilled water.

**Reagents**

- Normal Goat Serum **Jackson ImmunoResearch Laboratories, Inc. Catalog** #005-000-121
- PBS, pH 7.4 **Thermo Fisher Catalog** #10010001

Triton™ X-100 **Merck MilliporeSigma (Sigma-Aldrich) Catalog #**X100-5ML

Sodium Azide **Merck MilliporeSigma (Sigma-Aldrich) Catalog #**S2002-100G

2 **Antibodies**

**Primary antibodies (polyclonal)**

rabbit anti-eGFP

chicken anti-RFP

anti-mRuby

**Secondary antibodies**

goat anti-rabbit Atto647N

donkey anti-chicken Alexa568

GFP Polyclonal Antibody **Invitrogen - Thermo Fisher Catalog #**A-11122

anti RFP antibody **Synaptic Systems Catalog #**409 006

Anti-Rabbit-IgG - Atto 647N **Merck MilliporeSigma (Sigma-Aldrich) Catalog #**40839-1ML-F

Donkey anti-Chicken IgY (H L) Highly Cross Adsorbed Secondary Antibody, Alexa Fluor™ 568 **Thermo Fisher Scientific Catalog #**A78950

3 **PBST-0.5**

Can be prepared in advance and stored at room temperature.

1x PBS with 0.5% or 0.1% Triton

PBST-0.5 and PBST-0.1 respectively

Triton dissolves terribly. It hardens upon making contact with water and takes time to dissolve fully. Prepare a 10% stock solution in 1x PBS and use that for subsequent rounds.

Keep at RT.

Room temperature

**Reagents**

PBS, pH 7.4 **Thermo Fisher Catalog #**10010001

Triton™ X-100 **Merck MilliporeSigma (Sigma-Aldrich) Catalog #**X100-5ML

4 **Bleaching Solution**

Cannot be prepared in advance.

Combine 3% $H_2O_2$, 60mM KOH and 100mM (or more) NaN3 in PBS.

- Prepare 300mM NaN3: add 195mg of NaN3 to 10ml of PBS.

To make 5 ml of bleaching solution:

- 0.3 ml of 50% H2O2 solution
- 0.15 ml of 1M KOH solution (takes one hour with very little bubbling)
- fill the rest of 100mM NaN3 (i.e.: 4.25ml, 85mM NaN3)

Note

Many biological tissues contain catalase, which rapidly breaks down hydrogen peroxide into water and oxygen. This leads to a large number of bubbles, which can rupture the tissue. The tissue should first be passivated. NaN3 is a catalase inhibitor. Pre-incubate fish with NaN3 to block catalase. Then add bleach solution containing NaN3. The dissociation of H2O2 occurs faster at high pH, thus higher KOH should've used.

Hydrogen peroxide solution **Merck MilliporeSigma (Sigma-Aldrich) Catalog #**H1009-500ML

5 **1% low-melting temperature agarose**

Can be prepared in advance.

Dissolve 1 g of low-melting point agarose in 100 ml of 1x PBS.

Add 1 g of power to 100 ml 1x PBS at RT. Stir with magnetic stirrer.

Bring to a boil using a microwave. Stir with magnetic stirrer until full dissolved.

Repeat boil and stirring until solution is clear.

**Reagents**

Agarose, low gelling temperature **Merck MilliporeSigma (Sigma-Aldrich) Catalog #**A9414-100G

PBS - Phosphate-Buffered Saline (10X) pH 7.4, RNase-free **Thermo Fisher Scientific Catalog #**AM9625

6 **Acryloyl-X Solution (AcX)**

Cannot be prepared in advance.

Stock concentration: 10 mg/ml

Dissolve to 10 mg/ml in anhydrous DMSO. Aliquot in 20 µl batches.

Store in a desiccated environment at -20°C. Don't re-use AcX after thawing.

Working solution will be: 20 µg/ml, dilution 1:500 in 1x PBS

**Reagents**

Acryloyl-X, SE (6-((acryloyl)amino)hexanoic acid, succinimidyl ester) **Thermo Fisher Scientific Catalog #**A20770

DMSO, Anhydrous **Thermo Fisher Catalog #**D12345

7 **Na Acrylate Solution**

Fill a 500 ml beaker with ~300 ml of water for a water bath.

Place an open 50 ml Eppendorf tube into the bath and bring the setup to a fume hood.

The purpose of the water bath is to provide a highly conductive medium to cool the solution.

Add 9.0 ml of water to the Eppendorf tube.

Add 11 ml of acrylic acid to the Eppendorf tube. (Note that this is a flammable and reactive compound).

Add 14.4 ml of 10 M NaOH to the Eppendorf tube. This should be done dropwise to prevent excessive heating and boiling.

A yellow precipitate should be observable. Leave it to cool.

Calibrate a pH meter.
Remove the acrylate-filled tube from the fume hood.
By now, most of the acrylic acid will have been converted to non-volatile sodium acrylate.
Measure the pH of the solution.
Add NaOH to adjust the pH gradually to between 7.5-8 using a pH meter. Do NOT use pH test strips.
We recommend starting by adding 10 M NaOH solution, and when the pH gets close to the desired pH adjusting via the use of 1 M NaOH solution.
(As a general guidance: Add about 500 µl 10 M NaOH, 250 µl 10 M NaOH, 120 µL 10 M NaOH, then some 1 M NaOH)
Add water up to a final volume of 40 ml.
(Note: Acrylic acid has a pKa of 4.76 at pH 7.75 - this solution has about 4 mM remaining buffering capacity)

Acrylic acid **Merck MilliporeSigma (Sigma-Aldrich) Catalog #**147230-5G

Sodium Hydroxide Solution (10N/Certified) **Fisher Scientific Catalog #**SS255-1

UltraPure™ DNase/RNase-Free Distilled Water **Thermo Fisher Scientific Catalog #**10977023

8 **Monomer and Gelation Solutions #1 (Medium Density Gel with high Bis)**

| A | B | C | D | E | F |
|---|---|---|---|---|---|
| name | units | stock conc. | final conc. | vol (ul) | vol (ml) |
| Acrylamide | % | 40 | 10 | 250 | 2.5 |
| Na Acrylate | M | 4 | 0.5 | 125 | 1.25 |
| Bis | % | 1 | 0.1 | 100 | 1 |
| 10x PBS | x | 10 | 1 | 100 | 1 |
| Water | | | | 365 | 3.65 |

**Monomer Solution #1**

| A | B | C | D | E |
|---|---|---|---|---|
| name (units) | units | stock conc. | final conc. | vol (ul) |
| Monomer Solution #1 | | | | 940 |
| APS | % | 10 | 0.2 | 20 |
| TEMED | % | 10 | 0.2 | 20 |
| 4HT | % | 0.5 | 0.01 | 20 |

**Gelation Solution #1**

**Reagents**

40% Acrylamide Solution **Bio-Rad Laboratories Catalog #**1610140

2% bis-acrylamide solution **Bio-Rad Laboratories Catalog** #1610142

Acrylic acid **Merck MilliporeSigma (Sigma-Aldrich) Catalog** #147230-5G

Sodium Hydroxide Solution (10N/Certified) **Fisher Scientific Catalog** #SS255-1

UltraPure™ DNase/RNase-Free Distilled Water **Thermo Fisher Scientific Catalog** #10977023

Ammonium persulfate (APS) **Merck MilliporeSigma (Sigma-Aldrich) Catalog** #A3678-100G

4-Hydroxy-TEMPO (4HT) **Merck MilliporeSigma (Sigma-Aldrich) Catalog** #4-Hydroxy-TEMPO

N,N,N′,N′-Tetramethylethylenediamine (TEMED) **Merck MilliporeSigma (Sigma-Aldrich) Catalog** #T7024-25ML

9 **Monomer and Gelation Solutions #2 (Medium Density Gel with low Bis)**

| A | B | C | D | E | F |
|---|---|---|---|---|---|
| | | | | to make 1ml | to make 10ml |
| component | units | stock conc. | final conc. | vol (ul) | vol (ml) |
| Acrylamide | % | 40 | 10 | 250 | 2.5 |
| Na Acrylate | M | 4 | 0.5 | 125 | 1.25 |
| Bis | % | 1 | 0.02 | 20 | 0.2 |
| 10x PBS | x | 10 | 1 | 100 | 1 |
| Water | | | | 445 | 4.45 |

**Monomer Solution #2**

| A | B | C | D | E |
|---|---|---|---|---|
| | | | | to make 1ml |
| component | units | stock conc. | final conc. | vol (ul) |
| Monomer Solution #2 | | | | 940 |
| APS | % | 10 | 0.2 | 20 |
| TEMED | % | 10 | 0.2 | 20 |
| 4HT | % | 0.5 | 0.01 | 20 |

**Gelation Solution #2**

**Reagents**

40% Acrylamide Solution **Bio-Rad Laboratories Catalog** #1610140

2% bis-acrylamide solution **Bio-Rad Laboratories Catalog** #1610142

Acrylic acid **Merck MilliporeSigma (Sigma-Aldrich) Catalog** #147230-5G

Sodium Hydroxide Solution (10N/Certified) **Fisher Scientific Catalog** #SS255-1

UltraPure™ DNase/RNase-Free Distilled Water **Thermo Fisher Scientific Catalog** #10977023

Ammonium persulfate (APS) **Merck MilliporeSigma (Sigma-Aldrich) Catalog** #A3678-100G

4-Hydroxy-TEMPO (4HT) **Merck MilliporeSigma (Sigma-Aldrich) Catalog** #4-Hydroxy-TEMPO

N,N,N′,N′-Tetramethylethylenediamine (TEMED) **Merck MilliporeSigma (Sigma-Aldrich) Catalog** #T7024-25ML

10 **Disruption Buffer**

| A | B | C | D | E | F |
|---|---|---|---|---|---|
| | | | | to make 1ml | to make 100ml |
| component | units | stock conc. | final conc. | vol (ul) | vol |
| SDS | % | 10 | 5 | 0.5 | 50 |
| Tris pH 7.5 | mM | 1000 | 50 | 0.05 | 5 |
| NaCl | M | 5 | 0.2 | 0.04 | 4 |
| MilliQ water | | | | 0.41 | 41 |

**Disruption Buffer**

Stock **Disruption Buffer** can be prepared in advanced and stored at RT.

NaCl (5 M), RNase-free **Thermo Fisher Scientific Catalog** #AM9760G

UltraPure™ SDS Solution, 10% **Thermo Fisher Scientific Catalog** #15553027

UltraPure™ DNase/RNase-Free Distilled Water **Thermo Fisher Scientific Catalog** #10977023

11 **Total Protein Stains**

Prepare stock: 10 mg/ml stocks in anhydrous DMSO.
Aliquot in 5-10ul.
Store in a desiccated environment at -20°C.

**Reagents**

Alexa Fluor™ 488 NHS Ester (Succinimidyl Ester) **Thermo Fisher Scientific Catalog** #A20000

ATTO 647N maleimide **AAT Bioquest Catalog** #2857

DMSO, Anhydrous **Thermo Fisher Catalog** #D12345

12 **Gelation Chambers**

Silicone gaskets (Invitrogen P24743)

Glass slides (SuperFrost 12550123)

Scotch tape

Poly-l-lysine

Shaker (nutator - 75 RPM)

**Reagents**

Press-to-Seal™ Silicone Isolator with Adhesive, eight wells, 9 mm diameter, 0.5 mm deep **Thermo Fisher Scientific Catalog #**P24743

Fisherbrand™ Superfrost™ Disposable Microscope Slides **Fisher Scientific Catalog #**Catalog No.12-550-123

Poly-L-lysine hydrobromide **Merck MilliporeSigma (Sigma-Aldrich) Catalog #**Poly-L-lysine hydrobromide

Photo Flo 200 Solution **Electron Microscopy Sciences Catalog #**74257

13 **General note for sample handling**

Soak plastic transfer pipettes in PBST-0.5 before manipulating samples - otherwise, the fish will inevitably get stuck.

Never use forceps to handle or move the fish as this inevitably damages the sample.

#### Fixation and Permeabilization

10h

14 Euthanize samples with an overdose of MS-222 (a.k.a. tricaine) (200-300 mg/L).

15 Prepare fresh 4% PFA.

16 Place samples in 1 ml of 4% PFA.

Room temperature

17 Keep overnight in 4% PFA at 4°C on a shaker.

9h

09:00:00

4 °C

18 Rinse samples in 4 × 15 min 1 ml 1x PBS.

1h

01:00:00

Room temperature

#### Bleaching

19 Prepare fresh **Bleaching Solution**

Combine 3% $H_2O_2$, 0.5% KOH (~90mM) and 50mM (or more) NaN3 in PBS.

Prepare 100mM NaN3:
Add 65mg of NaN3 to 10ml of PBS.
To make 5 ml of Bleaching Solution:
- 0.3 ml of 50% H2O2 solution
- 0.45 ml of 1M KOH solution
- fill the rest of 100mM NaN3 (i.e.: 4.25ml) to get to 85mM of NaN30.5% KOH (~90mM) and 50mM (or more) NaN3 in PBS.

Hydrogen peroxide solution **Merck MilliporeSigma (Sigma-Aldrich) Catalog #**H1009-500ML

Hydrogen peroxide solution **Merck MilliporeSigma (Sigma-Aldrich) Catalog #**H1009-500ML

20 Preincubate the samples in 100mM NaN3 (~10mins).
Incubate the samples in 500 µL of **Bleaching Solution**. Very little bubbling should occur.

Room temperature Room temperature

21 Rinse the samples twice with 1x PBS.

Room temperature

#### Immunofluorescence (Optional)

5d 17h 10m

22 Prepare fresh **Blocking Buffer**.

23 Incubate the samples in **Blocking Buffer** (3 hr at RT).

3h

Room temperature

03:00:00

24 Incubate the samples in primary antibodies diluted 1:100 in **Blocking Buffer** in PCR tube (3 days at RT) on a shaker. Make sure Na Azide is included in the **Blocking Buffer** to prevent microbial growth. 3d

72:00:00

Room temperature

Note

**Can I place multiple samples in the same PCR tube?**
Yes, multiple samples can be placed in the same PCR tube (~10). The level of antibody is high enough that it should not get significantly depleted.

**What antibodies can I use?**
As we do IHC pre-expansion and digestion, any antibody that works well in fixed whole-mount preparation should work in expansion.

25 Wash sample in 1 ml **Blocking Buffer** (4 × 15 min at RT) on a shaker. 1h

Room temperature

01:00:00

26 Wash the samples in **Blocking Buffer** (3 × 2 hr or overnight at RT) on a shaker. 9h

Room temperature

09:00:00

27 Incubate the samples in secondary antibodies (1:100) in **Blocking Buffer** in PCR tube (2 days at RT) on a shaker. 2d

48:00:00

Room temperature

28 Wash the samples in **Blocking Buffer** (3× 2 hr, at RT). 4h

Room temperature

04:00:00

29 Wash the samples in 1x PBS (2× 5 min at RT) and transfer to fresh 1x PBS. 10m

Room temperature

00:10:00

30 Image one sample to confirm immunofluorescence staining was successful.

#### Permeabilization (if no IHC)

5h 10m

31 For specimens that have not been stained with antibodies, permeabilize (at least 5 hr or overnight) in **PBST-0.5**. 5h

05:00:00

32 Wash samples (2× 5 min at RT) in 1x PBS. 10m

Room temperature

00:10:00

#### Agarose Embedding

33 Heat agarose in microwave until it liquifies.
Cool agarose to 50°C (by placing in 50°C incubator).

34 Adhere silicone gasket (Invitrogen P24743 or Invitrogen P24740) on a glass slide (Superfrost Microscope Slides #12550123) to form a **Mounting Chamber**.

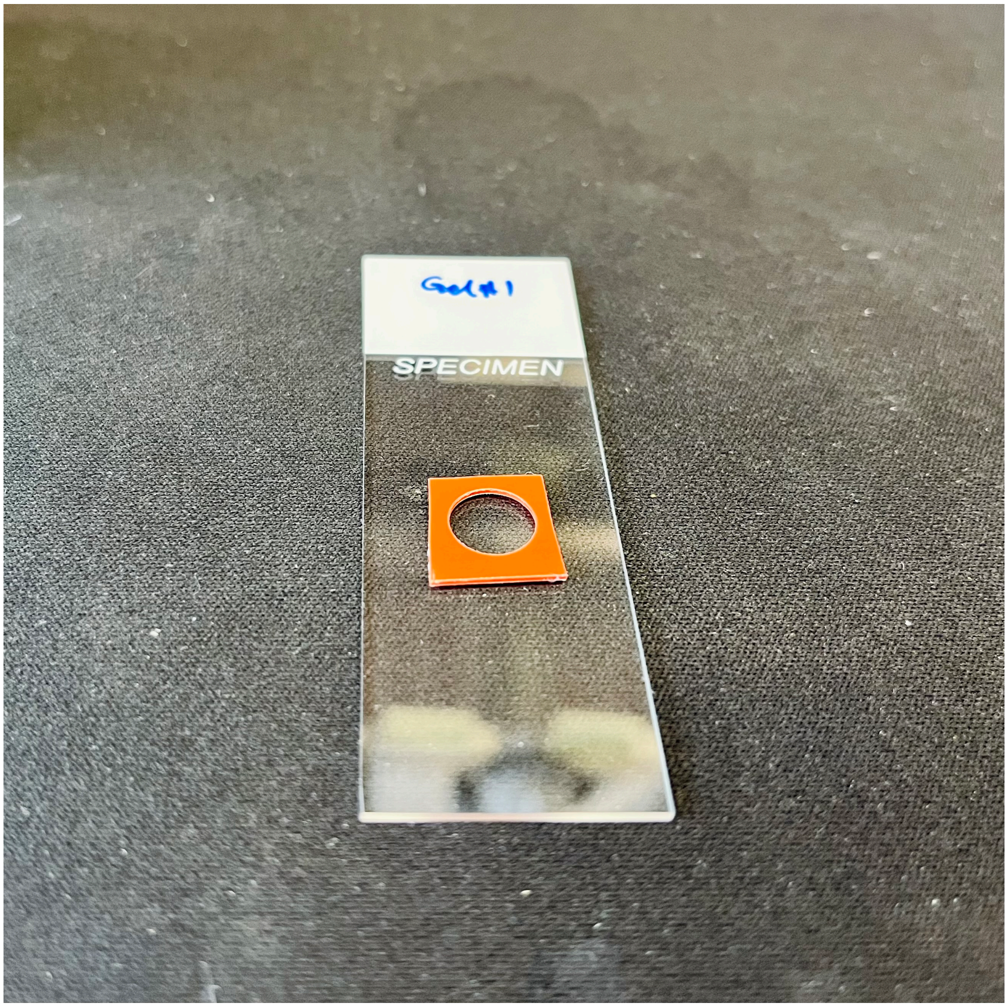

Mounting Chamber

**Reagents**

Press-to-Seal™ Silicone Isolator with Adhesive, eight wells, 9 mm diameter, 0.5 mm deep **Thermo Fisher Scientific Catalog** #P24743

Fisherbrand™ Superfrost™ Disposable Microscope Slides **Fisher Scientific Catalog** #Catalog No.12-550-123

35 Place fish in **Mounting Chamber**, remove any access PBS, and cover with ~70 µl of 1% low-melting-point agarose. Orientate as desired (on side or dorsal side up) and leave to solidify.
Additional notes:
Use a sharp and clean utensil to nudge the fish into the desired orientation.
If more time is needed, the mounting chamber can be placed on a heating block to ensure that the agarose doesn't solidify pre-emptively.
Always begin by nudging the fish to the very bottom of the agarose drop to avoid the sample being held by surface tension to the top of the drop. Make sure that the fish is fully submerged in agarose.

36 With a razor blade, cut out a rectangle around the agarose-embedded sample and delicately lift the sample and transfer it to 12 well-plate with 1 ml of 1x PBS/well.
Cut as close to the fish as possible with some safety margin.

#### Protein Anchoring

1h

37 Prepare **Acryloyl-X Solution** (stock solution 10 mg/ml, working solution: 20 µg/ml, dilution 1:500 in 1x PBS).
1 ml per sample is needed.
Do not prepare in advance.

38 Incubate each sample in 1 ml of **Acryloyl-X Solution** (1 hr at RT) shaking in 12-well plate. 1h

Room temperature

01:00:00

#### Preparation of Gelation Chamber #1

39 Prepare chambers for gelation, one for each sample.
Layer 11 pieces of Scotch tape together to create ~0.6 mm thick spacers.
Cut two strips of spacer material, about 2 cm long and 0.5 cm wide.
Stick two strips to a glass slide ~15 mm apart to form the side walls of the **Gelation Chamber #1**.

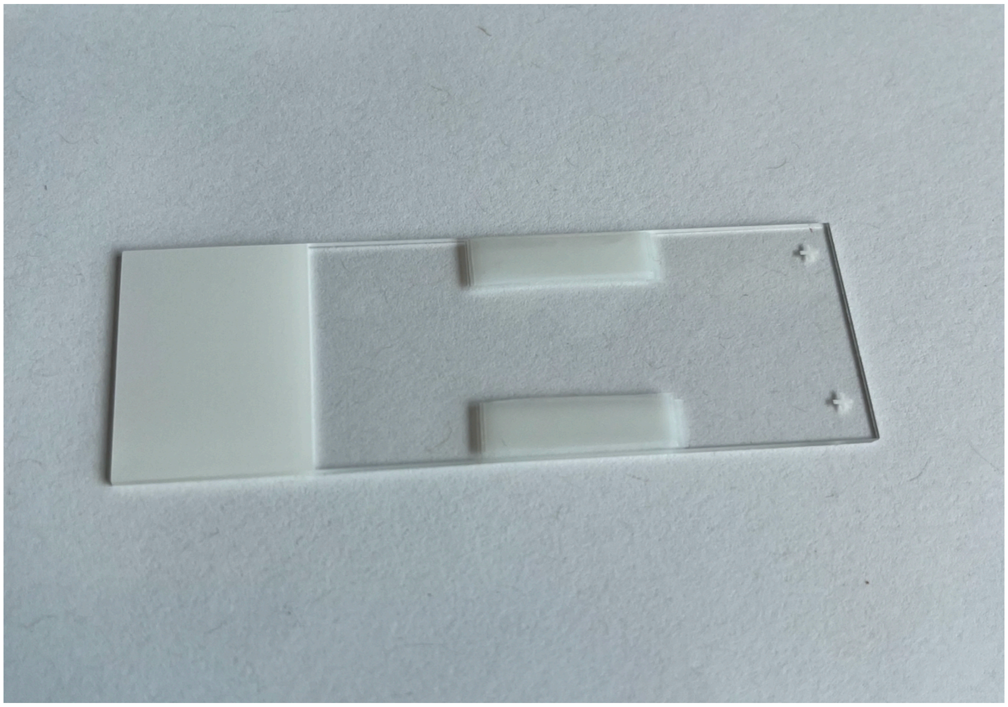

Gelation Chamber #1

#### Incubation with Gelation Solution 1

45m

40 Rinse samples with 1x PBS (3× 5 min).

00:15:00

15m

41 Thaw **Monomer Solution #1**, as well as 4HT, TEMED and APS. Vortex well and keep on ice.

42 Mix **Monomer Solution #1** and 4HT, TEMED and APS at a ratio of 94:2:2:2 to produce **Gelation Solution #1**.
Vortex.
Each sample needs ~3 ml of **Gelation Solution #1**..
~1 ml per incubation round.

43 Incubate samples in **Gelation Solution #1** on ice (3× 10 min at 4ºC) with 1 ml of **Gelation Solution #1** on a shaker in a 12-well plate.

00:30:00

30m

#### Gelation 1

2h

44 With spatula, transfer agarose block containing fish into the **Gelation Chamber #1**.

45 Gently place cover slip over the agarose block lying on the walls of **Gelation Chamber #1** (made of scotch tape).
The cover slip should lay flat on the agarose block.
Gently press to seal.

46 Slowly pipette **Gelation Solution #1** into **Gelation Chamber #1** until full. Avoid any air bubbles.
Solution will hold by water tension.
One can use **Gelation Solution #1** that was used during the previous incubation (no need to prepare fresh one).

47 Once **Gelation Chamber #1** is filled, place the chambers at 37ºC for 2 hr to induce polymerization. 2h
Ensure incubator is humidified.
37 °C
02:00:00

#### Disruption 9h 5m

48 Turn on the 100ºC heat block (for later disruption step).
Prepare **Disruption Buffer** (5 ml/sample).
Stock **Disruption Buffer** can be prepared in advanced and stored at RT (e.g.: 50-100 ml stock).

49 Place gels at RT, and allow them to cool on the bench for > 5 min. 5m
Room temperature
00:05:00

50 Take off coverslip lid with a razor blade.
Under a stereomicroscope, trim the gels into a rectangle with a ~2-3 mm border on either side of the sample.
Add a nick on the top right corner to be able to track orientation of the samples.
The gel should be about 9 mm x 4 mm.

51 Transfer each gel to a 2 ml Eppendorf tube and add 1.8 ml of **Disruption Buffer.**

52 Incubate each samples in **Disruption Buffer** (overnight at 100ºC). 9h
Make sure that lid is closed tightly to avoid evaporation.
100 °C
09:00:00
After disruption, the gel should be about 12.5 mm x 6 mm.

#### Chamber 2 preparation

53 Prepare **Gelation Chamber #2**, one for each sample.
Layer 20 pieces of Scotch tape together to create ~1.2 mm thick spacers.
Cut two strips of spacer material, about 2 cm long and 0.5 cm wide.
Stick two strips to a glass slide ~15 mm apart to form the side walls of the **Gelation Chamber #2**.

#### Incubation with Gelation Solution 2

1h 30m

54 Wash each samples in 1x PBS at RT (3× 20 min).

1h

Room temperature

01:00:00

55 Thaw **Monomer Solution #2,** as well as 4HT, TEMED and APS. Vortex well and keep on ice.

56 Mix **Monomer Solution #2** and 4HT, TEMED and APS at a ratio of 94:2:2:2 to produce **Gelation Solution #2**.
Vortex.
Each sample needs ~6 ml of **Gelation Solution #2**.
~2 ml per incubation round.

57 Incubate samples in **Gelation Solution #2** on ice (3× 10 mins @ 4°C) with 2ml of **Gelation Solution #2** on a shaker in 12-well plate.

30m

4 °C

00:30:00

#### Gelation 2

2h 5m

58 With spatula, transfer gel block containing fish into a **Gelation Chamber #2**.
Make sure there are no bubbles at the interface between chamber and gel.
If needed, add a small drop of gelation solution below the gel if air bubbles remain.
Add a small drop of gelation solution on top of the gel to minimize chances of air bubbles forming.
Carefully place coverslip on top (Corning #2855-25)

**Reagents**

Corning® 25×25 mm Square #2 Cover Glass **Corning Catalog** #2855-25

59 Slowly pipette fresh **Gelation Solution #2** into **Gelation Chamber #2** until full. Avoid any air bubbles on either side.
Solution will hold by water tension.
Wipe off any excess solution with a wipe.
The excess solution will prevent oxygen from reaching the gel too quickly, and avoid the inhibition of polymerization.

60 Once **Gelation Chamber #2** is filled, place the chambers at 37 ºC for 2 hr to induce polymerization.
Ensure incubator is humidified.

2h

37 °C

02:00:00

61 Place gels at RT, and allow them to cool on the bench for > 5 min.

5m

Room temperature

00:05:00

62 Take off coverslip lid with a razor blade.
Under a stereomicroscope, remove the scotch side walls and trim the gels into a rectangle, close to the first gel.
Removing any second gel material that formed outside of the first gel.

63 Transfer gel to a 12-well plate with 1x PBS.
Transfer each gel to a 5 ml Eppendorf tube and fill with **Disruption Buffer.**
Incubate overnight at 95ºC.
Make sure that lid is closed tightly to avoid evaporation.

100 °C

09:00:00

#### Staining

2h 30m

64 Incubate gels in Alexa488-NHS and Atto647N-maleimide dye at 1:1000 each in 1x PBS (1 hr at RT) in 12-well plate on gentle shaker.

1h

Room temperature

01:00:00

**Reagents**

Alexa Fluor™ 488 NHS Ester (Succinimidyl Ester) **Thermo Fisher Scientific Catalog** #A20000

ATTO 647N maleimide **AAT Bioquest Catalog** #2857

65 Wash in 1x PBS (3× 1 hr) on shaker at RT.

1h 30m

Room temperature

01:30:00

#### Mounting

66 Attach a 8mm circular coverslips (CS-8R, Warner Instruments, 64-0701) to a Z1 sample holder using superglue.

**Reagents**

CS-8R Coverslips, 0.15 mm (0.006 in), 8 mm diameter, pkg of 100 **Multi Channel Systems MCS GmbH Catalog** #640701

Z1 Sample Holder **Janelia Research Campus**

LIGHTSHEETHOLDER V13 Short.ipt LIGHTSHEETHOLDER V13 Short.stl

67 Coat coverslip with poly-lysine and let it dry.

68 Mount gel, sample side up.

#### Imaging

69 Image using a Zeiss Z1 Light sheet Microscope. Allow 45 min (with gel in place) for system to equilibrate before acquiring multi-tile acquisitions.
Image with 20× 1.0 NA water immersion objective.
Exposure time: 100-150 ms.

70 After imaging, gels can be teased off holder with a paint brush.

71 For long term storage, keep gels in 1x PBS at 4ºC.

#### Protocol references

1. Kunst, M. *et al.* A Cellular-Resolution Atlas of the Larval Zebrafish Brain. *Neuron* 103, 21-38.e5 (2019).
2. Randlett, O. *et al.* Whole-brain activity mapping onto a zebrafish brain atlas. *Nat Methods* 12, 1039–1046 (2015).
3. Tabor, K. M. *et al.* Brain-wide cellular resolution imaging of Cre transgenic zebrafish lines for functional circuit-mapping.
4. Copper, J. E. *et al.* Comparative analysis of fixation and embedding techniques for optimized histological preparation of zebrafish. *Comparative Biochemistry and Physiology Part C: Toxicology & Pharmacology* 208, 38–46 (2018).
5. Ding, Y. *et al.* Computational 3D histological phenotyping of whole zebrafish by X-ray histotomography. *eLife* 8, e44898 (2019).
6. Steib, E. *et al.* TissUExM enables quantitative ultrastructural analysis in whole vertebrate embryos by expansion microscopy. *Cell Reports Methods* 2, 100311 (2022).
7. Sim, J. *et al.* Nanoscale resolution imaging of the whole mouse embryos and larval zebrafish using expansion microscopy. Preprint at https://doi.org/10.1101/2021.05.18.443629 (2021).
8. Chen, F., Tillberg, P. W. & Boyden, E. S. Expansion microscopy.
9. Wang, Y. *et al.* EASI-FISH for thick tissue defines lateral hypothalamus spatio-molecular organization. *Cell* 184, 6361-6377.e24 (2021).

#### Acknowledgements

This research was funded by Janelia Research Campus, HHMI and the Gatsby Computational Neuroscience Unit.

### Whole-Body Expansion Microscopy with Fluorescence Insitu Hybridization (WB-ExM FISH)

RESERVED DOI:
**10.17504/protocols.io.5qpvo9y89v4o/v1**

Virginia Ruetten[1], Mark Eddision[2], Amy Hu[2], Misha B. Ahrens[2], Paul Tillberg[2]

[1]Janelia Research Campus, HHMI - Gatsby Computational Neuroscience Unit; [2]HHMI/Janelia Research Campus

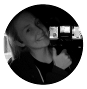

Virginia Ruetten
Janelia Research Campus, HHMI - Gatsby Computational Neurosc...

**Protocol Info:** Virginia Ruetten, Mark Eddision, Amy Hu, Misha B. Ahrens, Paul Tillberg . Whole-Body Expansion Microscopy with Fluorescence Insitu Hybridization (WB-ExM FISH). **protocols.io** **https://protocols.io/view/whole-body-expansion-microscopy-with-fluorescence-dr8859zw**

**Created:** November 22, 2024

**Last Modified:** March 10, 2025

**Protocol Integer ID:** 112640

**Keywords:** Expansion Microscopy, Zebrafish

#### Abstract

Whole-Body Expansion Microscopy with Fluorescence In-situ Hybridization for young zebrafish.

#### Protocol materials

- Poly-L-lysine hydrobromide **Merck MilliporeSigma (Sigma-Aldrich) Catalog #**Poly-L-lysine hydrobromide In 2 steps
- Acrylic acid **Merck MilliporeSigma (Sigma-Aldrich) Catalog #**147230-5G In 2 steps
- Triton™ X-100 **Merck MilliporeSigma (Sigma-Aldrich) Catalog #**X100-5ML Step 2
- ATTO 647N maleimide **AAT Bioquest Catalog #**2857 Step 9
- Melphalan **Cayman Chemical Company Catalog #**16665 Step 6
- Fisherbrand™ Superfrost™ Disposable Microscope Slides **Fisher Scientific Catalog #**Catalog No.12-550-123 Step 12
- DMSO, Anhydrous **Thermo Fisher Catalog #**D12345 In 3 steps
- 4-Hydroxy-TEMPO (4HT) **Merck MilliporeSigma (Sigma-Aldrich) Catalog #**4-Hydroxy-TEMPO In 2 steps
- Press-to-Seal™ Silicone Isolator with Adhesive, eight wells, 9 mm diameter, 0.5 mm deep **Thermo Fisher Scientific Catalog #**P24743 Step 12
- UltraPure™ DNase/RNase-Free Distilled Water **Thermo Fisher Scientific Catalog #**10977023 In 4 steps
- RNase-Free DNase Set **Qiagen Catalog #**79254 In 3 steps
- UltraPure™ SDS Solution, 10% **Thermo Fisher Scientific Catalog #**15553027 Step 11
- Ammonium persulfate (APS) **Merck MilliporeSigma (Sigma-Aldrich) Catalog #**A3678-100G In 2 steps
- PBS, pH 7.4 **Thermo Fisher Catalog #**10010001 Step 2
- CS-8R Coverslips, 0.15 mm (0.006 in), 8 mm diameter, pkg of 100 **Multi Channel Systems MCS GmbH Catalog #**640701 Step 72
- Alexa Fluor™ 488 NHS Ester (Succinimidyl Ester) **Thermo Fisher Scientific Catalog #**A20000 Step 9
- Proteinase K, Molecular Biology Grade **New England Biolabs Catalog #**P8107S Step 11
- EDTA (0.5 M), pH 8.0, RNase-free **Thermo Fisher Scientific Catalog #**AM9260G Step 11
- 40% Acrylamide Solution **Bio-Rad Laboratories Catalog #**1610140 In 2 steps
- Pierce™ 16% Formaldehyde (w/v), Methanol-free **Thermo Scientific Catalog #**28906 Step 1
- 2% bis-acrylamide solution **Bio-Rad Laboratories Catalog #**1610142 In 2 steps
- Z1 Sample Holder **Janelia Research Campus** Step 72
- Acryloyl-X, SE (6-((acryloyl)amino)hexanoic acid, succinimidyl ester) **Thermo Fisher Scientific Catalog #**A20770 Step 5
- Melphalan **Cayman Chemical Company Catalog #**16665 Step 4
- SSC (20X), RNase-free **Thermo Scientific Catalog #**AM9763 In 2 steps
- NaCl (5 M), RNase-free **Thermo Fisher Scientific Catalog #**AM9760G Step 11

Corning® 25x25 mm Square #2 Cover Glass **Corning Catalog #**2855-25 Step 47

MOPS (Fine White Crystals/Molecular Biology), Fisher BioReagents™ **Thermo Fisher Scientific Catalog #**BP308100

In 2 steps

Photo Flo 200 Solution **Electron Microscopy Sciences Catalog #**74257 In 2 steps

PBS - Phosphate-Buffered Saline (10X) pH 7.4, RNase-free **Thermo Fisher Scientific Catalog #**AM9625 Step 1

N,N,N′,N′-Tetramethylethylenediamine (TEMED) **Merck MilliporeSigma (Sigma-Aldrich) Catalog #**T7024-25ML

In 2 steps

Sodium Hydroxide Solution (10N/Certified) **Fisher Scientific Catalog #**SS255-1 In 2 steps

#### Before start

Make sure you have all the reagents at hand.

#### Reagents

1 **4% PFA**

Cannot be prepared in advance.

Take a new ampule of 16% PFA (10ml). Aliquot it into 1 ml aliquots. Store at -80°C. On the day take one aliquot out and thaw.

To make a total volume of 4 ml of 4% PFA: mix 1ml of 16% PFA, 0.4 ml of 10x PBS and 2.6 ml of distilled H20. Use within the next two days and store in a 4°C fridge.

> **Note**
>
> PFA deteriorates even if stored at -80°C. Avoid reusing aliquoted samples.
> When PFA is stored in sealed ampules, it is protected from atmospheric oxygen and moisture. This isolation prevents oxidative degradation

**Reagents**

- PBS - Phosphate-Buffered Saline (10X) pH 7.4, RNase-free **Thermo Fisher Scientific Catalog #**AM9625
- Pierce™ 16% Formaldehyde (w/v), Methanol-free **Thermo Scientific Catalog #**28906
- UltraPure™ DNase/RNase-Free Distilled Water **Thermo Fisher Scientific Catalog #**10977023

2 **PBST-0.5**

Can be prepared in advance.

1x PBS with 0.5% or 0.1% Triton

PBST-0.5 and PBST-0.1 respectively

Triton dissolves terribly. It hardens upon making contact with water and takes time to dissolve fully. Prepare a 10% stock solution in 1x PBS and use that for subsequent rounds.

Keep at RT.

Room temperature

**Reagents**

- PBS, pH 7.4 **Thermo Fisher Catalog #**10010001
- Triton™ X-100 **Merck MilliporeSigma (Sigma-Aldrich) Catalog #**X100-5ML

3 **MOPS Buffer**

Stock concentration: 200 mM (10x)

Dissolve 1046.5 mg in 25 ml in nuclease-free water, pH to 7.7 with **10N** NaOH.

Store at -20°C.

- MOPS (Fine White Crystals/Molecular Biology), Fisher BioReagents™ **Thermo Fisher Scientific Catalog #**BP308100
- UltraPure™ DNase/RNase-Free Distilled Water **Thermo Fisher Scientific Catalog #**10977023

4 **Melphalan stock**

Stock concentration: 2.5 mg/ml

Dissolve Melphalan (2.5 mg per ml) in anhydrous DMSO. To dissolve, heat to 37°C and vortex vigorously and place on a shaker. This may take an hour to dissolve. Aliquot in 800 µl batches.

Store in a desiccated environment at -20°C.

Melphalan **Cayman Chemical Company Catalog #**16665

DMSO, Anhydrous **Thermo Fisher Catalog #**D12345

5 **Acryloyl-X Solution (AcX)**

Cannot be prepared in advance.

Stock concentration: 10 mg/ml

Dissolve to 10 mg/ml in anhydrous DMSO. Aliquot in 20 µl batches.

Store in a desiccated environment at -20°C. Don't re-use AcX after thawing.

Working solution will be: 20 µg/ml, dilution 1:500 in 1x PBS

**Reagents**

Acryloyl-X, SE (6-((acryloyl)amino)hexanoic acid, succinimidyl ester) **Thermo Fisher Scientific Catalog #**A20770

DMSO, Anhydrous **Thermo Fisher Catalog #**D12345

6 **Melphalan-X Solution**

Stock concentration: 2 mg/ml

Combine an equal concentration of **Acryloyl-X** (10 mg/ml) and **Melphalan** (2.5 mg/ml) (1-part AcX to 4-parts **Melphalan** (i.e., 200 µl: 800 µl).

Incubate overnight at RT with shaking.

Store in 50 µl aliquots in a desiccated environment at -20°C.

Use at 1 mg/ml by 1:1 dilution in 20 mM **MOPS Buffer**.

Melphalan **Cayman Chemical Company Catalog #**16665

MOPS (Fine White Crystals/Molecular Biology), Fisher BioReagents™ **Thermo Fisher Scientific Catalog #**BP308100

7 **Monomer and Gelation Solutions #1 (Medium Density Gel with high Bis)**

| A | B | C | D | E | F |
|---|---|---|---|---|---|
| | | | | to make 1ml | to make 10ml |
| name | units | stock conc. | final conc. | vol (ul) | vol (ml) |
| Acrylamide | % | 40 | 10 | 250 | 2.5 |
| Na Acrylate | M | 4 | 0.5 | 125 | 1.25 |
| Bis | % | 1 | 0.1 | 100 | 1 |
| 10x PBS | x | 10 | 1 | 100 | 1 |
| Water | | | | 365 | 3.65 |

**Monomer Solution #1**

| A | B | C | D | E |
|---|---|---|---|---|
| | | | | to make 1ml |
| name (units) | units | stock conc. | final conc. | vol (ul) |
| Monomer Solution #1 | | | | 940 |
| APS | % | 10 | 0.2 | 20 |
| TEMED | % | 10 | 0.2 | 20 |
| 4HT | % | 0.5 | 0.01 | 20 |

**Gelation Solution #1**

**Reagents**

- 40% Acrylamide Solution **Bio-Rad Laboratories Catalog #**1610140
- 2% bis-acrylamide solution **Bio-Rad Laboratories Catalog #**1610142
- Acrylic acid **Merck MilliporeSigma (Sigma-Aldrich) Catalog #**147230-5G
- Sodium Hydroxide Solution (10N/Certified) **Fisher Scientific Catalog #**SS255-1
- Ammonium persulfate (APS) **Merck MilliporeSigma (Sigma-Aldrich) Catalog #**A3678-100G
- 4-Hydroxy-TEMPO (4HT) **Merck MilliporeSigma (Sigma-Aldrich) Catalog #**4-Hydroxy-TEMPO
- N,N,N′,N′-Tetramethylethylenediamine (TEMED) **Merck MilliporeSigma (Sigma-Aldrich) Catalog #**T7024-25ML
- UltraPure™ DNase/RNase-Free Distilled Water **Thermo Fisher Scientific Catalog #**10977023

8 **Monomer and Gelation Solutions #2 (Medium Density Gel with low Bis)**

| A | B | C | D | E | F |
|---|---|---|---|---|---|
| | | | | to make 1ml | to make 10ml |
| component | units | stock conc. | final conc. | vol (ul) | vol (ml) |
| Acrylamide | % | 40 | 10 | 250 | 2.5 |
| Na Acrylate | M | 4 | 0.5 | 125 | 1.25 |
| Bis | % | 1 | 0.02 | 20 | 0.2 |
| 10x PBS | x | 10 | 1 | 100 | 1 |
| Water | | | | 445 | 4.45 |

**Monomer Solution #2**

| | A | B | C | D | E |
|---|---|---|---|---|---|
| | | | | | to make 1ml |
| | component | units | stock conc. | final conc. | vol (ul) |
| | Monomer Solution #2 | | | | 940 |
| | TEMED | % | 10 | 0.2 | 20 |
| | 4HT | % | 0.5 | 0.01 | 20 |
| | APS | % | 10 | 0.2 | 20 |

**Gelation Solution #2**

**Reagents**

- 40% Acrylamide Solution **Bio-Rad Laboratories Catalog #**1610140
- 2% bis-acrylamide solution **Bio-Rad Laboratories Catalog #**1610142
- Acrylic acid **Merck MilliporeSigma (Sigma-Aldrich) Catalog #**147230-5G
- Sodium Hydroxide Solution (10N/Certified) **Fisher Scientific Catalog #**SS255-1
- Ammonium persulfate (APS) **Merck MilliporeSigma (Sigma-Aldrich) Catalog #**A3678-100G
- 4-Hydroxy-TEMPO (4HT) **Merck MilliporeSigma (Sigma-Aldrich) Catalog #**4-Hydroxy-TEMPO
- N,N,N′,N′-Tetramethylethylenediamine (TEMED) **Merck MilliporeSigma (Sigma-Aldrich) Catalog #**T7024-25ML
- UltraPure™ DNase/RNase-Free Distilled Water **Thermo Fisher Scientific Catalog #**10977023

9 **Total Protein Stains**

Prepare stock: 10 mg/ml stocks in anhydrous DMSO.

Aliquot in 5-10ul.

Store in a desiccated environment at -20ºC.

**Reagents**

- Alexa Fluor™ 488 NHS Ester (Succinimidyl Ester) **Thermo Fisher Scientific Catalog #**A20000
- ATTO 647N maleimide **AAT Bioquest Catalog #**2857
- DMSO, Anhydrous **Thermo Fisher Catalog #**D12345

10 **SSCT**

0.5x SSC, 0.1% Tween

- SSC (20X), RNase-free **Thermo Scientific Catalog #**AM9763

11 **Disruption Buffer #1:**

500 mM NaCl, 0.3% SDS, 50 mM Tris-HCL pH8 .0, 1 mM EDTA with Proteinase K diluted 1:50 from 800 U/ml stock

**Disruption Buffer #2:**

50 mM NaCl, 1% SDS, 50 mM Tris-HCL pH 8.0, 1 mM EDTA with Proteinase K diluted 1:50 from 800 U/ml stock

- NaCl (5 M), RNase-free **Thermo Fisher Scientific Catalog #**AM9760G
- Proteinase K, Molecular Biology Grade **New England Biolabs Catalog #**P8107S
- EDTA (0.5 M), pH 8.0, RNase-free **Thermo Fisher Scientific Catalog #**AM9260G
- UltraPure™ SDS Solution, 10% **Thermo Fisher Scientific Catalog #**15553027

12 **Gelation Chambers**

Silicone gaskets (Invitrogen P24743)
Glass slides (SuperFrost 12550123)
Scotch tape
Poly-l-lysine
Shaker (nutator - 75rpm)

**Reagents**

- Press-to-Seal™ Silicone Isolator with Adhesive, eight wells, 9 mm diameter, 0.5 mm deep **Thermo Fisher Scientific Catalog #**P24743
- Fisherbrand™ Superfrost™ Disposable Microscope Slides **Fisher Scientific Catalog #**Catalog No.12-550-123
- Poly-L-lysine hydrobromide **Merck MilliporeSigma (Sigma-Aldrich) Catalog #**Poly-L-lysine hydrobromide
- Photo Flo 200 Solution **Electron Microscopy Sciences Catalog #**74257

#### Fixation

2h

13 Euthanize samples with an overdose of MS-222 (a.k.a. tricaine) (200-300 mg/L).

14 Prepare fresh 4% PFA.

15 Place samples in 1 ml of 4% PFA.

Room temperature

16 Keep overnight in 4% PFA at 4°C on a shaker.

1h

Overnight

4 °C

17 Rinse samples in 4 x 15 min 1 ml 1x PBS at 4°C.
This should be done in a cold room in order to minimize RNA degradation.

1h

01:00:00

#### RNA Anchoring

50m

18 Place each fish in separate PCR tube and add 150 µl of 20 mM **MOPS Buffer**.

19 Incubate samples 1x 30mins.

30m

00:30:00

20 Thaw **Melphalan-X** and **Acryloyl-X** solutions during incubation.

21 Dilute **Melphalan-X** 1:1 with **MOPS Buffer**.
Add **Acryloyl-X** 1:100 to **Melphalan-X/MOPS Buffer**.
(100 µl needed per sample)

22 Remove as much of **MOPS Buffer** from the PCR tube as possible.
Add 100 µl **Melphalan-X/MOPS Buffer/Acryloyl-X** solution to each PCR tube.

23 Incubate overnight at 37°C.

37 °C

Overnight

24 Transfer fish to 24-well plate, one per well.
Wash samples 2x 5 min of 1x PBS (600 µl/sample).

10m

00:10:00

#### Permeabilization

1h

25 Permeabilize for 1 h in **PBST-0.5** at RT in 500 ul.

1h

01:00:00

#### Preparation of Gelation Chamber #1

26 Prepare chambers for gelation, one for each sample.
Layer 11 pieces of Scotch tape together to create ~0.6 mm thick spacers.
Cut two strips of spacer material, about 2 cm long and 0.5 cm wide.
Stick two strips to a glass slide ~15 mm apart to form the side walls of the **Gelation Chamber #1**.

#### Incubation with Gelation Solution 1

45m

27 Rinse samples with 1x PBS (3x 5 min).

00:15:00 15m

28 Thaw **Monomer Solution #1**, as well as 4HT, TEMED and APS. Vortex well and keep on ice.

29 Mix **Monomer Solution #1** and 4HT, TEMED and APS at a ratio of 94:2:2:2 to produce **Gelation Solution #1**.
Vortex.
Each sample needs ~1 ml of **Gelation Solution #1**.

30 Remove as much PBS from each sample well.

31 Incubate samples in **Gelation Solution #1** on ice (3x 10 min at 4°C) with 400 µl of gelation solution on a shaker in a 24-well plate. 30m

00:30:00

#### Gelation 1

2h

32 With spatula, transfer fish into the **Gelation Chamber #1**.

33 Gently place cover slip over the fish lying on the walls of **Gelation Chamber #1** (made of scotch tape).

34 Slowly pipette ~200 µl of **Gelation Solution #1** into **Gelation Chamber #1** until full. Avoid any air bubbles.
Solution will hold by water tension.

35 Once **Gelation Chamber #1** is filled, place the chambers at 37°C for 2 h to induce polymerization. 2h
Ensure incubator is humidified.

37 °C

02:00:00

#### Disruption

4h 5m

36 Place gels at RT, and allow them to cool on the bench for > 5 min. 5m

Room temperature

00:05:00

37 Take off coverslip lid with a razor blade.

Under a stereomicroscope, trim the gels into a rectangle with a ~2-3 mm border on either side of the sample.
Add a nick on the top right corner to be able to track orientation of the samples.

38 Transfer each gel to a 2 ml Eppendorf tube and add 750 µl of **Disruption Buffer #1**.
**Disruption Buffer #1:**
500 mM NaCl, 0.3% SDS, 50 mM Tris-HCL pH8 .0, 1 mM EDTA with Proteinase K diluted 1:50 from 800 U/mL stock

39 Incubate samples (50ºC for 4 hr). 4h

50 °C

04:00:00

40 Remove liquid from Eppendorf tube.
Add 1ml of **Disruption Buffer #2** to Eppendorf tube**.**
**Disruption Buffer #2:**
50 mM NaCl, 1% SDS, 50 mM Tris-HCL pH 8.0, 1 mM EDTA with Proteinase K diluted 1:50 from 800 U/mL stock

41 Incubate each samples at 50ºC overnight.

50 °C

Overnight

#### Chamber 2 preparation

42 Prepare **Gelation Chamber #2**, one for each sample.
Layer 20 pieces of Scotch tape together to create ~1.2 mm thick spacers.
Cut two strips of spacer material, about 2 cm long and 0.5 cm wide.
Stick two strips to a glass slide ~15 mm apart to form the side walls of the **Gelation Chamber #2**.

#### Incubation with Gelation Solution 2 1h 30m

43 Wash each samples in 1x PBS at RT (3x 20 min). 1h

Room temperature

01:00:00

44 Thaw **Monomer Solution #2,** as well as 4HT, TEMED and APS. Vortex well and keep on ice.

45 Mix **Monomer Solution #2** and 4HT, TEMED and APS at a ratio of 94:2:2:2 to produce **Gelation Solution #2**.
Vortex.
Each sample needs ~6 ml of **Gelation Solution #2**.
~2 ml per incubation round.

46 Incubate samples in **Gelation Solution #2** on ice (3x 10 mins @ 4°C) with 2ml of **Gelation Solution #2** on a shaker in 12-well plate.

4 °C

00:30:00

30m

#### Gelation 2

2h 5m

47 With spatula, transfer gel block containing fish into a **Gelation Chamber #2**.
Make sure there are no bubbles.
Add small drop of gelation chamber.
Carefully place coverslip on top (Corning #2855-25)

**Reagents**

Corning® 25x25 mm Square #2 Cover Glass **Corning Catalog #**2855-25

48 Slowly pipette **Gelation Solution #2** into **Gelation Chamber #2** until full. Avoid any air bubbles.
Solution will hold by water tension.
Cover.

49 Once **Gelation Chamber #2** is filled, place the chambers at 37°C for 2 hr to induce polymerization.
Ensure incubator is humidified.

37 °C

02:00:00

2h

50 Place gels at RT, and allow them to cool on the bench for > 5 min.

Room temperature

00:05:00

5m

51 Take off coverslip lid with a razor blade.
Under a stereomicroscope, remove the scotch side walls and trim the gels into a rectangle, close to the first gel.
Removing any second gel material that formed outside of the first gel.

52 Transfer gel to a 12-well plate with 1x PBS.

#### Second Disruption

4h

53 Transfer each gel to a 2 ml Eppendorf tube and add 750 µl of **Disruption Buffer #1**.
**Disruption Buffer #1:**
500 mM NaCl, 0.3% SDS, 50 mM Tris-HCL pH8 .0, 1 mM EDTA with Proteinase K diluted 1:50 from 800 U/mL stock

54 Incubate samples (50°C for 4 hr).

4h

50 °C

04:00:00

55 Remove liquid from Eppendorf tube.
Add 1ml of **Disruption Buffer #2** to Eppendorf tube**.**
**Disruption Buffer #2:**
50 mM NaCl, 1% SDS, 50 mM Tris-HCL pH 8.0, 1 mM EDTA with Proteinase K diluted 1:50 from 800 U/mL stock

56 Incubate each samples at 50°C overnight.

50 °C

Overnight

#### Hybridization with Probes

30m

57 Wash samples 4x 15 min with 1 ml 1x PBS.

58 Thaw and mix **Hybridization Buffer** and **Probe Wash Buffer**.

59 Incubate each gel in 600 µl **Hybridization Buffer** for 30 min at 37°C. 30m

37 °C

00:30:00

60 Dilute **HCR probes** 6 µl in 600 µl **Hybridization Buffer** per gel (10 nM final concentration).
Vortex.

61 Incubate gel with **HCR probes** overnight at 37°C. No shaking is necessary.

37 °C

Overnight

62 Place **Probe Wash Buffer** and 1x PBS at 37°C in preparation for washing the next day.

#### Probe Wash

1h 30m

63 Wash gels in pre-warmed (37°C) **Probe Wash Buffer** for 3x 30min. 1h 30m
Keep in 37°C incubator.

01:30:00

64 Wash gels in pre-warmed (37°C) 1x PBS for 3x 30 min, then 1x PBS for 3x 60 min.
Keep in 37°C incubator.

Wash gels in 1x PBS overnight at RT.

Overnight

#### Hybridization Chain Reaction

5h 20m

65 Incubate each gel in 1 ml **Amplification Buffer** for at least 30 min at RT.

66 Snap cool hairpins with PCR machine at 95ºC for 90 sec and cool to 25ºC for 30 min.

67 For each fluor mix, add hairpins h1 and h2 at 1:50 in 400 µl **Amplification Buffer** (1x per gel). Vortex.

68 Incubate each gel with hairpins for 4 hr at RT in the dark. 4h

04:00:00

69 Wash gels 2 x 20 min in 1 ml 5x SSCT at RT. 40m

00:40:00

SSCT: 5x SSC, 0.1% Tween

SSC (20X), RNase-free **Thermo Scientific Catalog #**AM9763

70 Wash gels 2 x 40 min in 1 ml 0.5x SSCT at RT. 40m

00:40:00

71 Place in 1x PBS and leave to equilibrate at least 1 h before imaging.

#### Mounting

72 Attach a 8mm circular coverslips (CS-8R, Warner Instruments, 64-0701) to a Z1 sample holder using superglue.

**Reagents**

CS-8R Coverslips, 0.15 mm (0.006 in), 8 mm diameter, pkg of 100 **Multi Channel Systems MCS GmbH Catalog #**640701

Z1 Sample Holder **Janelia Research Campus**

LIGHTSHEETHOLDER V13 Short.ipt

LIGHTSHEETHOLDER V13 Short.stl

73 Coat coverslip with poly-lysine and let it dry.

Poly-L-lysine hydrobromide **Merck MilliporeSigma (Sigma-Aldrich) Catalog #**Poly-L-lysine hydrobromide

Photo Flo 200 Solution **Electron Microscopy Sciences Catalog #**74257

74 Mount gel, sample side up.

#### Imaging

75 Image using a Zeiss Z1 Light Sheet Microscope. Allow 45 min (with gel in place) for system to equilibrate before acquiring multi-tile acquisitions.
Image with 20x 1.0 NA water immersion objective.
Exposure time: 100-150 ms.

76 After imaging, gels can be teased off holder with a paint brush.

#### DNAse treatment for Stripping Hairpins

77 Incubate gel for 30 min in 1 ml of **DNase1 Buffer** at 37°C

RNase-Free DNase Set **Qiagen Catalog #**79254

78 Add 450 µl of **DNase Buffer** to 50 ml DNase1. Mix. (increase to 750 µl if gel is not fully covered).

RNase-Free DNase Set **Qiagen Catalog #**79254

79 Incubate gel in DNase1 for 2 hr at 37°C.

RNase-Free DNase Set **Qiagen Catalog #**79254

80 Wash 4 x 15 min with 1 ml 1x PBS.

#### Protocol references

1. Kunst, M. *et al.* A Cellular-Resolution Atlas of the Larval Zebrafish Brain. *Neuron* 103, 21-38.e5 (2019).
2. Randlett, O. *et al.* Whole-brain activity mapping onto a zebrafish brain atlas. *Nat Methods* 12, 1039–1046 (2015).
3. Tabor, K. M. *et al.* Brain-wide cellular resolution imaging of Cre transgenic zebrafish lines for functional circuit-mapping.
4. Copper, J. E. *et al.* Comparative analysis of fixation and embedding techniques for optimized histological preparation of zebrafish. *Comparative Biochemistry and Physiology Part C: Toxicology & Pharmacology* 208, 38–46 (2018).
5. Ding, Y. *et al.* Computational 3D histological phenotyping of whole zebrafish by X-ray histotomography. *eLife* 8, e44898 (2019).
6. Steib, E. *et al.* TissUExM enables quantitative ultrastructural analysis in whole vertebrate embryos by expansion microscopy. *Cell Reports Methods* 2, 100311 (2022).
7. Sim, J. *et al.* Nanoscale resolution imaging of the whole mouse embryos and larval zebrafish using expansion microscopy. Preprint at https://doi.org/10.1101/2021.05.18.443629 (2021).
8. Chen, F., Tillberg, P. W. & Boyden, E. S. Expansion microscopy.
9. Wang, Y. *et al.* EASI-FISH for thick tissue defines lateral hypothalamus spatio-molecular organization. *Cell* 184, 6361-6377.e24 (2021).

#### Acknowledgements

This research was funded by Janelia Research Campus, HHMI and the Gatsby Computational Neuroscience Unit.

### Whole Body Expansion Microscopy for Danionella and older zebrafish

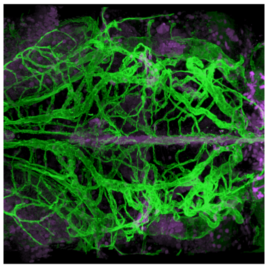

Virginia Ruetten[1], Yisheng He[2], Mark Eddison[3], Kari Close[3], Amy Hu[4], Misha B. Ahrens[3], Paul Tillberg[3]

[1]Janelia Research Campus, HHMI - Gatsby Computational Neuroscience Unit; [2]HHMI; [3]HHMI/Janelia Research Campus; [4]Janelia Research Campus

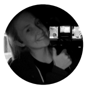

Virginia Ruetten
Janelia Research Campus, HHMI - Gatsby Computational Neurosc...

**Protocol Info:** Virginia Ruetten, Yisheng He, Mark Eddison, Kari Close, Amy Hu, Misha B. Ahrens, Paul Tillberg . Whole Body Expansion Microscopy for Danionella and older zebrafish. **protocols.io** **https://protocols.io/view/whole-body-expansion-microscopy-for-danionella-and-dxqr7mv6**

**Created:** January 13, 2025

**Last Modified:** March 10, 2025

**Protocol Integer ID:** 118257

**Keywords:** Expansion Microscopy, Zebrafish, immunohistochemistry, clearing

#### Abstract

The interpretation of Whole Body Imaging (WBI) data necessitates comprehensive anatomical knowledge to accurately determine cell-type identity; however, resources pertaining to the internal anatomy of Danionella species are limited. To mitigate this gap, we established an advanced Whole Body Expansion-Microscopy (WB-ExM) protocol, facilitating the acquisition of molecular and cell-type information at subcellular resolution throughout the entire organism. This method involves the use of high-temperature (100°C) chemical hydrolysis to uniformly soften tissues, embedding within a medium-density gel with reduced protein-gel anchoring, and repeated embedding post-digestion with moderate ~1.5x expansion factors in each cycle, culminating in a robust and uniform expansion even of challenging structures such as cartilage embedded in soft tissue. This protocol results in excellent optical clearing and retains high levels of antibody signals.

### Image Attribution

Virginia M. S. Ruetten

#### Protocol materials

- Z1 Sample Holder **Janelia Research Campus** Step 60
- PBS, pH 7.4 **Thermo Fisher Catalog #**10010001 Step 1
- DMSO, Anhydrous **Thermo Fisher Catalog #**D12345 In 2 steps
- Sodium Hydroxide Solution (10N/Certified) **Fisher Scientific Catalog #**SS255-1 Step 6
- Corning® 25x25 mm Square #2 Cover Glass **Corning Catalog #**2855-25 Step 52
- ATTO 647N maleimide **AAT Bioquest Catalog #**2857 Step 58
- Pierce™ 16% Formaldehyde (w/v), Methanol-free **Thermo Scientific Catalog #**28906 Step 0.1
- 4-Hydroxy-TEMPO (4HT) **Merck MilliporeSigma (Sigma-Aldrich) Catalog #**4-Hydroxy-TEMPO Step 5
- 2% bis-acrylamide solution **Bio-Rad Laboratories Catalog #**1610142 Step 5
- 4-Hydroxy-TEMPO (4HT) **Merck MilliporeSigma (Sigma-Aldrich) Catalog #**4-Hydroxy-TEMPO Step 6
- Acrylic acid **Merck MilliporeSigma (Sigma-Aldrich) Catalog #**147230-5G Step 6
- UltraPure™ SDS Solution, 10% **Thermo Fisher Scientific Catalog #**15553027 Step 7
- Hydrogen peroxide solution **Merck MilliporeSigma (Sigma-Aldrich) Catalog #**H1009-500ML Step 15
- Triton™ X-100 **Merck MilliporeSigma (Sigma-Aldrich) Catalog #**X100-5ML Step 1
- Hydrogen peroxide solution **Merck MilliporeSigma (Sigma-Aldrich) Catalog #**H1009-500ML Step 2
- Agarose, low gelling temperature **Merck MilliporeSigma (Sigma-Aldrich) Catalog #**A9414-100G Step 3
- PBS - Phosphate-Buffered Saline (10X) pH 7.4, RNase-free **Thermo Fisher Scientific Catalog #**AM9625 Step 3
- Acryloyl-X, SE (6-((acryloyl)amino)hexanoic acid, succinimidyl ester) **Thermo Fisher Scientific Catalog #**A20770 Step 4
- Poly-L-lysine hydrobromide **Merck MilliporeSigma (Sigma-Aldrich) Catalog #**Poly-L-lysine hydrobromide Step 9
- 40% Acrylamide Solution **Bio-Rad Laboratories Catalog #**1610140 Step 6
- Photo Flo 200 Solution **Electron Microscopy Sciences Catalog #**74257 Step 9
- Ammonium persulfate (APS) **Merck MilliporeSigma (Sigma-Aldrich) Catalog #**A3678-100G Step 5
- PBS - Phosphate-Buffered Saline (10X) pH 7.4, RNase-free **Thermo Fisher Scientific Catalog #**AM9625 Step 0.1
- UltraPure™ DNase/RNase-Free Distilled Water **Thermo Fisher Scientific Catalog #**10977023 In 4 steps
- ATTO 647N maleimide **AAT Bioquest Catalog #**2857 Step 8
- Press-to-Seal™ Silicone Isolator with Adhesive, one well, 20 mm diameter, 0.5 mm deep **Invitrogen - Thermo Fisher Catalog #**P24740 In 2 steps
- Sodium Hydroxide Solution (10N/Certified) **Fisher Scientific Catalog #**SS255-1 Step 5
- Acrylic acid **Merck MilliporeSigma (Sigma-Aldrich) Catalog #**147230-5G Step 5

N,N,N′,N′-Tetramethylethylenediamine (TEMED) **Merck MilliporeSigma (Sigma-Aldrich) Catalog #**T7024-25ML

Step 6

2% bis-acrylamide solution **Bio-Rad Laboratories Catalog #**1610142 Step 6

CS-8R Coverslips, 0.15 mm (0.006 in), 8 mm diameter, pkg of 100 **Multi Channel Systems MCS GmbH Catalog #**640701

Step 60

NaCl (5 M), RNase-free **Thermo Fisher Scientific Catalog #**AM9760G Step 7

Alexa Fluor™ 488 NHS Ester (Succinimidyl Ester) **Thermo Fisher Scientific Catalog #**A20000 Step 8

Fisherbrand™ Superfrost™ Disposable Microscope Slides **Fisher Scientific Catalog #**Catalog No.12-550-123

In 2 steps

Alexa Fluor™ 488 NHS Ester (Succinimidyl Ester) **Thermo Fisher Scientific Catalog #**A20000 Step 58

40% Acrylamide Solution **Bio-Rad Laboratories Catalog #**1610140 Step 5

N,N,N′,N′-Tetramethylethylenediamine (TEMED) **Merck MilliporeSigma (Sigma-Aldrich) Catalog #**T7024-25ML

Step 5

Ammonium persulfate (APS) **Merck MilliporeSigma (Sigma-Aldrich) Catalog #**A3678-100G Step 6

#### Before start

Make sure you have all the reagents at hand.

#### Reagents

1 **4% PFA**

Cannot be prepared in advance.

Take a new ampule of 16% PFA (10ml). Aliquot it into 1 ml aliquots. Store at -80ºC. On the day take one aliquot out and thaw.

To make a total volume of 4 ml of 4% PFA: mix 1ml of 16% PFA, 0.4 ml of 10x PBS and 2.6 ml of distilled H20. Use within the next two days and store in a 4ºC fridge.

**Note**

PFA deteriorates even if stored at -80ºC. Avoid reusing aliquoted samples.
When PFA is stored in sealed ampules, it is protected from atmospheric oxygen and moisture. This isolation prevents oxidative degradation

**Reagents**

- PBS - Phosphate-Buffered Saline (10X) pH 7.4, RNase-free **Thermo Fisher Scientific Catalog #**AM9625
- Pierce™ 16% Formaldehyde (w/v), Methanol-free **Thermo Scientific Catalog #**28906
- UltraPure™ DNase/RNase-Free Distilled Water **Thermo Fisher Scientific Catalog #**10977023

1 **PBST-0.5**

Can be prepared in advance.

1x PBS with 0.5% or 0.1% Triton

PBST-0.5 and PBST-0.1 respectively

Triton dissolves terribly. It hardens upon making contact with water and takes time to dissolve fully. Prepare a 10% stock solution in 1x PBS and use that for subsequent rounds.

Keep at RT.

Room temperature

**Reagents**

- PBS, pH 7.4 **Thermo Fisher Catalog #**10010001
- Triton™ X-100 **Merck MilliporeSigma (Sigma-Aldrich) Catalog #**X100-5ML

2 **Bleaching Solution**

Cannot be prepared in advance.

Combine 3% $H_2O_2$, 1% KOH (by weight) and 96% $H_2O$.

- Hydrogen peroxide solution **Merck MilliporeSigma (Sigma-Aldrich) Catalog #**H1009-500ML

3 **1% low-melting temperature agarose**

Can be prepared in advance.

Dissolve 1 g of low-melting point agarose in 100 ml of 1x PBS.

Add 1 g of power to 100 ml 1x PBS at RT. Stir with magnetic stirrer.
Bring to a boil using a microwave. Stir with magnetic stirrer until full dissolved.
Repeat boil and stirring until solution is clear.

**Reagents**

Agarose, low gelling temperature **Merck MilliporeSigma (Sigma-Aldrich) Catalog #**A9414-100G

PBS - Phosphate-Buffered Saline (10X) pH 7.4, RNase-free **Thermo Fisher Scientific Catalog #**AM9625

4 **Acryloyl-X Solution (AcX)**

Cannot be prepared in advance.
Stock concentration: 10 mg/ml
Dissolve to 10 mg/ml in anhydrous DMSO. Aliquot in 20 µl batches.
Store in a desiccated environment at -20°C. Don't re-use AcX after thawing.
Working solution will be: 20 µg/ml, dilution 1:500 in 1x PBS

**Reagents**

Acryloyl-X, SE (6-((acryloyl)amino)hexanoic acid, succinimidyl ester) **Thermo Fisher Scientific Catalog #**A20770

DMSO, Anhydrous **Thermo Fisher Catalog #**D12345

5 **Monomer and Gelation Solutions #1 (Medium Density Gel with high Bis)**

| A | B | C | D | E | F |
|---|---|---|---|---|---|
| | | | | to make 1ml | to make 10ml |
| component | units | stock conc. | final conc. | vol (ul) | vol (ml) |
| Acrylamide | % | 40 | 10 | 250 | 2.5 |
| Na Acrylate | M | 4 | 0.5 | 125 | 1.25 |
| Bis | % | 1 | 0.1 | 100 | 1 |
| 10x PBS | x | 10 | 1 | 100 | 1 |
| Water | | | | 365 | 3.65 |

**Monomer Solution #1**

| A | B | C | D | E |
|---|---|---|---|---|
| | | | | to make 1ml |
| component | units | stock conc. | final conc. | vol (ul) |
| Monomer Solution #1 | x | 1 | 1 | 990 |
| TEMED | % | 10 | 0.1 | 10 |

**Gelation Solution #1 with TEMED only**

| A | B | C | D | E |
|---|---|---|---|---|
| | | | | to make 1ml |
| component | units | stock conc. | final conc. | vol (ul) |
| Monomer Solution #1 | x | 1 | 1 | 980 |
| TEMED | % | 10 | 0.1 | 10 |
| APS | % | 10 | 0.1 | 10 |

**Gelation Solution #1 with TEMED and APS**

**Reagents**

- 40% Acrylamide Solution **Bio-Rad Laboratories Catalog #**1610140
- 2% bis-acrylamide solution **Bio-Rad Laboratories Catalog #**1610142
- Acrylic acid **Merck MilliporeSigma (Sigma-Aldrich) Catalog #**147230-5G
- Sodium Hydroxide Solution (10N/Certified) **Fisher Scientific Catalog #**SS255-1
- UltraPure™ DNase/RNase-Free Distilled Water **Thermo Fisher Scientific Catalog #**10977023
- Ammonium persulfate (APS) **Merck MilliporeSigma (Sigma-Aldrich) Catalog #**A3678-100G
- 4-Hydroxy-TEMPO (4HT) **Merck MilliporeSigma (Sigma-Aldrich) Catalog #**4-Hydroxy-TEMPO
- N,N,N′,N′-Tetramethylethylenediamine (TEMED) **Merck MilliporeSigma (Sigma-Aldrich) Catalog #**T7024-25ML

6 **Monomer and Gelation Solutions #2 (Medium Density Gel with low Bis)**

| A | B | C | D | F |
|---|---|---|---|---|
| | | | | to make 10ml |
| component | units | stock conc. | final conc. | vol (ml) |
| Acrylamide | % | 40 | 10 | 2.5 |
| Na Acrylate | M | 4 | 0.5 | 1.25 |
| Bis | % | 1 | 0.02 | 0.2 |
| 10x PBS | x | 10 | 1 | 0.9 |
| Gel (1x PBS) | x | | | ~1 |
| Water | | | | 4.15 |

**Monomer Solution #2**

| A | B | C | D | E |
|---|---|---|---|---|
| | | | | to make 1ml |
| component | units | stock conc. | final conc. | vol (ul) |
| Monomer Solution #2 | | | | 990 |
| TEMED | % | 10 | 0.1 | 10 |

**Gelation Solution #2 with TEMED only**

| A | B | C | D | E |
|---|---|---|---|---|
| | | | | to make 1ml |
| component | units | stock conc. | final conc. | vol (ul) |
| Monomer Solution #2 | | | | 980 |
| APS | % | 10 | 0.1 | 10 |
| TEMED | % | 10 | 0.1 | 10 |

**Gelation Solution #2 with TEMED and APS**

**Reagents**

- 40% Acrylamide Solution **Bio-Rad Laboratories Catalog #**1610140
- 2% bis-acrylamide solution **Bio-Rad Laboratories Catalog #**1610142
- Acrylic acid **Merck MilliporeSigma (Sigma-Aldrich) Catalog #**147230-5G
- Sodium Hydroxide Solution (10N/Certified) **Fisher Scientific Catalog #**SS255-1
- UltraPure™ DNase/RNase-Free Distilled Water **Thermo Fisher Scientific Catalog #**10977023
- Ammonium persulfate (APS) **Merck MilliporeSigma (Sigma-Aldrich) Catalog #**A3678-100G
- 4-Hydroxy-TEMPO (4HT) **Merck MilliporeSigma (Sigma-Aldrich) Catalog #**4-Hydroxy-TEMPO
- N,N,N′,N′-Tetramethylethylenediamine (TEMED) **Merck MilliporeSigma (Sigma-Aldrich) Catalog #**T7024-25ML

7

| A | B | C | D | E | F |
|---|---|---|---|---|---|
| | | | | to make 1ml | to make 100ml |
| component | units | stock conc. | final conc. | vol (ul) | vol |
| SDS | % | 10 | 5 | 0.5 | 50 |
| Tris pH 7.5 | mM | 1000 | 50 | 0.05 | 5 |
| NaCl | M | 5 | 0.2 | 0.04 | 4 |
| MilliQ water | | | | 0.41 | 41 |

**Disruption Buffer**

Stock **Disruption Buffer** can be prepared in advanced and stored at RT.

**Reagents**

- NaCl (5 M), RNase-free **Thermo Fisher Scientific Catalog #**AM9760G
- UltraPure™ SDS Solution, 10% **Thermo Fisher Scientific Catalog #**15553027
- UltraPure™ DNase/RNase-Free Distilled Water **Thermo Fisher Scientific Catalog #**10977023

8 **Total Protein Stains**

Prepare stock: 10 mg/ml stocks in anhydrous DMSO.
Aliquot in 5-10ul.
Store in a desiccated environment at -20°C.

**Reagents**

- Alexa Fluor™ 488 NHS Ester (Succinimidyl Ester) **Thermo Fisher Scientific Catalog #**A20000
- ATTO 647N maleimide **AAT Bioquest Catalog #**2857
- DMSO, Anhydrous **Thermo Fisher Catalog #**D12345

9 **Gelation Chambers**

Silicone gaskets 20mm (Invitrogen P24740)
Glass slides (SuperFrost 12550123)
Scotch tape
Poly-l-lysine
Shaker (nutator - 75 RPM)

**Reagents**

- Press-to-Seal™ Silicone Isolator with Adhesive, one well, 20 mm diameter, 0.5 mm deep **Invitrogen - Thermo Fisher Catalog #**P24740
- Fisherbrand™ Superfrost™ Disposable Microscope Slides **Fisher Scientific Catalog #**Catalog No.12-550-123
- Poly-L-lysine hydrobromide **Merck MilliporeSigma (Sigma-Aldrich) Catalog #**Poly-L-lysine hydrobromide

Photo Flo 200 Solution **Electron Microscopy Sciences Catalog #**74257

#### Fixation and Permeabilization

10 Euthanize samples with an overdose of MS-222 (a.k.a. tricaine) (200-300 mg/L).

11 Prepare fresh 4% PFA.

12 Place samples in 1 ml of 4% PFA.

Room temperature

13 Keep overnight in 4% PFA at 4°C on a shaker.

Overnight

4 °C

14 Rinse samples in 4 x 15 min 1 ml 1x PBS.

Room temperature

#### Bleaching

15 Prepare fresh **Bleaching Solution**

Combine 3% $H_2O_2$, 1% KOH (by weight) and 96% $H_2O$.

Room temperature

Hydrogen peroxide solution **Merck MilliporeSigma (Sigma-Aldrich) Catalog #**H1009-500ML

16 Incubate samples in 500 µL of **Bleaching Solution** until bubbling is observed. Replace the bleaching solution with fresh bleaching solution. Monitor the samples until pigmentation disappears (~2 min).

Room temperature

17 Rinse the samples twice with 1x PBS.

Room temperature

#### Permeabilization (if no IHC)

5h 10m

18 Permeabilize samples (at least 5 hr or overnight) in **PBST-0.5**.

5h

05:00:00

19 Wash samples (2x 5 min at RT) in 1x PBS. 10m

Room temperature

00:10:00

#### Agarose Embedding

20 Heat agarose in microwave until it liquifies.
Cool agarose to 50°C (by placing in 50°C incubator).

21 Layer 2 to 3 layers of silicone gasket (Invitrogen P24743 or Invitrogen P24740) on top of each other. (For ~1.5 months old 2 layer typically suffices, for older 3). Adhere silicone gaskets on a glass slide (Superfrost Microscope Slides #12550123) to form a **Mounting Chamber**.

**Reagents**

Press-to-Seal™ Silicone Isolator with Adhesive, one well, 20 mm diameter, 0.5 mm deep **Invitrogen - Thermo Fisher Catalog #**P24740

Fisherbrand™ Superfrost™ Disposable Microscope Slides **Fisher Scientific Catalog #**Catalog No.12-550-123

22 Place fish in **Mounting Chamber**, and cover with 400 µl of 1% agarose. Orientate as desired (on side or dorsal side up) and leave to solidify.

23 With a razor blade, cut out a rectangle around the sample and delicately lift and transfer to 12 well-plate with 1ml of PBS/well. Cut as close to the fish as possible with some safety margin.

#### Protein Anchoring 1h

24 Prepare **Acryloyl-X Solution** (stock solution 10 mg/ml, working solution: 20 µg/ml, dilution 1:500 in 1x PBS).
1 ml per sample is needed.
Do not prepare in advance.

25 Incubate each sample in 1 ml of **Acryloyl-X Solution** (1 hr at RT) shaking in 12-well plate. 1h

Room temperature

01:00:00

#### Preparation of Gelation Chamber #1

26 Prepare chambers for gelation, one for each sample.
Layer 20 pieces of Scotch tape together to create ~1.2 mm thick spacers.
Cut two strips of spacer material, about 2.5 cm long and ~0.20 cm wide.
Stick two strips to a glass slide ~21 mm apart to form the side walls of the **Gelation Chamber #1**.

Before placing the gel in the chamber, add a small amount of vacuum grease on the side walls to ensure a tight seal upon placing the glass cover slip on top of the chamber.

#### First Incubation with Gelation Solution 1 (with TEMED only)

15m

27 Rinse samples with 1x PBS (3x 5 min).

00:15:00

15m

28 Thaw **Monomer Solution #1**, as well as TEMED. (Note: no APS required). Vortex well and keep on ice.

29 Mix **Monomer Solution #1** and TEMED at a ratio of 99:1 to produce **Gelation Solution #1**.
Vortex.
Each sample needs ~2 ml of **Gelation Solution #1**..
~2 ml per incubation round.

30 Incubate samples in **Gelation Solution #1** overnight in an Eppendorf tube with 2 ml of **Gelation Solution #1** on a rotator (4°C).

#### Second incubation with Gelation Solution 1 (with TEMED and APS only)

15m

31 Thaw **Monomer Solution #1**, as well as APS and TEMED. Vortex well and keep on ice.

32 Mix **Monomer Solution #1** and APS and TEMED at a ratio of 98:1:1 to produce **Gelation Solution #1**.
Vortex.
Each sample needs ~5 ml of **Gelation Solution #1**..
~5 ml per incubation round.

33 Incubate samples in **Gelation Solution #1** in a 5ml Eppendorf tube with 5 ml of **Gelation Solution #1** on a rotator (4h at 4°C).

#### Gelation 1

2h

34 With spatula, transfer agarose block containing fish into the **Gelation Chamber #1**.

35 Gently place cover slip over the agarose block lying on the walls of **Gelation Chamber #1** (made of scotch tape).
The cover slip should lay flat on the agarose block.
Gently press to seal.

36 Slowly pipette **Gelation Solution #1** into **Gelation Chamber #1** until full. Avoid any air bubbles.
Solution will hold by water tension.

One can use **Gelation Solution #1** that was used during the previous incubation (no need to prepare fresh one).

37 Once **Gelation Chamber #1** is filled, place the chambers at 37°C for 2 hr to induce polymerization. 2h
Ensure incubator is humidified.

37 °C

02:00:00

#### Disruption 9h 5m

38 Turn on the 95°C heat bath (for later disruption step).
Prepare **Disruption Buffer** (5 ml/sample).
Stock **Disruption Buffer** can be prepared in advanced and stored at RT.

39 Place gels at RT, and allow them to cool on the bench for > 5 min. 5m

Room temperature

00:05:00

40 Take off coverslip lid with a razor blade.
Under a stereomicroscope, trim the gels into a rectangle with a ~2-3 mm border on either side of the sample.
Add a nick on the top right corner to be able to track orientation of the samples.
The gel should be about 9 mm x 4 mm.

41 Transfer each gel to a 5 ml Eppendorf tube and add 5 ml of **Disruption Buffer.**

42 Incubate each samples in **Disruption Buffer** (overnight at 95°C). 9h
Make sure that lid is closed tightly to avoid evaporation.

95 °C

09:00:00

#### Chamber 2 preparation

43 Prepare **Gelation Chamber #2**, one for each sample.
Layer 40 pieces of Scotch tape together to create ~2.4 mm thick spacers.
Cut two strips of spacer material, about 2.5 cm long and ~0.20 cm wide.
Stick two strips to a glass slide ~21 mm apart to form the side walls of the **Gelation Chamber #2**.
Before placing the gel in the chamber, add a small amount of vacuum grease on the side walls to ensure a tight seal upon placing the glass cover slip on top of the chamber.

#### First incubation with Gelation Solution 2 (with TEMED only) 1h

44 Wash each samples in 1x PBS at RT (3x 20 min).

1h

Room temperature

01:00:00

45 Thaw **Monomer Solution #2,** as well as TEMED. Vortex well and keep on ice.

46 Trim the gels to remove excess material.
Weigh the gels (~400 to 500mg), and transfer each to 6-well plate.
Add 1x PBS to make total weigh 500 mg (i.e.: 500 µl).
As the gel is so large, this represents a substantial amount of PBS. Therefore, the weight of the gel (which is mostly PBS) should be discounted from the PBS added when making **Gelation Solution #2**. That is to say, assume the gel is 1x PBS.

47 Mix **Monomer Solution #2** and TEMED at a ratio of 99:1 to produce **Gelation Solution #2**.
Vortex.
Each sample needs ~5 ml of **Gelation Solution #1**..
~5 ml per incubation round.

48 Incubate samples in **Pre-incubation Gelation Solution #2** overnight in an 5ml Eppendorf tube on a rotator (4°C).

#### Second Incubation with Gelation Solution 2 (with TEMED and APS)

49 Thaw **Monomer Solution #2**, as well as APS and TEMED.
Vortex well and keep on ice.

50 Mix **Monomer Solution #2** and APS and TEMED at a ratio of 98:1:1 to produce **Gelation Solution #2**.
Vortex.
Each sample needs ~5 ml of **Gelation Solution #2**..
~5 ml per incubation round.

51 Incubate samples in **Gelation Solution #2** in a 5ml Eppendorf tube fill up to 4.5ml of **Gelation Solution #2** on a rotator (4h at 4°C). It is important to leave some air gap (to prevent polymerization).

#### Gelation 2

5m

52 With spatula, transfer gel block containing fish into a **Gelation Chamber #2**.
Make sure there are no bubbles at the interface between chamber and gel.
If needed, add a small drop of gelation solution below the gel if air bubbles remain.
Add a small drop of gelation solution on top of the gel to minimize chances of air bubbles forming.
Carefully place coverslip on top (Corning #2855-25).

Do not add extra gelation solution!
Wipe off the extra gelation solution with a Kimwipe.

**Reagents**

Corning® 25x25 mm Square #2 Cover Glass **Corning Catalog #**2855-25

53 Once **Gelation Chamber #2** is ready, place a piece of wet paper towel in a plastic box.
Next place the prepared chamber on the wet paper towel.
Finally place the plastic box into a plastic bag.

54 Fill the plastic bag with nitrogen. Seal the bag.
Place the chambers at 37°C for 2 hr to induce polymerization.

55 Place gels at RT, and allow them to cool on the bench for > 5 min. 5m

Room temperature

00:05:00

56 Take off coverslip lid with a razor blade.
Under a stereomicroscope, remove the scotch side walls.

57 Turn on the 95°C heat pebble bath (for later disruption step).
Prepare **Disruption Buffer** (5 ml/sample).
Transfer each gel to a 5 ml Eppendorf tube and fill with **Disruption Buffer.**
Incubate overnight at 95°C.
Make sure that lid is closed tightly to avoid evaporation.

95 °C

09:00:00

#### Staining

1h 30m

58 Incubate gels in Alexa488-NHS and Atto647N-maleimide dye at 1:1000 each in 1x PBS (overnight at 4°C) in 5ml Eppendorf tube on gentle shaker.

**Reagents**

Alexa Fluor™ 488 NHS Ester (Succinimidyl Ester) **Thermo Fisher Scientific Catalog #**A20000

ATTO 647N maleimide **AAT Bioquest Catalog #**2857

59 Wash in 1x PBS (3x 3 hr) on shaker at RT. 1h 30m

Room temperature

01:30:00

#### Mounting

60 Attach a 8mm circular coverslips (CS-8R, Warner Instruments, 64-0701) to a Z1 sample holder using superglue.

**Reagents**

CS-8R Coverslips, 0.15 mm (0.006 in), 8 mm diameter, pkg of 100 **Multi Channel Systems MCS GmbH Catalog #**640701

Z1 Sample Holder **Janelia Research Campus**

LIGHTSHEETHOLDER V13 Short.ipt LIGHTSHEETHOLDER V13 Short.stl

61 Coat coverslip with poly-lysine and let it dry.

62 Mount gel, sample side up.

#### Imaging

63 Image using a Zeiss Z1 Light sheet Microscope. Allow 45 min (with gel in place) for system to equilibrate before acquiring multi-tile acquisitions.
Image with 20x 1.0 NA water immersion objective.
Exposure time: 100-150 ms.

64 After imaging, gels can be teased off holder with a paint brush.

65 For long term storage, keep gels in 1x PBS at 4°C.

#### Acknowledgements

All samples were supplied by Chie Satou.

#### Table of transgenic lines generated and used

| Transgenic Line | Target Cell Population | Protein Function | Source | Species | References |
| --- | --- | --- | --- | --- | --- |
| Tg(ubi:tTA; TRE:jRGECO1b) | Pancellular | Red calcium indicator | generated | Danio rerio | Mosimann et al. (2011) \| Dana et al. (2016) |
| Tg(ubi:tTA); Tg(TRE:GCaMP7f) | Pancellular | Green calcium indicator | generated | Danio rerio | Mosimann et al. (2011) \| Dana et al. (2019) |
| Tg(ubbR:jGCaMP8m) | Pancellular | Green calcium indicator | generated | Danio rerio | Bakunaite et al. (2024) \| Zhang et al. (2023) |
| Tg(foxj1a:GCaMP7f) | Motile ciliated cells (ependymal cells) | Green calcium indicator | generated | Danio rerio | Zhang et al. (2020) \| Dana et al. (2019) |
| Tg(elavl3:gtACR2-eYFP) | Panneuronal | Inhibitory opsin & yellow fluorescent marker | generated | Danio rerio | Park et al. (2000) \| Mohammad et al. (2017) |
| Tg(β-actin2:mCherry-CAAX; myl7:GFP) | Pancellular | Red membrane marker & Green cardiac marke | generated | Danio rerio | Fowler et al. (2016) |
| Tg(VAChTa:Gal4); Tg(UAS:CoChR-eGFP)jf4 | Cholinergic neurons (motor neurons) | Excitatory opsin & green fluorescent marker | | Danio rerio | Taniguchi et al. (2017) \| Mu et al. (2019) |
| Tg(th:Gal4); Tg(UAS:GCaMP6f)jf46 | Catecholaminergic neurons (sympathetic gangl | Green calcium indicator | | Danio rerio | Li et al. (2015) \| Mu et al. (2019) |
| Tg(flk1:dsRED-CAAX) | Vascular endothelial cells | Red membrane marker | | Danio rerio | Fujita et al. (2011) |
| Tg(gata1:dsRED) | Erythrocytes | Red fluorescent marker | | Danio rerio | Traver et al. (2003) |
| Tg(isl1CREST-hsp70l:mRFP) | Motor neurons | Red fluorescent marker | | Danio rerio | Grant et al. (2010) |
| Tg(phox2bb:eGFP) | Autonomic nervous system | Green fluorescent marker | | Danio rerio | Nechiporuk et al. (2007) |
| Tg(foxj1a:eGFP) | Motile ciliated cells (ependymal cells) | Green fluorescent marker | | Danio rerio | Grimes et al. (2016) |
| Tg(gfap:jRGECO1b) | Glia | Red calcium indicator | | Danio rerio | Mu et al. (2019) |
| Tg(elavl3:H2B-jRGECO1a) | Neuronal | Nuclear localized red calcium inidcator | | Danio rerio | Yang et al. (2022) |
| Tg(ubbR:jGCaMP8m) | Pancellular | Green calcium indicator | generated | Danionella cerebrum | Bakunaite et al. (2024) \| Zhang et al. (2023) |
| TgBAC(lamC1:lamC1-sfGFP) | Basement membrane | Green fluorescent marker | | Danio rerio | Yamaguchi et al. (2022) |
| mitfa(-/-) | Melanocytes | Missense mutant | generated | Danionella cerebrum | Schulze et al. (2018) |

#### Example of known roles of calcium in various cell types

| Cell type | Associated Organ/System | Example of role of calcium in cell type | Reference | Reference URL |
|---|---|---|---|---|
| Neuron | Nervous system (CNS/PNS) | $Ca^{2+}$ influx through voltage-gated CaV channels couples electrical activity to neurotransmitter release by binding synaptotagmins to trigger synaptic vesicle fusion; it also regulates synaptic plasticity and gene transcription. | Sudhof TC. Calcium control of neurotransmitter release. Cold Spring Harbor Perspectives in Biology. 2012 Jan;4(1):a011353. https://doi.org/10.1101/cshperspect.a011353 | https://cshperspectives.cshlp.org/content/4/1/a011353.full |
| Astrocyte | Nervous system (CNS) | $Ca^{2+}$ elevations in astrocytes mediate ATP release. | Khakh, B.S. & McCarthy, K.D. (2015). Astrocyte calcium signaling: from observations to functions and the challenges therein. Cold Spring Harb Perspect Biol. 7(4):a020404. | https://pubmed.ncbi.nlm.nih.gov/25605709/ |
| Microglia | Nervous system (CNS) | $Ca^{2+}$ transients control motility and release of cytokines. | Kettenmann, H. et al. (2011). Physiology of microglia. Physiol Rev. 91(2):461-553. | https://pubmed.ncbi.nlm.nih.gov/21527731/ |
| Radial glial cell | Nervous system (CNS) | $Ca^{2+}$ dynamics regulate neural stem cell proliferation/differentiation. $Ca^{2+}$ oscillations influence cell-cycle progression. | Weissman, T.A. et al. (2004). Calcium waves propagate through radial glial cells and modulate proliferation in the developing neocortex. Neuron. 43 (5):647-61. | https://pubmed.ncbi.nlm.nih.gov/15339647/ |
| Oligodendrocyte | Nervous system (CNS) | $Ca^{2+}$ transients control myelin sheath growth. Activity-dependent $Ca^{2+}$ in sheaths promotes actin remodeling and elongation. | Oligodendrocyte calcium signaling promotes actin-dependent myelin sheath extension | https://pubmed.ncbi.nlm.nih.gov/38177161/ |
| Photoreceptors | Eye (retina) | $Ca^{2+}$ controls phototransduction adaptation and synaptic release. | Pugh, E.N. Jr. & Lamb, T.D. (2000). Phototransduction in vertebrate rods and cones: molecular mechanisms of amplification, recovery and light adaptation. Handb Biol Phys. 3:183-255. | https://www.sciencedirect.com/science/article/pii/S1383812100800081 |
| Inner ear hair cell | Ear (auditory/vestibular) | Mechanotransduction induces $Ca^{2+}$ entry and contributes to synaptic release. | Fuchs, P.A. (2005). Time and intensity coding at the hair cell's ribbon synapse. J Physiol. 566(Pt 1):7-12. | |
| Neuromast hair cell | Lateral line | $Ca^{2+}$ involved in mechanotransduction and synaptic release. | Synaptically silent sensory hair cells in zebrafish are recruited after damage | https://www.nature.com/articles/s41467-018-03806-8 |
| Endothelial cell | Vasculature | $Ca^{2+}$ controls permeability and angiogenesis. | Endothelial Cell Calcium Signaling during Barrier Function and Inflammation | https://pmc.ncbi.nlm.nih.gov/articles/PMC7074364/ |
| Cardiomyocyte | Heart | Excitation-contraction coupling via $Ca^{2+}$-induced $Ca^{2+}$ release. L-type $Ca^{2+}$ influx triggers RyR-mediated SR $Ca^{2+}$ release, activating troponin C. | Bers, D.M. (2002). Cardiac excitation-contraction coupling. Nature. 415(6868):198-205. | https://pubmed.ncbi.nlm.nih.gov/11805843/ |
| Mural cell (pericyte/smooth muscle) | Vasculature | Pericyte/smooth muscle contraction depends on $Ca^{2+}$-calmodulin-MLCK. Vasoactive signals elevate $Ca^{2+}$ to control tone and capillary diameter. | Hill-Eubanks, D.C. et al. (2011). Calcium signaling in smooth muscle. Cold Spring Harb Perspect Biol. 3(9):a004549. | https://pubmed.ncbi.nlm.nih.gov/21709182/ |
| Erythrocyte (red blood cell) | Blood | $Ca^{2+}$ is involved in regulating volume and intercellular adhesion. | Stimulation of human red blood cells leads to Ca2+-mediated intercellular adhesion | https://pubmed.ncbi.nlm.nih.gov/21616535/ |
| Thrombocyte (platelet-equivalent) | Blood | Platelet activation and secretion require $Ca^{2+}$. $Ca^{2+}$ release/influx drives shape change, integrin activation, granule exocytosis. | Varga-Szabo, D. et al. (2009). Calcium signaling in platelets. J Thromb Haemost. 7(7):1057-66. | https://pubmed.ncbi.nlm.nih.gov/19422456/ |
| Skeletal muscle fiber | Muscle (somitic) | Depolarization triggers SR $Ca^{2+}$ release and contraction. DHPR-RyR coupling releases $Ca^{2+}$; troponin C binding initiates cross-bridge cycling. | Melzer, W. et al. (1995). The role of $Ca^{2+}$ ions in excitation-contraction coupling of skeletal muscle fibres. Biochim Biophys Acta. 1241(1):59-116. | https://pubmed.ncbi.nlm.nih.gov/7742348/ |
| Smooth muscle cell | Muscle (visceral/vascular) | $Ca^{2+}$-calmodulin-MLCK pathway drives contraction. | Nelson et al. (2011). Calcium Signaling in Smooth Muscle. Cold Spring Harbor Perspectives in Biology. | https://pmc.ncbi.nlm.nih.gov/articles/PMC3181028/ |
| Chondrocyte | Cartilage | $Ca^{2+}$ mediates mechanotransduction and response to osmotic stress | Li W, Zhou Y, Han L, Wang L, Lucas Lu X. Calcium signaling of primary chondrocytes and ATDC5 chondrogenic cells under osmotic stress and mechanical stimulation. J Biomech. 2022 Dec;145:111388. doi: 10.1016/j.jbiomech.2022.111388. Epub 2022 Nov 17. PMID: 36413831; PMCID: PMC10472919. | https://pmc.ncbi.nlm.nih.gov/articles/PMC10472919/ |
| Osteoblast | Bone | Regulates osteoblast differentiation. | Hao Y, Yang N, Sun M, Yang S, Chen X. The role of calcium channels in osteoporosis and their therapeutic potential. Front Endocrinol (Lausanne). 2024 Aug 7;15:1450328. doi: 10.3389/fendo.2024.1450328. PMID: 39170742; PMCID: PMC11335502. | https://pmc.ncbi.nlm.nih.gov/articles/PMC11335502/ |
| Osteocyte | Bone | $Ca^{2+}$ transients encode mechanical load, induce vesicular release and induce actin contraction. | Morrell, A.E., Brown, G.N., Robinson, S.T. et al. Mechanically induced Ca2+ oscillations in osteocytes release extracellular vesicles and enhance bone formation. Bone Res 6, 6 (2018). https://doi.org/10.1038/s41413-018-0007-x | https://www.nature.com/articles/s41413-018-0007-x |
| Osteoclast | Bone | $Ca^{2+}$ regulates cytoskelatal organization, elevated calcium inhibits resorbing activity. | Li, Zhenpeng, Kangmei Kong, and Weili Qi. "Osteoclast and its roles in calcium metabolism and bone development and remodeling." Biochemical and biophysical research communications 343.2 (2006): 345-350. | https://www.sciencedirect.com/science/article/pii/S0006291X06004608#section.0070 |
| Notochord cell | Axial skeleton (embryonic) | $Ca^{2+}$ contributes to cell elongation and intercalation. | Mapping calcium dynamics in a developing tubular structure<br>Jorgen Hoyer, Morsal Saba, Daniel Dondorp, Kushal Kolar, Riccardo Esposito, Marios Chatzigeorgiou<br>bioRxiv 2020.10.16.342535; doi: https://doi.org/10.1101/2020.10.16.342535 | https://www.biorxiv.org/content/10.1101/2020.10.16.342535v1.full |
| Goblet cell | Intestine | $Ca^{2+}$ triggers mucin granule exocytosis. | Vivek Malhotra (2013) TRPM5-mediated calcium uptake regulates mucin secretion from human colon goblet cells eLife 2:e00658. | https://elifesciences.org/articles/00658#content |
| Hepatocyte | Liver | Hormone-evoked $Ca^{2+}$ waves induce glucose mobilisation through regulation of gluocogenolysis and neoglucogenesis. | Humbert, Alexandre, et al. "Calcium signalling in hepatic metabolism: Health and diseases." Cell Calcium 114 (2023): 102780. | https://www.sciencedirect.com/science/article/pii/S0143416023000921#sec0002 |
| Pancreatic alpha cell | Pancreas (endocrine) | Voltage-gated $Ca^{2+}$ drives glucagon release at low glucose. | Rorsman P, Braun M, Zhang Q. Regulation of calcium in pancreatic α- and β-cells in health and disease. Cell Calcium. 2012 Mar-Apr;51(3-4):300-8. doi: 10.1016/j.ceca.2011.11.006. Epub 2011 Dec 15. PMID: 22177710; PMCID: PMC3334273. | https://pmc.ncbi.nlm.nih.gov/articles/PMC3334273/ |
| Pancreatic beta cell | Pancreas (endocrine) | $Ca^{2+}$ influx directly triggers insulin exocytosis. KATP closure→depolarization→Cav channels→$Ca^{2+}$-triggered granule fusion. | Rorsman, P. & Ashcroft, F.M. (2018). Pancreatic β-cell electrical activity and insulin secretion: of mice and men. Physiol Rev. 98(1):117-214. | https://pubmed.ncbi.nlm.nih.gov/29212789/ |
| Juxtaglomerular cells | Kidney (glomerulus) | $Ca^{2+}$ inhibits renin secretion. | Staruschenko A, Alexander RT, Caplan MJ, Ilatovskaya DV. Calcium signalling and transport in the kidney. Nat Rev Nephrol. 2024 Aug;20(8):541-555. doi: 10.1038/s41581-024-00835-z. Epub 2024 Apr 19. PMID: 38641658; PMCID: PMC12036682. | https://pmc.ncbi.nlm.nih.gov/articles/PMC12036682/#S15 |
| Chromaffin cell | Head kidney (adrenal medulla homolog) | Depolarization-triggered $Ca^{2+}$ induces catecholamine secretion. | García, Antonio G., et al. "Calcium signaling and exocytosis in adrenal chromaffin cells." Physiological reviews 86.4 (2006): 1093-1131 | https://journals.physiology.org/doi/full/10.1152/physrev.00039.2005 |
| Interrenal steroidogenic cell | Head kidney (adrenal cortex homolog) | Sustained $Ca^{2+}$ influx increases steroidogenesis. | Rossier, M. F. "T-type calcium channel: a privileged gate for calcium entry and control of adrenal steroidogenesis. Front Endocrinol. 2016; 7: 43." | https://pmc.ncbi.nlm.nih.gov/articles/PMC4873500/ |
| Pituitary corticotroph (ACTH) | Pituitary | CRH stimulates $Ca^{2+}$ signaling to release ACTH. cAMP-PKA primes secretion; $Ca^{2+}$ is final trigger for granule fusion. | Tse, Amy, Andy K. Lee, and W. Tse Frederick. "Ca2+ signaling and exocytosis in pituitary corticotropes." Cell Calcium 51.3-4 (2012): 253-259 | https://www.sciencedirect.com/science/article/pii/S0143416011002363 |
| Kupffer cell (liver macrophage) | Liver | $Ca^{2+}$ contributes to Kupffer cell M2 polarization. | Xu, Xue-song, et al. "SCARF1 promotes M2 polarization of Kupffer cells via calcium-dependent PI3K-AKT-STAT3 signalling to improve liver transplantation." Cell proliferation 54.4 (2021): e13022. | https://onlinelibrary.wiley.com/doi/10.1111/cpr.13022 |
| Ultimobranchial (calcitonin) cell | Ultimobranchial gland | Systemic $Ca^{2+}$ regulates calcitonin secretion. High plasma $Ca^{2+}$ stimulates calcitonin release to lower $Ca^{2+}$; $Ca^{2+}$ acts as both signal and target. | Fudge, N.J., Kovacs, C.S. Physiological studies in heterozygous calcium sensing receptor (CaSR) gene-ablated mice confirm that the CaSR regulates calcitonin release in vivo . BMC Physiol 4, 5 (2004). https://doi.org/10.1186/1472-6793-4-5 | https://bmcphysiol.biomedcentral.com/articles/10.1186/1472-6793-4-5#:~:text=The%20calcium%20sensing%20receptor%20(CaSR,responsiveness%20of%20calcitonin%20to%20calcium. |
| Skin epithelial cells | Skin | Ca2+ signaling drives cell cycle progression | Moore, Jessica L., et al. "Cell cycle controls long-range calcium signaling in the regenerating epidermis." Journal of Cell Biology 222.7 (2023): e202302 | https://rupress.org/jcb/article/222/7/e202302095/214066/Cell-cycle-controls-long-range-calcium-signaling |
| Enteroendocrine cell | Intestine | Stimulus-secretion coupling via $Ca^{2+}$. Nutrient cues evoke $Ca^{2+}$ to release peptides (e.g., 5-HT, CCK); | Davison, Adam, Frank Reimann, and Fiona M. Gribble. "Molecular mechanisms of stimulus detection and secretion in enteroendocrine cells." Current O | https://www.sciencedirect.com/science/article/pii/S0959438825000765#: |
| Monocyte/Macrophage | Immune (innate) | Purinergic receptors is strongly coupled to Ca2+-signaling; Ca2+ involved in phagosome maturation, engulfment of apoptotic cells, migration. | Desai, Bimal N., and Norbert Leitinger. "Purinergic and calcium signaling in macrophage function and plasticity." Frontiers in immunology 5 (2014): 580 | https://pmc.ncbi.nlm.nih.gov/articles/PMC4245916/ |

#### Limitations

We foresee multiple extensions of WHOLISTIC which would extended the scope of this approach. In this instantiation, a) samples are embedded in agarose which limits their movement; moreover, to best image the viscera, b) fish are placed on their sides which is not a natural posture; furthermore, spinning-disk confocal microscopy was used which induces c) more bleaching than light-sheet microscopy or 2-photon microscopy and is relatively slower. Finally, d) with the main objective used in this study, 20x 0.75NA, the field of view was restricted to 800 µm ×800 µm. By using a large field of view 2-photon microscope[21] and positioning a 45º mirror next to the fish it would be possible to image the entire length of the fish, positioned in its natural posture, and access both the brain and viscera (via the mirror) simultaneously. Such a microscope also includes a resonant scanner and offers the capacity to do dual plane imaging which would double the frame rate. Similar extensions could be made to freely swimming imaging setups[?] to enable whole body imaging in freely behaving animals. Furthermore, GCaMP has primarily been optimized for imaging neuronal calcium dynamics, and the dynamic range of the sensor might not be optimal for all other cell types. Whilst we did not observe saturation of variance in cells with high baseline levels, this does not rule out sensor saturation. Quantitative calcium measurements using fluorescence life-time imaging would be the definitive way of assessing the dynamic range of calcium concentrations across cells of the body.

[4] Jinxing Lin, Qiusheng Chen, and Jianhua Hu. *Color atlas of zebrafish histology and cytology.* Springer Nature, 2022.

[5] David Grant Colburn Hildebrand, Marcelo Cicconet, Russel Miguel Torres, Woohyuk Choi, Tran Minh Quan, Jungmin Moon, Arthur Willis Wetzel, Andrew Scott Champion, Brett Jesse Graham, Owen Randlett, George Scott Plummer, Ruben Portugues, Isaac Henry Bianco, Stephan Saalfeld, Alexander David Baden, Kunal Lillaney, Randal Burns, Joshua Tzvi Vogelstein, Alexander Franz Schier, Wei Chung Allen Lee, Won Ki Jeong, Jeff William Lichtman, and Florian Engert. Whole-brain serial-section electron microscopy in larval zebrafish. *Nature*, 545(7654):345–349, 2017. ISSN 14764687. doi: 10.1038/nature22356. URL `http://dx.doi.org/10.1038/nature22356`.

[6] B. Thisse, V. Heyer, A. Lux, V. Alunni, A. Degrave, I. Seiliez, J. Kirchner, JP. Parkhill, and C. Thisse. Spatial and temporal expression of the zebrafish genome by large-scale in situ hybridization screening. *Methods Cell Biol*, 2004. doi: 10.1016/s0091-679x(04)77027-2.

[7] Ulrike Von Luxburg. A tutorial on spectral clustering. *Statistics and Computing*, 17(4): 395–416, 2007. ISSN 09603174. doi: 10.1007/s11222-007-9033-z.

[8] NMD Niezink. *Consensus in networked multi-agent systems.* PhD thesis, Faculty of Science and Engineering, 2011.
